# Repurposing of Synaptonemal Complex Proteins for Kinetochores in Kinetoplastida

**DOI:** 10.1101/2021.02.06.430040

**Authors:** Eelco C. Tromer, Thomas A. Wemyss, Ross F. Waller, Bungo Akiyoshi

## Abstract

Chromosome segregation in eukaryotes is driven by a macromolecular protein complex called the kinetochore that connects centromeric DNA to microtubules of the spindle apparatus. Kinetochores in well-studied model eukaryotes consist of a core set of proteins that are broadly conserved among distant eukaryotic phyla. In contrast, unicellular flagellates of the class Kinetoplastida have a unique set of kinetochore components. The evolutionary origin and history of these kinetochores remains unknown. Here, we report evidence of homology between three kinetoplastid kinetochore proteins KKT16–18 and axial element components of the synaptonemal complex, such as the SYCP2:SYCP3 multimers found in vertebrates. The synaptonemal complex is a zipper-like structure that assembles between homologous chromosomes during meiosis to promote recombination. Using a sensitive homology detection protocol, we identify divergent orthologues of SYCP2:SYCP3 in most eukaryotic supergroups including other experimentally established axial element components, such as Red1 and Rec10 in budding and fission yeast, and the ASY3:ASY4 multimers in land plants. These searches also identify KKT16–18 as part of this rapidly evolving protein family. The widespread presence of the SYCP^2-3^ gene family in extant eukaryotes suggests that the synaptonemal complex was likely present in the last eukaryotic common ancestor. We found at least twelve independent duplications of the SYCP^2-3^ gene family throughout the eukaryotic tree of life, providing opportunities for new functional complexes to arise, including KKT16–18 in *Trypanosoma brucei*. We propose that kinetoplastids evolved their unique kinetochore system by repurposing meiotic components of the chromosome synapsis and homologous recombination machinery that were already present in early eukaryotes.

## Introduction

Chromosome segregation in eukaryotes is driven by spindle microtubules and kinetochores. Microtubules are dynamic polymers that consist of α-/β-tubulin subunits, while the kinetochore is the macromolecular protein complex that assembles onto centromeric DNA and interacts with spindle microtubules during mitosis and meiosis [1]. All studied eukaryotes use spindle microtubules to drive chromosome movement, and α-/β-tubulins are amongst the most conserved proteins in eukaryotes [2]. The kinetochore is a highly complicated structure that consists of more than 30 unique structural proteins even in a relatively simple budding yeast kinetochore [3]. CENP-A is a centromere-specific histone H3 variant that specifies kinetochore assembly sites, while the NDC80 complex (NDC80, NUF2, SPC24, SPC25) constitutes the primary microtubule-binding activity of kinetochores [4]. Functional studies have established that CENP-A and the NDC80 complex are essential for kinetochore function in several model eukaryotes (e.g. yeasts, worms, flies, and humans) [3]. However, CENP-A is absent in some eukaryotic lineages such as holocentric insects [5], early-diverging Fungi [6], and Kinetoplastida [7]. Furthermore, it is known that compositions of kinetochores can vary considerably among eukaryotes [8], and that these components are highly divergent at the sequence level [8–10]. The most extreme case known to date is found in Kinetoplastida, for which no apparent direct homologues of the canonical kinetochore proteins were detected [11].

Kinetoplastida comprises unicellular flagellates characterised by the presence of a conspicuous mitochondrial structure called the kinetoplast that contains a unique form of mitochondrial DNA [12]. This group belongs to the phylum Euglenozoa and is evolutionarily divergent from many popular fungal and animal model eukaryotes (all belonging to Opisthokonta) [13]. There are several medically important kinetoplastid parasites such as *Trypanosoma brucei, Trypanosoma cruzi*, and *Leishmania* spp., which cause neglected tropical diseases [14]. Although chromosome segregation depends on spindle microtubules in *T. brucei* [15], we previously identified 24 unique kinetoplastid kinetochore proteins (KKT1–20 and KKT22–25) that localise at centromeres in this species, which together constitute a functionally analogous kinetochore structure [16–18]. Furthermore, 12 additional proteins, named KKT-interacting proteins (KKIP1–12), have been identified that associate with kinetochores during mitosis [19,20]. None of these 36 kinetoplastid kinetochore subunits appear to be orthologous to canonical kinetochore proteins, and the evolutionary relationship between the two kinetochore systems remains unclear [21]. Interestingly, while non-kinetoplastid species have significant distance in-between sister kinetochores in metaphase cells (the space is called the inner centromere) [22–24], there is no clear separation between sister kinetochores in all studied kinetoplastids [25–28]. This structural difference is consistent with compositional differences between canonical and kinetoplastid kinetochores.

Tromer and van Hooff *et al*. traced the origin of canonical kinetochore subunits back to before the last eukaryotic common ancestor (LECA), using a combination of phylogenetic trees, profile-versus-profile homology detection and structural comparisons of its protein components [29]. They found that duplications played a major role in shaping the ancestral eukaryotic kinetochore, and that its components share a deep evolutionary history with proteins of various other prokaryotic and eukaryotic pathways e.g. ubiquitination, DNA damage repair and the flagellum [29]. In these analyses, none of the 36 KKT/KKIP proteins of the kinetoplastid kinetochore were considered because they were found only in Kinetoplastida and were therefore deemed not to have been part of the kinetochore in LECA. *If KKT/KKIP proteins were not part of the kinetochore in ancestral eukaryotes, when and from where did these new components of the kinetochore originate?* (i) They may have a mosaic origin similar to the LECA kinetochore [29], and might have been pieced together and co-opted from other processes, (ii) they may have arisen from external sources (e.g. viral integrations into the genome, bacterial endosymbionts), or (iii) they may have arisen through a combination of genomic translocations, fusion of existing genes and de novo gene birth in the first ancestors of Kinetoplastida. (iv) An alternative hypothesis is that KKT/KKIP subunits might be canonical kinetochore proteins that diverged to such an extent that they cannot be readily identified through sequence searches. Indeed, a recent study found intriguing similarities between the outer kinetochore protein KKIP1 and the coiled coils, but not the microtubule-binding Calponin Homology (CH) domains, of two NDC80 complex members (NDC80 and NUF2) [19]. In addition, Aurora kinase and INCENP, two members of the chromosome passenger complex that localise at the inner centromere region in other eukaryotes, are found in kinetoplastids [30], suggesting that some processes or components of canonical chromosome segregation systems are conserved in kinetoplastids as well. Detailed phylogenetic analyses and sensitive homology searches of KKT/KKIP proteins will need to be performed to evaluate which of the above evolutionary scenarios applies to the kinetoplastid kinetochore.

Previous studies identified several domains in KKT/KKIP proteins that are commonly found among both eukaryotic and prokaryotic proteins, providing some clues to the functionality and evolutionary history of the proteins that make up the kinetoplastid kinetochore. These include a BRCA1 carboxy-terminal (BRCT) domain in KKT4, a forkhead-associated (FHA) domain in KKT13, a 7-bladed WD40 β-propeller in KKT15, a Gcn5-related N-acetyltransferase (GNAT) domain in KKT23, a protein kinase domain of unknown affiliation in KKT2 and KKT3, a Cdc2-like kinase (CLK) domain in KKT10 and KKT19, and large coiled coil regions (e.g. KKIP1 and KKT24) [16–19]. BRCT and FHA domains are frequently found in proteins involved in the DNA damage response [31], which is important not only for DNA damage repair in somatic cells but also for homologous recombination in meiotic cells [32]. Furthermore, it has been proposed, based on the similarity in the C-terminal polo boxes, that KKT2,3 and 20 share common ancestry with Polo-like kinases (PLKs), which regulate various biological functions such as the cell cycle progression, the DNA damage response, kinetochores, centrosomes, and synaptonemal complexes [31,33]. PLKs localise at kinetochores in some organisms but are not thought to be a structural kinetochore component [34,35]. Although the kinase domains of KKT2,3 have been classified as unique among eukaryotic kinase subfamilies [36], they have some similarity to the kinase domain of PLKs [21]. KKT2 and KKT3 also have a unique zinc-binding domain in the central region, which promotes their centromere localisation [37]. Gene duplication plays an important role in generating new functions using pre-existing proteins [38], and this process has further contributed to kinetoplastid kinetochore development. The paralogues KKT2,3 and 20, and KKT10,19 have arisen through gene duplications, and this also appears to be the case for KKT17 and KKT18 [16]. Although KKT17,18 are conserved even in divergent kinetoplastids [39], previous studies did not reveal any recognisable domains in these proteins [16].

While the identification of conserved domains in kinetoplastid kinetochore proteins offers some insight into the molecular mechanisms of function for these proteins, there has previously been no direct link to any pre-existing machinery from which it might be derived. In this study, we report evidence of homology between the KKT16–18 complex and the axial elements of the synaptonemal complex, a meiosis-specific tripartite structure that assembles in between homologous chromosomes [40]. We speculate on the implications of this finding for the origin of kinetoplastid kinetochores.

## Results

### KKT16, KKT17, and KKT18 form the KKT16 complex

Our previous immunoprecipitation and mass spectrometry of KKT proteins suggested that KKT16, KKT17, and KKT18 likely form a subcomplex [16]. Consistent with this possibility, these proteins have a similar localisation pattern during the cell cycle, showing diffuse nuclear signals in G1 and forming kinetochore dots from S phase to anaphase in *T. brucei* [16]. KKT17 and KKT18 have a moderate degree of sequence identity and similarity (23% shared identity and 38% similarity for *T. brucei* sequences), suggesting that these two proteins are the product of a duplication event [16]. Indeed, their predicted secondary structure revealed a highly similar topology, with an N-terminal globular region consisting of repeated alpha helices, followed by a beta sheet-rich domain, and a C-terminal coiled coil, connected by a disordered linker region (Figure 1a). KKT16 consists of a single coiled-coil region, signifying the coiled coils of KKT16, 17 and 18 as a potential basis for their association (Figure 1a). To test whether KKT16–18 interact with each other, we expressed these three proteins in bacteria using a polycistronic expression system [41] (Figure 1b). We found that KKT17 and KKT18 co-purified with 6xHis-KKT16 (Figure 1c), showing that these three proteins on their own are sufficient to form a complex, which we refer to as the KKT16 complex.

**Figure 1.**
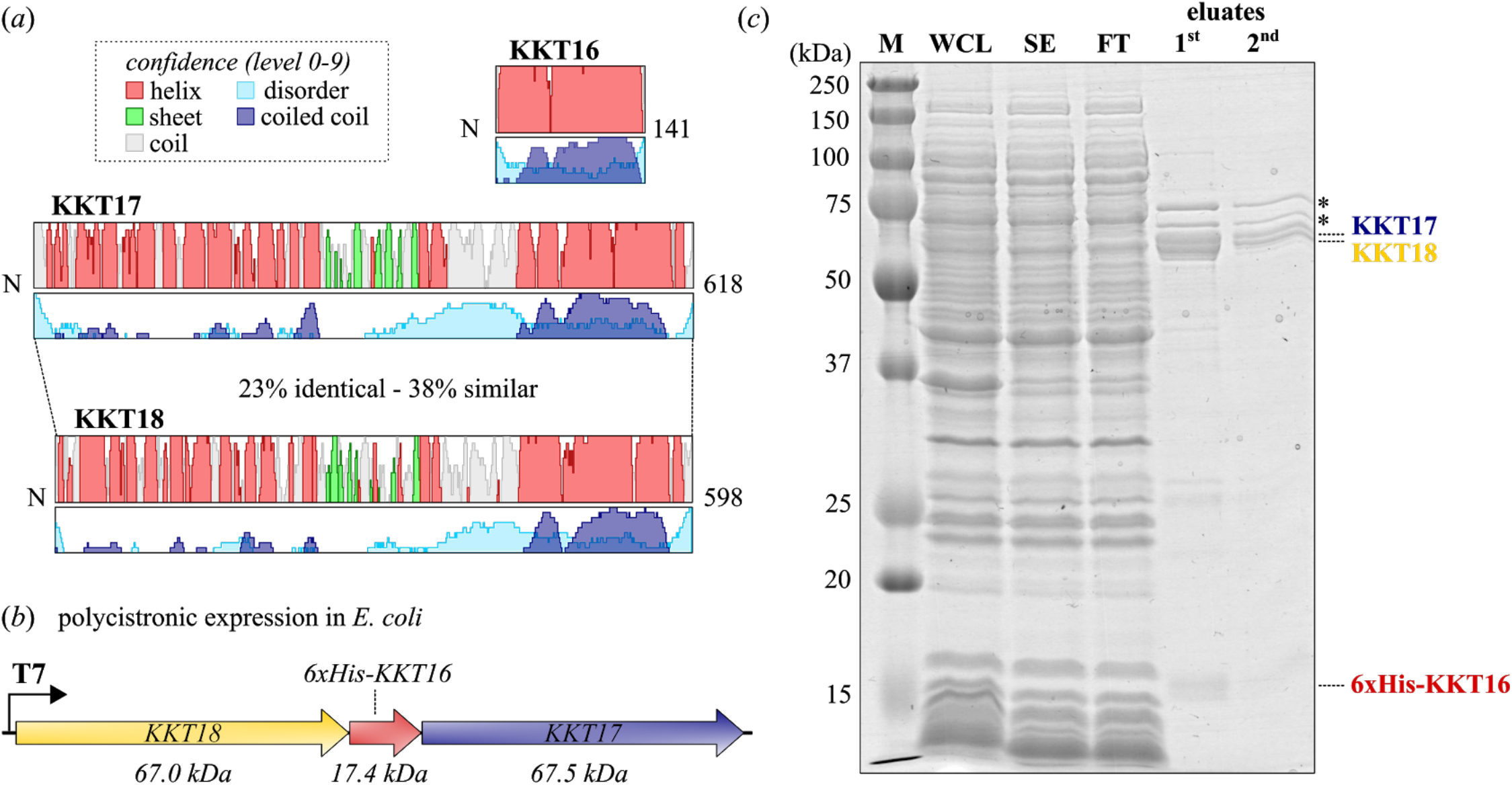
KKT16, KKT17, and KKT18 form the KKT16 complex. (A) Cartoon of the KKT16, KKT17, and KKT18 proteins, including secondary structure predictions based on multiple sequence alignments of kinetoplastid homologues with *T. brucei* KKT proteins as a query. Height of each track indicates the confidence of each prediction (Data & Methods). Confidence levels are discretised into ten levels (0-9). Identity percentages for *T. brucei* KKT17 and KK18 were calculated using the Needleman-Wunsch global alignment algorithm, with the BLOSUM62 matrix to derive their similarity score [87]. (B) Schematic of the construct used to co-express the three subunits. See Figure S1 for expression tests. (C) Expression and purification of the KKT16 complex from *E. coli*. Coomassie-stained 12% acrylamide gel of the purification is shown. Asterisks (**\***) indicate common contaminants; M: marker; WCL: whole cell lysate; SE: soluble extract; FT: flow through.

### KKT16–18 are similar to axial element components of the synaptonemal complex

To identify potential homologues of the KKT16 complex proteins and to detect proteins or protein complexes with similar domain architectures, we used Hidden Markov Modelling methods (Data & Methods). We generated multiple sequence alignments (MSAs) of previously detected homologues of KKT16–18 found among divergent kinetoplastids [16,39]. We then constructed profile Hidden Markov Models (HMM) based on these MSAs and performed secondary structure-aware profile-versus-profile HMM searches against databases of known conserved domains and structures (Data & Methods, Figure 2a, Figure 3a) using HHsearch [42]. Because KKT17 and KKT18 appeared to be paralogues, we included all kinetoplastid homologues of these proteins into a single MSA, aiming to increase the similarity detection sensitivity. Using this approach, we found that KKT17 and KKT18 consist of three domains frequently found in eukaryotic proteins (Figure 2a, see File S1 for output files): (i) an N-terminal armadillo repeat region (ARM, many high probability homologues: Prob>80%, E-value~1), followed by (ii) a β-barrel Pleckstrin Homology (PH) domain found in chromatin (e.g. histone chaperone Rtt106, PfamA: PF08512, Prob: 75%, E~10) and membrane-associated proteins (e.g. ESCRT complex protein Vps36, PfamA: PF11605, Prob: 84%, E~10), which are linked by a structurally disordered region to (iii) a C-terminal coiled coil that has similarity to many higher order-forming coiled-coil proteins, with the extracellular lipid-binding apolipoproteins as strongest scoring profile HMM (APO, PfamA: PF01442, Prob:97%, E~10^-1^). Searches with the profile HMM of KKT16 revealed similarity with a large number of coiled-coil components of various eukaryotic protein complexes (Figure 3a, see File S1 for raw output files), similar to KKT17 and KKT18, and with apolipoprotein as the highest scoring HMM (APOE, pdb: 2L7B, Prob: 95%, E~1).

**Figure 2.**
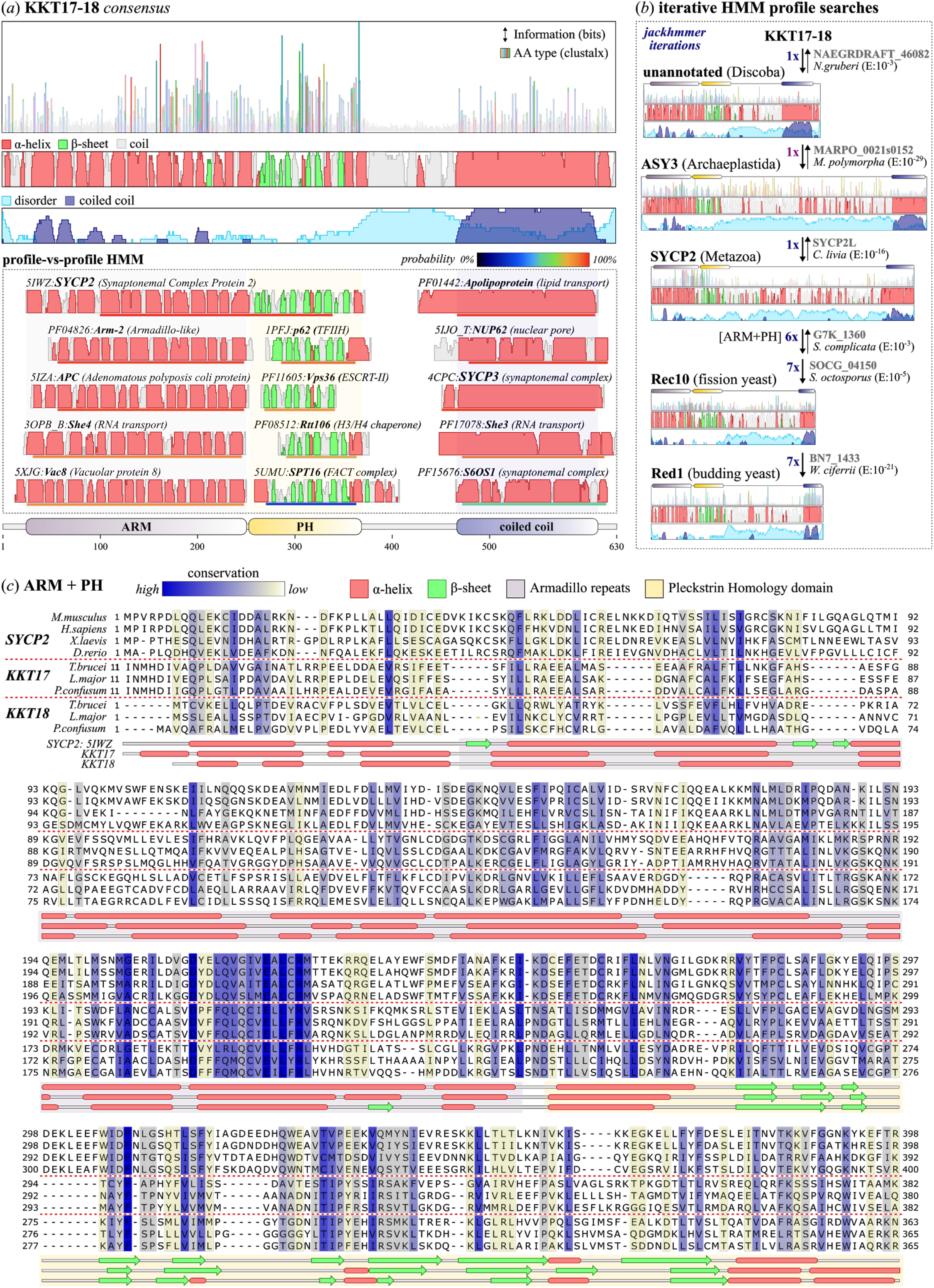
KKT17,18 are orthologues of the eukaryotic SC proteins SYCP2, ASY3, Red1 and Rec10. (A) Cartoon of the conserved domain architecture and sequence features of a consensus of homologues of KKT17 and KKT18 (see File S3 for MSA, File S6 for details of the sequence annotation). From top to bottom: (1) collapsed sequence logo with different conserved amino acids for each position represented by the ClustalX colouring scheme. Height indicates the bits of information per position and is used as a proxy for conservation of amino acids at given positions; (2) secondary structure prediction, similar to Figure 1. Height indicates the probability of the prediction for each position (level 0-9); (3) overview of selected results (best scoring hits - top, generic hits - towards the bottom) from a profile-vs-profile HMM search (HHsearch, see Data & Methods) of KKT17,18 against the PFAM (conserved domains) and PDB (3D structures) database. Identifiers starting with ‘PF’ indicate PFAM domains and 4 digit identifiers (e.g. 5IWZ) indicate PDB entries. Terms between parentheses indicate relevant functional or domain annotation. For each hit only a proportion of the total protein/domain is shown around the region that has a significant similarity with the KKT17,18 HMM. Coloured bars below each of the proteins/domains indicate the HHsearch probability score; (4) topology diagram of conserved domains in KKT17 and KKT18; ARM: armadillo repeats; PH: pleckstrin homology domain. (B) Cartoon of the similarity detection protocol (Data & Methods) for establishing homology between highly divergent synaptonemal complex proteins KKT17,18, SYCP2, ASY3, Red1 and Rec10. Similarity detection does not necessarily have to be performed in this order, but has been visualised in a linear manner to showcase the searchpath from KKT17,18 to other eukaryotic SYCP2-type proteins. The seed HMM for KKT17,18 is based on the same multiple sequence alignment as shown in panel A. Thick arrows and E-values indicate the direction and significance of the similarity searches. Thin arrows in the reverse direction indicate that reciprocal searches yield similar homologous connections. Dark purple numbers indicate the number of iterations of HMM searches needed to include a particular sequence. Uniprot identifiers (grey) on the right indicate the highest scoring orthologues after each iteration belonging to a larger group of related sequences (e.g. SYCP2-type proteins in Metazoa) for which separate conservation and architecture cartoons are presented as in panel A (see File S3 for cartoons of all SYCP2-type HMMs). The positions of the ARM and PH domain, and coiled coils are projected on top of the cartoons. Brackets indicate when only selected domains are used for similarity detection. Full species names: *Naegleria gruberi; Marchantia polymorpha; Columba livia; Saitoella complicata; Schizosaccharomyces octosporus; Wickerhamomyces ciferrii*. (C) Multiple sequence alignment of the N-terminal ARM repeats and PH domain of SYCP2, KKT17 and KKT18 homologues (see for colour coding settings jalview session files File S5). Columns with single amino acid occupancy were removed. Secondary structure prediction by PSIPRED based on a multiple sequence alignment of KKT17 and KKT18 using the *T. brucei* proteins as seed sequence (Data & Methods). The secondary structure of SYCP2 is mapped based on the PDB file 5IWZ (mouse SYCP2) [43]. Full species names: *Mus musculus; Homo sapiens;Xenopus laevis; Danio rerio; Trypanosoma brucei; Leishmania major; Paratrypanosoma confusum*.

**Figure 3.**
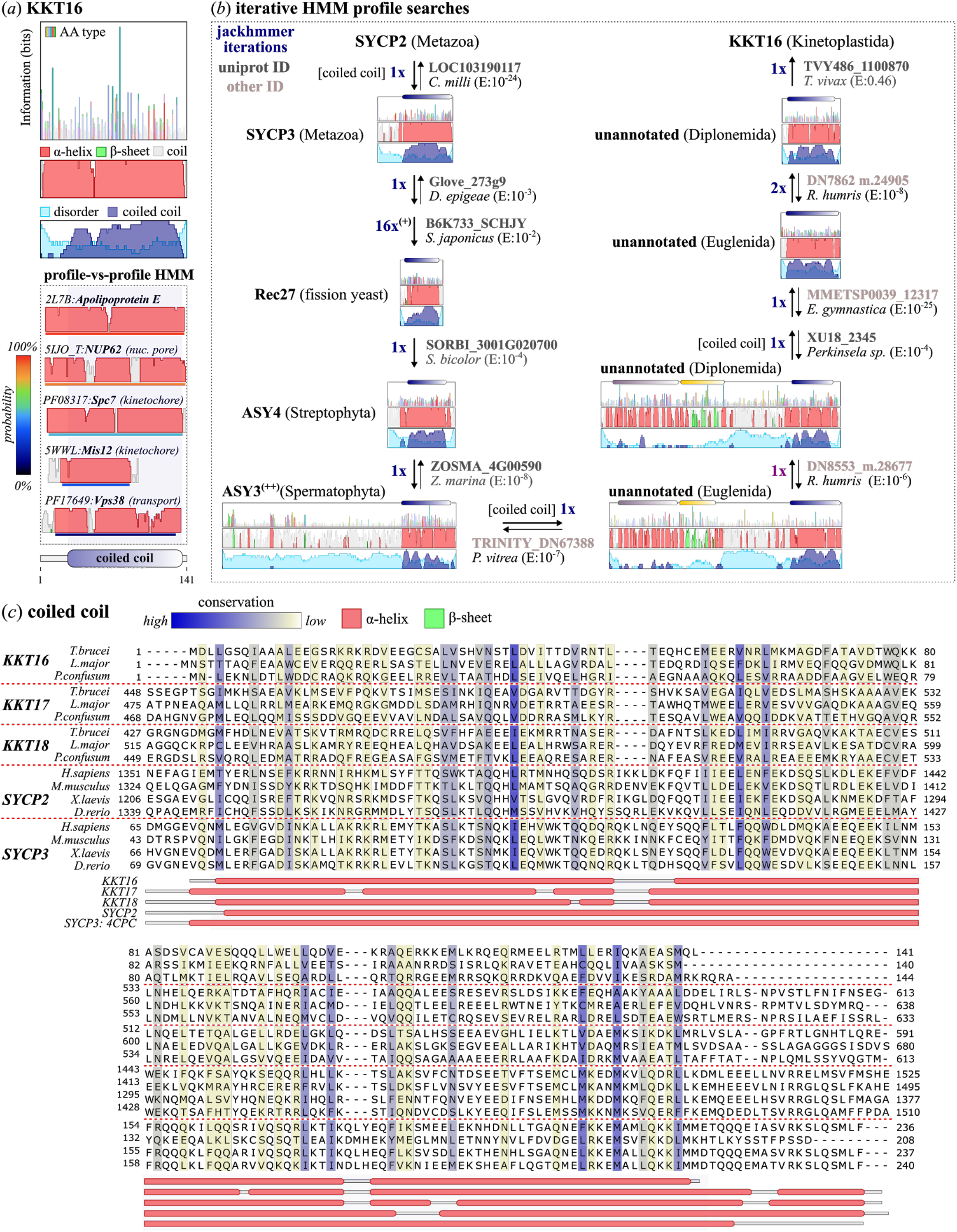
KKT16 is a coiled-coil protein homologous to the C-terminus of SYCP2 and KKT17,18. (A) Cartoon of the conserved domain architecture and sequence features of KKT16. See for explanation Figure 2a. The best scoring domains, similar to KKT16, are all part of coiled-coil proteins, but no clear connection could be established between KKT16 and SYCP2-SYCP3 or any other synaptonemal complex protein using profile-vs-profile HMM searches. (B) Cartoon of the similarity detection protocol (see Data & Methods) for establishing homology between the coiled coils of highly divergent SYCP^2-3^ proteins (SYCP2,3, Red1, Rec10,27, ASY3,4, KKT17,18) and KKT16. Similarity detection does not necessarily have to be performed in this order, but has been visualised in a linear manner to specifically showcase the long path towards establishing homology between the coiled coils of SYCP2,3 proteins to KKT16 (only a ‘grey zone hit’ with E-value 0.46). The seed HMM for metazoan SYCP2 is the same as shown in Figure 2b. Thick arrows and Evalues indicate the direction and significance of the similarity searches. Thin arrows in the reverse direction indicate that reciprocal searches yield similar homologous connections. Dark blue numbers indicate the number of iterations of HMM searches needed to include a particular sequence. Uniprot (dark grey) and non-Uniprot identifiers (lighter grey, searches performed on local sequence database, for sources see Table S2) indicate the highest scoring homologues after each iteration belonging to a larger phylogenetic group for which separate conservation and architecture cartoons are presented as in panel A (see File S3 for cartoons of all SYCP^2-3^ HMMs). The position of the ARM and PH domain and the coiled coil are projected on top of the cartoons (colours correspond to panel A). Brackets indicate when only selected domains are used for similarity detection. ^**(+)**^HMM used to detect all SYCP2-type and SYCP3-type homologues in Opisthokonta based on 16 iterations (fungi and animals, see File S3) ^**(++)**^ASY3 homologues in Spermatophyta (e.g. in *A. thaliana*) lost the N-terminal ARM and PH domains. Full species names: *Callorhinchus miiii; Diversispcra epigaea; Schizosaccharomyces japonicus; Sorghum bicdor; Zostera marina; Pioeotia vitrea; Rhynchopus humris; Eutreptieiia gymnastica; Trypanosoma vivax*. (C) Multiple sequence alignment of the coiled coils of KKT16–18 and SYCP2,3 homologues. See Figure 2c and File S5 for further explanation on annotation and the conservation colouring scheme. The secondary structure of SYCP3 is mapped based on the PDB file 4CPC (human SYCP3) [45].

Although the KKT16 complex subunits consist of generic domains, the specific ARM-PH topology of KKT17,18 was only detected in the HMM profile of one metazoan protein (Figure 2a): the Synaptonemal Complex Protein 2 (SYCP2, pdb:5IWZ [43], Prob>99%, E~10^-5^). In addition, the C-terminal coiled-coil domain of KKT17 and KKT18 showed similarity to a known SYCP2 multimerisation partner [44], the single coiled-coil protein Synaptonemal Complex Protein 3 (SYCP3, pdb:4CPC [45], Prob:95%, E~1). SYCP2:SYCP3 multimers constitute the axial elements of the synaptonemal complex (SC), a zipper-like structure that forms the linkage between parental chromosomes to facilitate homologous recombination during meiosis [32,46,47]. The significant similarity (E<10^-2^) and high HHsearch probability scores (Prob>85%) suggested that the KKT16 complex and the axial elements of the SC shared a common ancestor, providing an important clue to the possible origin of kinetoplastid kinetochores.

### KKT16–18 are part of the highly divergent SYCP^2-3^ gene family, including the SC components Red1, Rec10:Rec27, ASY3:ASY4 and SYCP2:SYCP3 found in Fungi, Archaeplastida and Metazoa

Although structures of the synaptonemal complex are highly conserved among eukaryotes [40], previous sequence similarity searches using BLAST, mostly found homologues of SYCP2 and SYCP3 in Metazoa, and failed to detect significant sequence similarity among the wide range of eukaryotes [48]. Based on our detection of similarity between the KKT16 complex in Kinetoplastida and the SYCP2:SYCP3 multimer in Metazoa, we sought to test for the presence of homologues among diverse eukaryotes using a more sensitive HMM-based approach. For ease of reference we use the term ‘SYCP^2-3^’ to indicate all genes similar to SYCP2,3 and KKT16–18. The term ‘SYCP2-type’ is used for those SYCP^2-3^ genes with an ‘ARM-PH-coiled coil’ domain topology (e.g. KKT17 and SYCP2). ‘SYCP3-type’ refers to other coiled coil-only genes (e.g. KKT16 and SYCP3).

To exploit sequence diversity in our search for putative SYCP^2-3^ homologues, we collated a large sequence database of 343 genomes and transcriptomes broadly sampled from the eukaryotic tree of life, but specifically focussed on including lineages more closely related to Kinetoplastida, such as Diplonemida, Euglenida and other species from the superphylum Discoba (see for full species list Table S2). Using an iterative HMM profile ‘hopping’ protocol for homology detection (see for detailed description Data & Methods), we identified candidate homologues throughout the eukaryotic tree of life (Figure 2b-c, Figure 3b-c, see for overview Table S2, File S2). We found that significant similarities (E<10^-2^) with the region spanning the last part of the ARM repeats and the full PH domain in combination with the presence of a C-terminal coiled coil, were the best criteria to distinguish SYCP2-type homologues from other eukaryotic ARM repeat or PH domain-bearing proteins. Importantly, we established homology based on significant sequence similarity between the KKT16 complex, SYCP2,3 and previously identified SC proteins Red1 (*Saccharomyces cerevisiae*), Rec10 (*Schizosaccharomyces pombe*), and ASY3:ASY4 (*Arabidopsis thaliana*) [44,49–51] (see for examples of linear homology detection paths: Figure 2b, Figure 3b). Red1 and Rec10 have a highly divergent N-terminal ARM-PH domain, but lack significant similarity between their C-terminal coiled coils and those in other SYCP^2-3^ genes (Figure 2b). It is unclear whether coiled coils in these proteins evolved convergently or diverged beyond recognition. However, based on the finding that Red1 and ASY3:ASY4 are structurally and functionally analogous to metazoan SYCP2:SYCP3 multimers [44], it is likely that these SC proteins share common ancestry with SYCP2 and SYCP3.

It is known that coiled coils are often unsuitable for sequence similarity searches and phylogenetic analyses due to a high degree of sequence redundancy. For example, searches with some clade-specific SYCP^2-3^ HMM profiles (e.g. metazoan SYCP3 and KKT17,18) yielded homologues of the metazoa-specific extracellular apolipoproteins after several iterations (Data & Methods), which are unlikely to be SYCP^2-3^ homologues based on their known function and location [52]. In most other searches we performed, however, coiled-coil domains of SYCP^2-3^ genes were sufficient to identify homologues using reciprocal iterative searches. To restrict inclusion of false positive coiled coil proteins, we only considered candidates with bidirectional best similarity to either: (1) *bona fide* eukaryotic SC components (Red1, Rec10, Rec27, ASY3, ASY4, SYCP2 and SYCP3), (2) KKT16 complex subunits, and/or (3) a SYCP2- or SYCP3-type gene of the same clade when a candidate for both is present (e.g. SYCP^2-3^ genes in Diplonemida and Euglenida, Figure 3b).

Interestingly, searches with HMM profiles of clade-specific SYCP2- and SYCP3-type genes often had overlapping candidate homologues in reciprocal searches, signifying potential duplications of their C-terminal coiled coils. For instance, closely overlapping candidate homologues were found between SYCP2 and SYCP3 in Metazoa, and between ASY3 and ASY4 in Streptophyta (Figure 3b). Similarly, we found that the *S. pombe* linear element (SC-like structure) components Rec10 and Rec27 likely share a common ancestor (Figure 2b, Figure 3b) [53]. KKT17,18 also showed evidence of homology to KKT16, albeit with borderline significance (0.01<E<1, Figure 3b). The homology of the C-terminal coiled coils of SYCP^2-3^ genes suggested that the KKT16–18 and Rec10:Rec27 complexes are likely based on the same coiled coil-mediated interactions found for SYCP2:SYCP3 (mammals), ASY3:ASY4 (plants) and Red1 (budding yeast) hetero- or homomultimers [44].

### The SYCP^2-3^ gene family evolved through recurrent duplications

Our identification of different numbers of homologues of both SYCP2 and SYCP3-type proteins among distant eukaryotic lineages, raised the question of what evolutionary events led to the diversification of the SYCP^2-3^ gene family: *How and when did these paralogues arise? Are they the result of lineage-specific duplications events or do they point to a more ancient origin for the SYCP2 and SYCP3-type genes?* To answer these questions, we inferred separate phylogenetic trees for the N-terminal ARM-PH region of SYCP2-type genes and the coiled-coil domains of all SYCP^2-3^ gene family members. Due to the divergent nature of SYCP^2-3^ homologues, we adopted a previously used alignment strategy [29] that generates a super alignment of distantly related sequences through iteratively aligning increasingly less similar clade-specific MSAs (Data & Methods). Some coiled-coil domains of the SYCP^2-3^ homologues were too divergent to yield MSAs of sufficient quality and were excluded from our analyses (e.g. the coiled coil of the SYCP^2-3^ genes in Fungi and SAR supergroups, see Data & Methods). To subsequently infer phylogenetic trees, we used the maximum-likelihood phylogenomics software IQ-Tree [54] (1000x Ultrafast bootstrap replicates, automated model selection, Data & Methods). To resolve duplications we reconciled the resulting phylogenetic trees of both the ARM-PH and coiled-coil tree with the known eukaryotic species tree [55]. Although the highly divergent nature of the SYCP^2-3^ sequences and short length of their coiled coils precluded the faithfull recovery of ancient patterns of eukaryotic evolution, they did provide evidence of more recent instances of gene duplication within well-resolved lineages (Figure 4).

**Figure 4.**
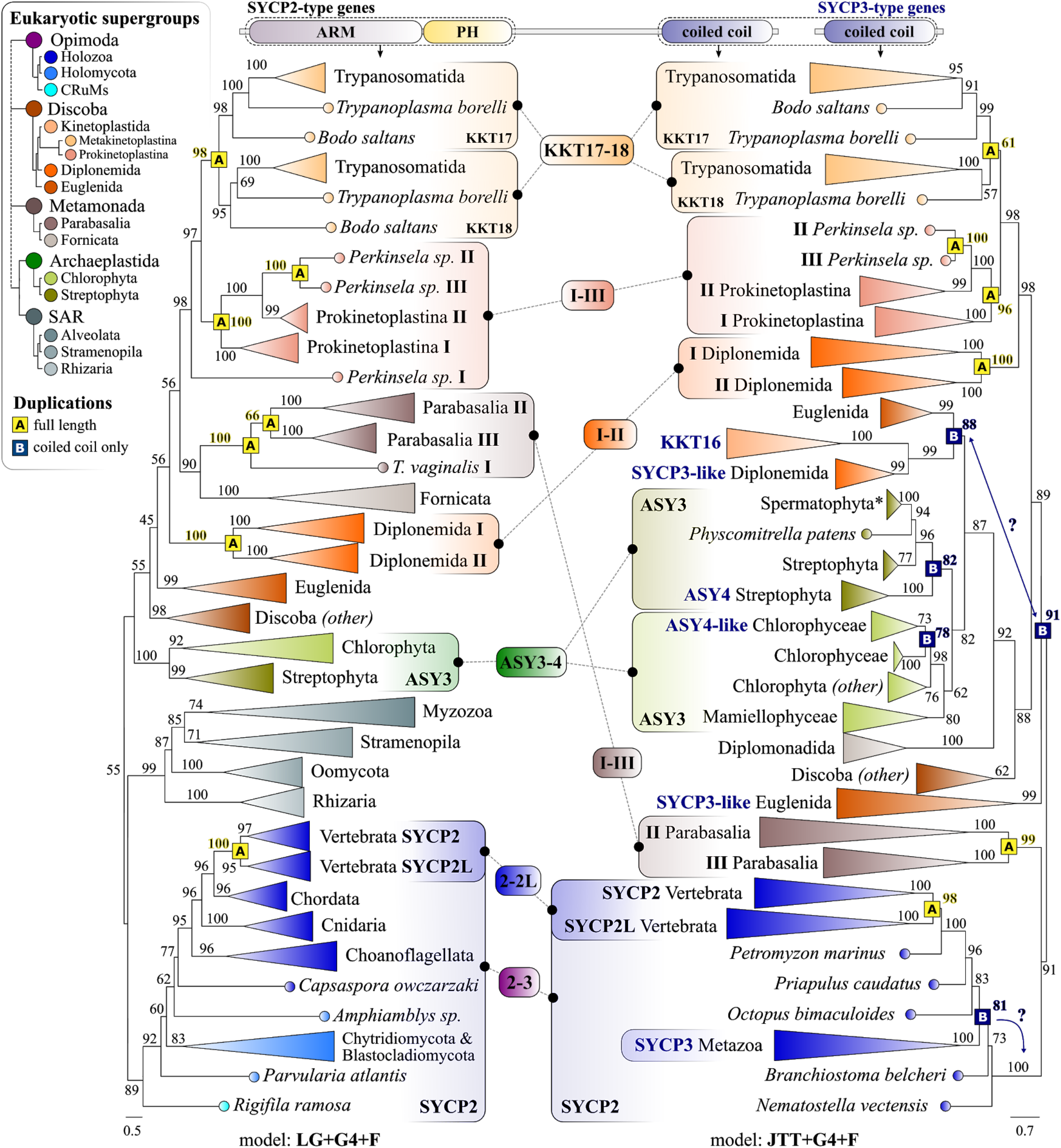
The SYCP^2-3^ gene family expanded through independent duplications. Mirrored phylogenetic trees (IQ-Tree: maximum likelihood, see Data & Methods) of the N-terminal ARM repeats and PH domain of SYCP2-type genes (left), and coiled coils of SYCP2 and SYCP3-type genes (right, names are in dark blue). A and B indicate independent duplications of either the full-length SYCP2-type genes (yellow, A) or the specific duplication of the C-terminal coiled coil (dark blue, B). Colours indicate taxonomic levels to which the various lineages belong (see legend top left). Values at branches indicate UltraFast Bootstrap support (1000x replicates: see Data & Methods), those associated with duplications are in bold and highlighted in yellow (full-length duplications) and dark blue (coiled coil duplications). Models of sequence evolution that were used to infer the phylogenetic trees are shown below each phylogram. Branch lengths are scaled and indicate the number of substitutions per site (see scale bar below each phylogram). A full overview of the uncollapsed phylograms, phylogenetic analyses details, and the underlying alignments can be found in File S4. (*) ASY3 in Spermatophyta (e.g. in *A. tha/iana*) lost the N-terminal ARM repeats and PH domain.

In total, we found evidence for seven independent duplications of full-length SYCP2-type homologues that were consistent between both ARM-PH and the coiled coil trees (see yellow A in Figure 4). Of these, we found three recurrent duplications among Kinetoplastida. One duplication in the common ancestor of the subclass Metakinetoplastina gave rise to KKT17 and KKT18, including in Trypanosomatida and Bodonida. Separate duplication events gave rise to the three SYCP2-type paralogues (I–III) present in the prokinetoplastid endosymbiont *Perkinsela sp*. (high support, bootstrap>95). Interestingly, we also detected an independent duplication event in the sister lineage to Kinetoplastida, the poorly described flagellate order Diplonemida. Why these duplications occur, especially in these two euglenozoan lineages, is unclear. It is known that SYCP2L, the vertebrate-specific paralogue of SYCP2 (bootstrap support>95), localises at centromeres during specific stages of meiosis [56], indicating that the centromeric localisation of SYCP^2-3^ paralogues is not unique to KKT17,18. Furthermore, we found two duplications that gave rise to the three SYCP2-type paralogues (I–III) present in the metamonad parasite *Trichomonas vaginales* of the class Parabasalia. Lastly, we also found a species-specific duplication of the SYCP2-type gene ASY3 (I-II) in *Chlamydomonas reinhardtii* (not shown in Figure 4, see File S4).

In a similar fashion, we traced the origins of SYCP3-type genes to four independent duplications of the coiled-coil domains of clade-specific SYCP2-type ancestors, albeit generally with lower bootstrap support and positions less clearly reconcilable with the eukaryotic tree of life (see dark blue B, Figure 4). In Archaeplastida, two independent duplications gave rise to an SYCP3-type paralogue in both Streptophyta (ASY4), and the green algal class Chlorophyceae (ASY4L). In accordance with our HMM searches (Figure 2a, Figure 3b), the closest homologue of SYCP3 was SYCP2, signifying a duplication of the coiled-coil domain in the common ancestor of Metazoa (Figure 4). The close association of KKT16 in Kinetoplastida with SYCP3-type genes in Diplonemida (bootstrap support: 99) and SYCP2-type homologues from Euglenida (bootstrap support: 88), suggested that these duplications must have occurred in the ancestral Euglenozoa, and that both SYCP2- and SYCP3-type genes were present in this ancestor (Figure 4).

Altogether, our phylogenetic analysis indicated that SYCP2-type paralogues and SYCP3-type genes originated independently through recurrent duplications of multiple SYCP2-type ancestors at various taxonomic levels. Because we could not detect any SYCP^2-3^ proteins in prokaryotes, we considered all eukaryotic members of the SYCP^2-3^ gene family a single orthologous group, which was likely descendant from a single SYCP2-type gene present in the Last Eukaryotic Common Ancestor (LECA). A graphical overview of the evolutionary scenario for SYCP^2-3^ duplications, phylogenetic profiles and inferred ancestral states is summarised in Figure 5.

**Figure 5.**
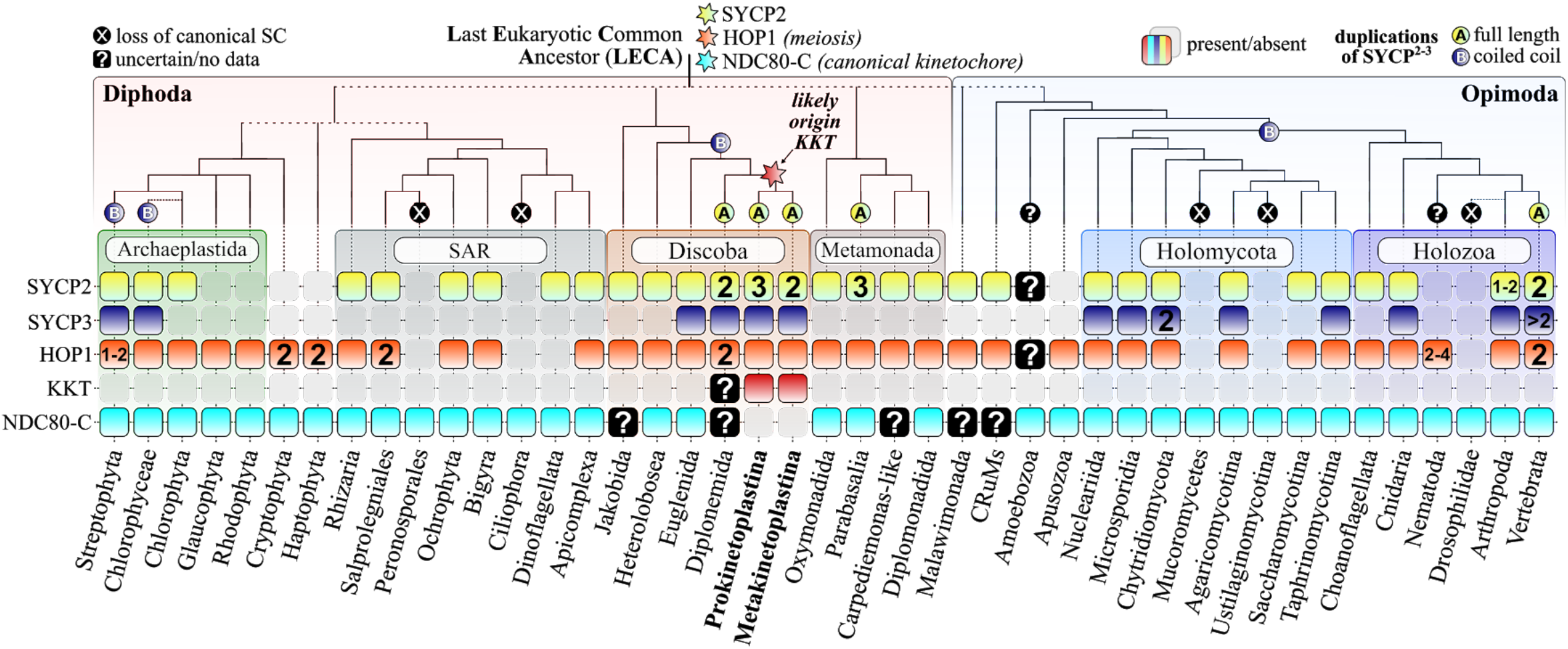
Evolutionary scenario for the SYCP^2-3^ gene family in the light of meiosis and kinetochores. Phylogenetic profiles of SYCP^2-3^ and HOP1 orthologues, and the kinetoplastid and canonical kinetochore throughout the eukaryotic tree of life. See for sequences Table S2 and File S2. Canonical kinetochore (cyan, harbouring NDC80-based kinetochores), kinetoplastid kinetochore (dark red) and question marks (uncertain/no data) indicate evidence for the type of kinetochore proteins found among each eukaryotic lineage (see Table S2 for references and comments). Cartoon and classification of the eukaryotic tree of life, and the position of the Last Eukaryotic Common Ancestor (LECA) is guided by Burki *et al*. [55]. Numbers indicate the amount of paralogues found among particular lineages. ‘A’ and ‘B’ refer to the two types of independent duplications found for the SYCP^2-3^ gene family similar to Figure 4. (X) indicate co-loss of SYCP^2-3^ and HOP1 (e.g. Ciliophora and Ustilaginomycotina). Hexagrams (stars) indicate the likely origin of each feature/gene family.

## Discussion

### Widespread conservation of the SYCP^2-3^ gene family in diverse eukaryotes points to an ancient origin for axial elements of the synaptonemal complex

This study revealed similarities in KKT16–18 to the axial elements of synaptonemal complexes (SC). This was an unexpected finding because KKT16–18 are kinetochore proteins present in mitotic trypanosomes cells, while the SC is a strictly-meiotic structure [40]. During meiosis, chromosome axes assemble on each pair of sister chromatids and serve as a platform from which chromatin loops radiate. The SC subsequently assembles in between the axes of homologous chromosomes along their lengths, forming a conspicuous zipper-like ultrastructure visible by electron microscopy (Figure 6a). [32,46,47]. The structure of the SC is widely conserved among diverse eukaryotes and appears as a tripartite protein structure that consists of two axial elements (also known as lateral elements) that flank a central element to which they are connected via transverse filaments (Figure 6a) [57]. In vertebrates, components of axial elements include SYCP2, SYCP3, meiotic HOP1/HORMAD family proteins, and cohesin complexes [58]. HOP1/HORMADs and cohesins are conserved in many eukaryotes (Figure 5) [8,59,60]. In contrast, it has been difficult to detect homologues for SYCP2 and SYCP3 or to establish their significant sequence similarity with SC components identified in other model systems (e.g. Red1 in *S. cerevisiae*, Rec10 in *S. pombe*, and ASY3:ASY4 in *A. thaliana*) [48].

**Figure 6.**
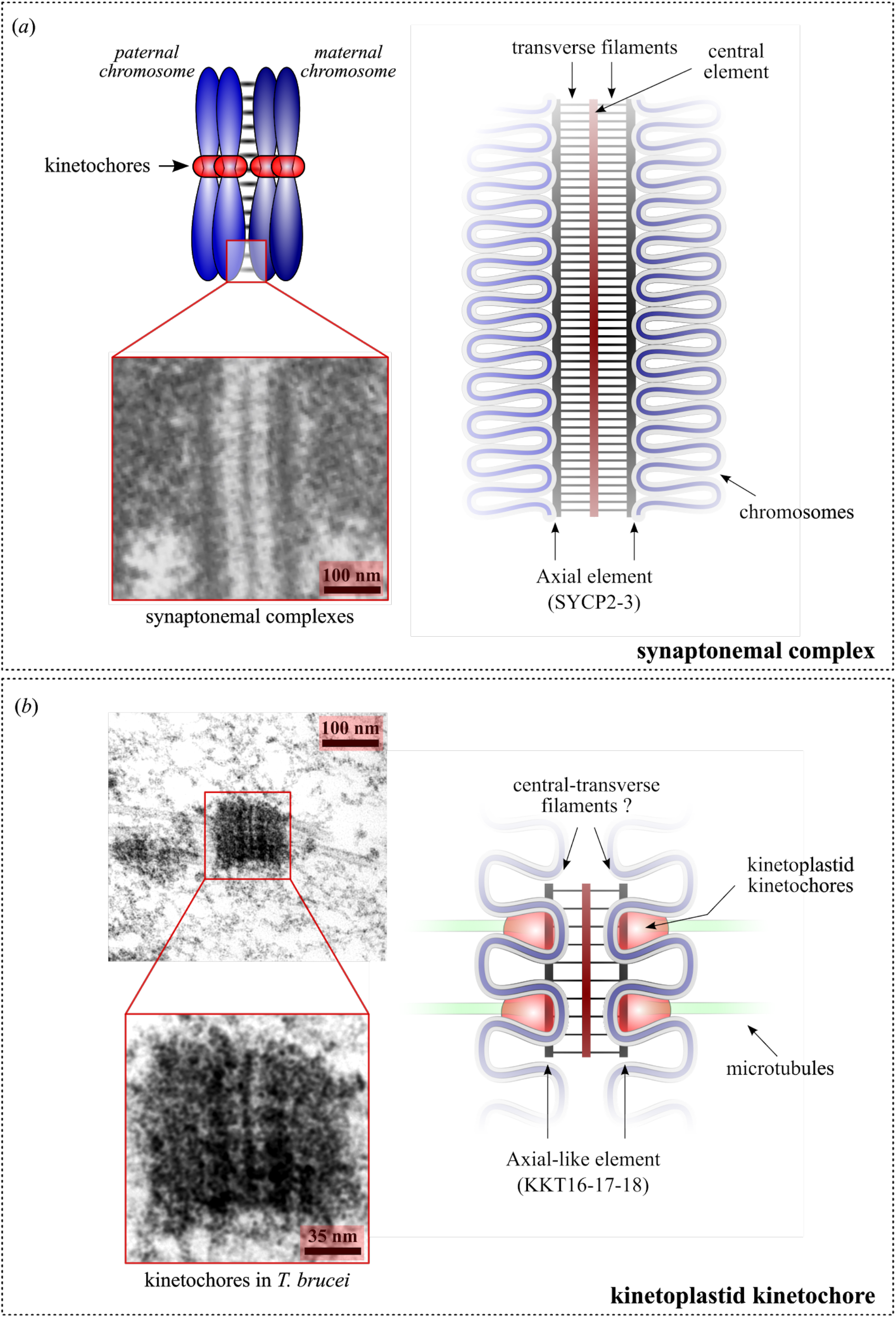
Genomic and microscopic similarity of kinetoplastid kinetochores and the synaptonemal complex suggests a common origin. (A) Schematic of the meiotic synapsis (left) and the synaptonemal complexes (right). SYCP2 and SYCP3 are components of the axial element. Zoom in is an electron micrograph of a mouse spermatocyte, showing synaptonemal complexes in between homologous chromosomes. Reproduced from [75] under CC-BY. (B) Left: An electron micrograph of a mitotic trypanosome cell, showing that the putative kinetochore structure attaches spindle microtubules from opposite poles. Adapted from [28] with permission from Springer Nature. Right: A hypothetical model showing that the KKT16 complex may form an axial element-like structure in the kinetoplastid kinetochores.

In this study we used a remote homology detection protocol combining both profile-versus-profile and iterative HMM searches to identify highly divergent orthologues of the metazoan SYCP2:SYCP3 multimers and subunits of the KKT16 complex (Figure 2, Figure 3). Specifically, we made use of the ‘hopping’ strategy which was previously employed to detect divergent homologues of canonical kinetochore proteins [8,61,62]. The ‘hopping’ strategy uses the following logic: ‘if A is homologous to B, and B is homologous to C, then A must be homologous to C’. This approach has the potential to find many more orthologues of highly divergent gene families such as those involved in meiosis and the kinetochore, but inclusion criteria other than sequence similarity must be considered to prevent false positive candidate orthologues. In particular, for SYCP^2-3^ proteins, which harbour generic domains such as ARM repeats and coiled coils, we selected only those candidate sequences that either had the full-length SYCP2-type ARM-PH-coiled coil topology or coiled coil-only proteins that showed significant similarity in reciprocal HMM searches with either known SC components or KKT16–18 (e.g. ASY3, KKT17 or SYCP3) and/or a full-length SYCP2-type sequence of the same eukaryotic lineages. These stricter criteria potentially resulted in exclusion of SYCP3-type coiled coil-only genes among eukaryotic lineages, for instance in Apicomplexa and other phyla from the SAR supergroup (see absences in Figure 5). Further experimental validation of the localisation and molecular function of such candidates from the SAR supergroup will be needed to gain more confidence that these genes would be part of the SYCP^2-3^ gene family and exert a function in the SC.

In addition to showing that experimentally verified axial SC components of several models (animals, fungi and plants) and KKT16–18 belong to a common gene family, we identified SYCP^2-3^ orthologues in all eukaryotic supergroups (Figure 5), although not in any prokaryote. The widespread presence of these proteins suggests that axial elements were part of the ancient eukaryotic SC and that SYCP^2-3^ genes were likely present in the LECA (Figure 5). We detected highly divergent SYCP^2-3^ orthologues in Metamonada (e.g. *Giardia intestinalis* and *Trichomonas vaginales*), Microsporidia (e.g. *Encelphalitozoon intestinalis*) and a wide variety of fungi (e.g. *Neurospora crassa, Fusarium oxysporum, Spizellomyces punctatus* see Table S2, File S2). We specifically searched for SYCP^2-3^ genes in Drosophila-related lineages, since they seem to have a largely analogous SC and SYCP^2-3^ genes were previously not identified [63]. We found divergent SYCP^2-3^ orthologues in various insect lineages (Lepidoptera, Diptera), but none in Drosophilidae, signifying the specific loss of SYCP^2-3^ genes in this lineage (Figure 5). Interestingly, we also did not detect any SYCP^2-3^homologues in lineages which were previously described to lack SC structures or pathways of meiotic recombination, such as Ciliophora [64,65], Amoebozoa [66], and Ustilaginomycotina [67,68]. The status of Amoebozoa was somewhat unclear as we found one SYCP2-type gene in the amoeba *Planoprotosteiium fungivorum* (Figure 5), but no significant sequences were found in the reciprocal similarity searches. In contrast, we detected candidate homologues in lineages with previously described SC-like structures during meiosis, such as Apicomplexa [69] and Oxymonadida [70]. The general concordance between the presence of SYCP^2-3^ genes and SC-like structures suggests that these genes could be used as a good predictor for the presence of canonical SCs. However, we also found cases where SC-like structures were described, but no SYCP^2-3^ genes were detected, such as Bacillariophyta (diatoms) and in Rhodophyta (red algae) [57,71,72]. To examine the possibility of missed detection of SYCP^2-3^, we searched for orthologues of the meiotic HORMA domain protein HOP1/HORMAD, which is a meiosis-specific interactor of SYCP2-type proteins [44,58] and is typically expected to co-occur with the presence of canonical SC and meiosis [50,63]. The phylogenetic profiles of SYCP^2-3^ and HOP1 corresponded well (31/43 shared presences and absences, see Figure 5), but we found six lineages (Rhodophyta, Glaucophyta, Cryptophyta, Haptophyta, Apusozoa and Nematoda) that do contain HOP1, but not SYCP^2-3^ (Figure 5). Conversely, we detected several highly divergent SYCP2-type proteins among dinoflagellates, in contrast to a recent report [73], while such lineages lacked HOP1 (see Table S2, File S2). Whether these discordances between HOP1 and SYCP^2-3^ point to a lack of homology detection or whether SC-like structures in these lineages contain analogous SC components like in Nematoda [74] and Drosophilidae [63] remains unclear. In any case, the absence of HOP1 and SYCP^2-3^ in Mucoromycetes, Zoopagomycota (see Table S2) and the oomycote class Peronosporales (potato blight pathogen *Phytophthora infestans*) likely signifies the absence of a canonical SC structure in these lineages.

We detected 14 recurrent duplications for the SYCP^2-3^ gene family. Why these duplications occurred remains unclear. In the case of the lineages-specific coiled coil-only duplications that gave rise to different SYCP3-type genes (i.e. SYCP3, KKT16, ASY4, ASY4L), we speculate a specific need for the coiled coil to facilitate the formation of axial element-like structures apart from the function of the N-terminal domain ARM-PH, which is currently unknown. In the case of SYCP2-type genes there is only limited data available on the two paralogues present in vertebrates: SYCP2 and SYCP2L. SYCP2L is expressed specifically in oocytes and localises at SCs and centromeres [56], although its function remains unclear. It is noteworthy that both Kinetoplastida and Diplonemida show recurrent duplications of SYCP2-type genes (Figure 4, Figure 5). This apparent increase in paralogues might correlate with new functions of these proteins in the kinetochore rather than in the SC. It will therefore be of interest to study SYCP^2-3^ homologues in Diplonemida and assess whether these proteins play a role in the kinetochore and/or the SC.

### Hypothesis: kinetoplastids repurposed meiotic structures to build kinetochores

Beyond the shared ancestry of the KKT16 complex and axial element components of the synaptonemal complex, other KKT proteins have conserved domains with relevance to homologous recombination and chromosome synapsis (i.e. BRCT domain, FHA domain, and Polo-like kinases). We therefore propose that ancient kinetoplastid ancestors repurposed parts of the meiotic machinery to assemble a kinetochore-like structure by restricting its formation to one chromosomal region and acquiring microtubule-binding activities (Figure 6b). This hypothesis could explain the unique organisation of kinetoplastid kinetochores that lack a clear gap between sister kinetochores even at metaphase [25–28] and, indeed, are strikingly similar in structure to synaptonemal complexes (Figure 6) [75].

Functions of the KKT16 complex at the kinetoplastid kinetochore remain unknown. Our previous mass spectrometry analysis of the KKT16 complex purifications from mitotically-growing cells did not reveal significant co-purification of cohesin subunits or HOP1 [16]. It is therefore currently unclear whether the KKT16 complex has a similar function to SYCP^2-3^ homologues found in other eukaryotes. Because KKT16–18 are the only members of the SYCP^2-3^ gene family present in kinetoplastids, it will be interesting to examine whether KKT16–18 are also used as components of synaptonemal complexes during meiosis, which takes place when trypanosomes transmit in the salivary glands of the tsetse fly vector [76]. Although the main functions of the SC are to hold homologous chromosomes together and promote recombination, it is known that SC components have non-canonical functions in certain lineages. For example, some organisms rely on the SC or its components for connection between homologous chromosomes beyond prophase I (when the SC is disassembled in most organisms). In the female silkworm *Bombyx mori* that lacks meiotic recombination, homologous chromosomes are joined together by the retention of modified SCs until metaphase I [77]. Similarly, some SC components remain at the centromeric or non-centromeric region and promote biorientation of non-exchange chromosomes [78]. Functions of the KKT16 complex remain unknown, so it will be interesting to test whether it plays any role in connecting and properly orienting sister chromatids in trypanosomes.

### Origins of kinetoplastid kinetochores in the light of early eukaryotic evolution

#### Why do kinetoplastids have unique kinetochores, while other eukaryotes have canonical kinetochore proteins?

The absence of canonical kinetochore proteins among Kinetoplastida provides several hypothetical scenarios for the evolution of the kinetoplastid kinetochore. In the case that the first common ancestors of Kinetoplastida possessed the canonical kinetochore system, it must have been secondarily lost, and presumably replaced by the new unique kinetochore system now found in this group (Figure 5). This scenario seems to be most consistent with the current consensus on the eukaryotic tree of life, where Kinetoplastida are part of the phylum Euglenozoa, which also includes Diplonemida, Symbiontida and Euglenida [55]. Furthermore, most eukaryotes have the canonical kinetochore system [8], including euglenids [79] (Figure 5). It is noteworthy that an initial survey of diplonemid transcriptomes found only limited evidence for the presence of a canonical kinetochore system with putative candidates for the centromeric H3 variant CENP-A, but no subunits of the NDC80 complex or other structural kinetochore components [39] (Figure 5). Intriguingly, no clear orthologues of the kinetoplastid kinetochore proteins could be identified either [39], suggesting that Diplonemida might potentially have yet another kinetochore system. Identification of kinetochore proteins in Diplonemida and in-depth sequence analyses of these and kinetoplastid kinetochore proteins using our sensitive homology detection workflow will be needed to shed further light on how ancestral kinetoplastids acquired a unique kinetochore system.

An alternative scenario is that early kinetoplastid ancestors never possessed the canonical kinetochore system. There is a controversial hypothesis that places the root of the eukaryotic tree of life between Euglenozoa (or deeply within Euglenozoa) and all other eukaryotes [80]. In this ‘Euglenozoa-first’ scenario (discussed in [21]), it is possible that kinetoplastid kinetochores and canonical kinetochores were invented independently and they are both derived systems, meaning that ancestral eukaryotes might have possessed a chromosome segregation machinery that does not exist anymore today. It is still unclear whether mitosis or meiosis evolved first [81–85]. In contrast, it is known that some species of Archaea are capable of homologous recombination and cell fusion [86]. Although we have been unable to find any SYCP^2-3^ genes in Archaea or Bacteria, the widespread presence of SCs among eukaryotes, including Euglenozoa, suggests that SCs were likely present in the LECA (Figure 5). Under the Euglenozoa-first hypothesis, our findings that KKT16 complex subunits have similarities to SC components raise a possibility that some features of meiosis (i.e. chromosome synapsis and genetic exchange) might have evolved prior to an active chromosome segregation mechanism that relies on kinetochores and spindle microtubules.

### Concluding remarks

Although the kinetochore is at the heart of chromosome segregation, substantial compositional diversity and rapid sequence evolution of its subunits is widespread throughout the eukaryotic tree of life. This presents us with fundamental questions: *how can kinetochores be essential and divergent at the same time? How (and why) do cells replace one kinetochore system with another?* While Drinnenberg *et al*. used the elegant analogy with the ‘ship of Theseus’ to explain this remarkable evolutionary behaviour of kinetochores [10], the radically different composition of the kinetoplastid kinetochore seems to be at odds with such a piece-by-piece replacement model. Our study provides a new concept for understanding such an extreme jump in the evolution of kinetochores in eukaryotes, namely the apparent ability to adapt and repurpose meiotic complexes for mitotic functions. Further functional and evolutionary characterisation of divergent kinetochores in Kinetoplastida and other eukaryotes will thus not only benefit our understanding of their inner workings, but also shed light on how this core cellular system has been allowed to diverge in such a radical fashion.

## Data & Methods

### Primers and plasmids

Primers used in this study are listed in Table S1. To make pBA198 (6His-KKT16), KKT16 was amplified from genomic DNA using BA509/BA510 and cloned into the EcoRI/HindIII sites of the pST44 polycistronic expression vector (RRID: Addgene_64007) [41]. To make pBA200 (KKT18, 6His-KKT16), KKT18 was amplified from genomic DNA using BA511/BA512 and cloned into pBA198 using XbaI/BglII sites. To make pBA202 (KKT18, 6His-KKT16, KKT17), KKT17 was amplified from genomic DNA using BA513/BA514 and cloned into pBA200 using KpnI/MluI sites.

### Expression and purification of the recombinant KKT16 complex

pBA202 (KKT18, 6His-KKT16, KKT17) was transformed into Rosetta 2(DE3)pLysS *E. coli* cells (Novagen, 71403). Cells were grown in LB media (Fisher Scientific, BP1426-2) at 37°C to an OD600 of ~0.6 and protein expression was induced by 0.2 mM Isopropyl β-d-1-thiogalactopyranoside (IPTG) (Sigma-Aldrich, I6758) at 18°C overnight. Recombinant proteins were purified using an Ni-NTA Fast Start Kit under native condition (Qiagen, 30600). To check the expression of KKT16 complex subunits, protein expression was induced by 0.2 mM IPTG at 37°C for 3 hours.

### Secondary structure-guided profile-vs-profile HMM comparisons and visualisation

Distant homologues of KKT16 complex members (KKT16–18) were extracted from a previous extensive survey of kinetoplastid kinetochore proteins by Butenko *et al*. [39]. For each subunit we constructed multiple sequence alignments (MSAs) using MAFFT (v7.475 [88], RRID: SCR_011811: option ‘eins-i’ or ‘lins-i). Secondary structure-annotated profile Hidden Markov Models (HMMs) were constructed based on MSAs of both full-length sequences and domains (ARM repeats, PH and coiled coil) using the ‘hhmake’ script from HHsuite3 (RRID: SCR_010277 [42]). To map the domain architecture and potentially uncover highly divergent homologues of KKT16 complex members, we searched the pre-compiled pdb70 (RRID:SCR_012820) and pfam-A (RRID: SCR_004726) profile HMM databases from the HHsuite repository (link, downloaded November 1, 2020), using the KKT16 complex subunit HMM models (see File S1 for HHsearch text output for full-length and subdomain HMM-vs-HMM profile searches). All HMMs (including iterative HMM searches, see below) used during this study were annotated and visualized using custom python scripts (see File S3 for HMM and MSA text files and visualisation, and File S6 for relevant settings and sources for visualisation). The following predictions/annotations were included: (1)amino acid conservation (Shannon entropy: bits of information), derived via the Skylign API, RRID: SCR_001176 [89]), (2) secondary structure (PSIPRED, RRID:SCR_010246 [90]), (3) coiled coil (DeepCoil version 1.0: https://github.com/labstructbioinf/DeepCoil) [91]), (4) intrinsic structural disorder (IUPRED version 2a, RRID: SCR_014632 [92]). MSA columns that were not present in either the first or the MSA consensus sequence were removed to ensure gapless HMM visualisation. Plots were generated for each HMM and manually compiled into figures using the open source scalable vector graphics editor Inkscape 1.0rc1 for macOS (Inkscape Project 2020, retrieved from https://inkscape.org, RRID: SCR_014479).

### Sequence database

We compiled a large sequence database consisting of 343 (single cell) genomes and transcriptomes of a wide variety of eukaryotes [8,93,94] (see Table S2 for sources). We specifically focussed on including lineages closely related to Kinetoplastida, such as Diplonemida, Euglenida and other Discoba [39,95,96], and taxa related to lineages that lack canonical synaptonemal complex proteins (e.g. nematodes, Drosophilidae, Ciliophora and Amoebozoa). For several species it was not possible to obtain annotated protein coding regions. In these cases, we used TransDecoder v5.5.0 to predict open reading frames (https://github.com/TransDecoder/TransDecoder).

### Supervised remote homology detection (‘hopping’) protocol

Because of the highly divergent sequence composition of KKT16–18 and other axial element components of the SC found in diverse eukaryotes, we employed a supervised homology detection protocol to optimise multiple HMMs based on iterative reciprocal similarity searches using jackhmmer and hmmsearch (HMMER 3.1 and 3.3; RRID:SCR_005305 [97]. Searches were executed with standard inclusion thresholds until convergence, unless otherwise specified. HMMs were constructed using ‘hmmbuild’. Our protocol was based on the following steps/considerations:

a. To increase initial search sensitivity, we constructed profile HMMs of automatically defined clade-specific orthologous groups (OrthoFinder 2.0, RRID: SCR_017118 [98]: standard settings) of KKT16–18 and other axial element components (e.g. SYCP2,3 and ASY3,4). We used both full-length and subdomain (ARM, PH, coiled coil) HMMs as seeds for iterative sequence searches.
b. When queries using seed HMMs returned few hits, we searched Uniprot (RRID: SCR_002380) with the jackhmmer web server (https://ww.ebi.ac.uk/Tools/hmmer/search/jackhmmer [99]) for multiple iterations until no new putative candidate homologues could be included (E<0.01, domain E<0.03).
c. Iterative HMM searching is highly sensitive, and either too many homologues or other non-homologous sequences can be included by mistake due to the presence of highly common domains or due to local sequence biases such as coiled-coil regions. To prevent the inclusion of potentially false positive candidates, we used either the full-length sequence or other, non-common domain/coiled-coil regions of these new candidates as a query for jackhmmer searches. In case neither searches yielded reciprocal hits after numerous iterations or hits were clearly part of non-homologous proteins (with other domains or lacking either the ARM repeats or coiled coil), they were discarded as putative homologues. For example, we noted that multiple iterations with full-length SYCP2-type HMM profiles resulted in the frequent eventual inclusion of homologues of long ARM repeat proteins such as Vac8p and APC (also found to be similar based on HHsearch: Figure 2a). Such ARM repeat similarities point to more ancient homologous connections between these groups of proteins, but the absence of a PH domain and C-terminal coiled coil prompted us to remove these sequences to prevent their inclusion as candidate SYCP2-type genes in subsequent iterations. Altogether, we found that significant similarities with the region spanning the last part of the ARM repeats and the PH domain in combination with the presence of a C-terminal coiled coil, were the best inclusion criteria to distinguish between SYCP2-type homologues and other eukaryotic ARM repeat or PH domain-bearing proteins. Our profile HMM searches frequently returned metazoan apolipoprotein sequences as putative homologues of SYCP2 and SYCP3-type proteins (see also the HHsearch output in Figure 2a, Figure 3a) through similarity of their coiled-coil regions. While these extracellular lipid-binding proteins could formally be homologous to synaptonemal complex proteins, we deemed it more likely that the coiled coils of these proteins evolved convergently. To restrict further inclusion of false positive coiled coil homologues, we only included candidates with bidirectional best similarity to either to experimentally verified eukaryotic SC components (Red1, Rec10 and ASY3,4, SYCP2,3), subunits of the KKT16 complex, and/or (3) a SYCP2- or SYCP3-type gene of the same clade when a candidate for both is present.
d. Due to the highly divergent nature of SC components and KKT proteins we could not establish a single optimised profile HMM that captured all orthologous sequences. Instead in many instances we used the sequence ‘HMM hopping’ method, which follows from the logic that homology is a transitive feature by nature: if A is homologous to B, and B is homologous C, then A is homologous to C. Where possible reciprocal ‘HMM hopping’ searches were performed to increase our confidence in distant homologous relationships. In the case only unidirectional searches yielded new candidates or experimentally verified SC components, searches were repeated with more permissive bit scores (18-25) or E-values (0.1-1) to achieve reciprocal homologous relationships. Such cases were specifically scrutinised for similarity in secondary structure as well as their sequence composition. Examples of profile HMM hopping schemes for establishing homology between SC components and KKT16–18 are visualised in Figure 2b and Figure 3b.
e. If any of the iterative (reciprocal) searches using all of the (clade-specific) seed and optimised profile HMMs yielded overlapping hits and met the criteria mentioned above, we included these sequences as orthologues (see File S2 for SYCP2 and SYCP3-type sequences). HMMs of clade-specific SYCP^2-3^ genes used to establish the homology between SYCP2,3, ASY3,4, Red1, Rec10,27 and KKT16–18 can be found in File S3.

### Phylogenetic analyses

Due to the highly divergent nature of the SYCP^2-3^ gene family, generating a high quality MSA including all sequences was not feasible. We therefore adopted a previously used [29] iterative alignment protocol to generate a super alignment consisting of separate clade-specific MSAs (see Data & Methods for definition of clade-specific MSAs) using the ‘merge’ option in MAFFT (MAFFT v7.475 [88], RRID:SCR_011811: merge, ginsi, unalignlevel 0.6). Before addition to the super alignment, each clade-specific MSA was trimmed with trimAl (v1.4.rev22, RRID: SCR_017334 [100]) to remove unconserved positions. The order of MSA merging was determined based on bidirectional next-best sequence recovery using hmmsearch [97]. For instance, the coiled coil of metazoan SYCP2 and SYCP3 were closest, as reciprocal searches using the SYCP2 HMM yielded SYCP3 homologues as best next hits, and vice versa. MSAs were manually scrutinised for apparent misalignments, and re-run using either the MAFFT ‘eins-i’ or ‘linsi-i’ option to yield better alignments. This procedure was performed separately for the N-terminal ARM repeats and PH domain of SYCP2-type homologues and C-terminal coiled coils of SYCP2 and SYCP3-type homologues. The coiled coils of SYCP2 and SYCP3-type proteins in species of the SAR supergroup, Fungi (e.g. Rec10, Rec27 and Red1), *Bodo saltans* KKT18, *Perkinsela sp*. I and *Trichomonas vaginales* I, and the N-terminal ARM-PH domain of Red1/Rec10-like homologues, were too divergent to yield MSAs of sufficient quality. Furthermore, phylogenetic tree inference including these sequences showed signs of long-branch attraction, and were therefore left out of the phylogenetic analysis. For the final super alignments, only positions with a column occupancy higher than 30% (ARM repeats + PH domain) and 70% (coiled coil) were considered for further analysis. Maximum-likelihood phylogenetic analyses (phylograms shown in Figure 4, for data files see File S4) were performed with IQ-Tree (version 1.6.12, RRID: SCR_017254) with automatic substitution model selection using ModelFinder [101], a GAMMA model of rate heterogeneity, and 1000 Ultrafast bootstrap and SH-like approximate likelihood ratio test replicates [54]. Parameters for the final phylogenetic analyses: ARM repeats + PH domain (alignment: 424 positions, model: LG+G4+F); coiled coil (alignment: 165 positions, model JTT+G4+F). Trees were visualised and annotated using FigTree v1.4.4 [102].

## Data and Code accessibility

All data, annotations and scripts are included in the manuscript. The scripts to execute the visualisation of the secondary structure, conservation and the HHsearch output cannot be shared directly due to local computational environment dependencies. A detailed description/manual is available to replicate these analyses/visualisations (see Data & Methods and File S6 for description). The same supplementary files were also deposited in the FigShare repository at DOI: 10.6084/m9.figshare.13725787.

## Authors’ contributions

E.C.T. performed all the bioinformatic analyses and compiled the scripts to generate the figures. T.A.W. assisted in setting up the evolutionary analysis using the HHsearch algorithm and wrote the associated python script for visualisation of the results. R.F.W. supervised the project. B.A. reconstituted the KKT16 complex and noticed the similarity between the SC and trypanosome kinetochore architectures in electron microscopy images. B.A. and E.C.T. wrote the paper together with input from R.F.W. and T.A.W. All authors gave final approval for publication and agreed to be held accountable for the work performed therein.

## Competing interests

We declare we have no competing interests.

## Funding

E.C.T was funded through a personal Postdoctoral Research Fellowship awarded by the Herchel Smith Fund at the University of Cambridge (United Kingdom). B.A. was supported by a Wellcome Trust Senior Research Fellowship (grant no. 210622/Z/18/Z) and the European Molecular Biology Organisation Young Investigator Program. R.F.W. was funded by the Wellcome Trust (grant no. 214298/Z).

## Acknowledgments

We thank Alastair Simpson, Gordon Lax and Julius Lukeš for providing access to Euglenida, Diplonemida and/or Kinetoplastida transcriptomes and genomes before publication, and Keith Gull for providing an original electron micrograph of *T. brucei* kinetochores. We also thank Kim Nasmyth and David Sherratt for discussion. We thank members of the Akiyoshi and Waller labs for feedback.

## Supplementary Material

### Figure S1

Expression test of the KKT16 complex subunits. pBA198 (6His-KKT16), pBA200 (KKT18, 6His-KKT16), and pBA202 (KKT18, 6His-KKT16, KKT17) were transformed into *E. coli*. Protein expression was induced with IPTG at 37°C for 3 hours. Coomassie-stained 12% acrylamide gel of the purification is shown. M: marker.

### Table S1

List of primers used in this study.

### Table S2

Presence/absence profiles of the SYCP^2-3^ and HOP1 gene family in eukaryotes, including: (1) the sources for 343 genomes and translated transcriptomes, (2) taxonomy for each species, (3) numbers of paralogues per species for SYCP2, SYCP3 and HOP1-like genes, (4) notes and gene names of SYCP2,3 and HOP1 orthologues, (5) information evidence for the kinetochore archetype present in each species (canonical kinetochores: CENP-A and or NDC80 based, kinetoplastid kinetochore: KKT/KKIP proteins present, unknown: no data available, likely canonical: annotated as such, given the presence of canonical kinetochores in related lineages). *Of note:* orthologues of the meiotic HOP1 HORMA protein were searched in our local database based on previously established HMM models with [8,29]. Divergent candidates were further included based on the presence of extra non-HORMA domains/motifs found among other orthologues (i.e. winged-helix, PHD zinc finger, HORMA-binding closure motif and an extended carboxyterminal disorder tail).

### File S1

Text files (.hhr) with the outputs of HMM-vs-HMM profile searches (HHsearch) of full-length KKT16 and KKT17,18 and its separate domains (ARM, PH and coiled coil) against the combined PfamA and pdb70 HHsuite3 databases (see Data & Methods).

### File S2

Text files containing sequences of the eukaryotic SYCP2, SYCP3 and HOP1-like orthologues used in this study (see Table S2 for presence/absence profiles).

### File S3

Text files and visualisation of profile HMMs: (1) hmm3.1 profiles used for iterative HMM searches, (2) annotated .hhm + alignment files (.a3m), (3) graphical overview of conservation and secondary structure of annotated .hhm file (.pdf), (4) underlying alignments (.fasta files; if available). See for scripts and explanation of output File S6.

### File S4

Session files pertaining to the phylogenetic analyses for the ARM+PH and coiled coil Figure 4: (1) newick/nexus tree files, (2) alignments used to infer the phylogenetic trees, (3) log files of the IQ-Tree analysis and the untrimmed alignments.

### File S5

Jalview session files of panels C of Figure 2 and Figure 3. To achieve the same conservation scoring colour coding use the following settings in Jalview (version 2.11.1.3, RRID: SCR_006459 [103]): ‘colour by annotation’ > Conservation > Above Threshold > Threshold is min/max > threshold 3.5.

### File S6

Relevant settings/description of software packages and example scripts used to annotate and visualise HMMs and the HHsearch output in Figure 1, Figure 2, Figure 3 and File S3. This uncompressed file contains the following: (1) example folder with executables and generated files, (2) a library of python and perl scripts, (3) visual guidance on how to interpret graphical overviews of HMM figures, (4) overview and manual on how to generate annotated HMM (.pdf). The scripts incorporate a modified version of the visualisation script ‘hhsearch-figgen’ (https://github.com/twemyss/hhsearch-figgen) developed by T.A.W.

## Notes

https://doi.org/10.6084/m9.figshare.13725787

### Competing Interest Statement

The authors have declared no competing interest.

