## Supplementary figures and images for "Repurposing of Synaptonemal Complex Proteins for Kinetochores in Kinetoplastida"

### 3_Graphical_overview_HHM_explained.pdf

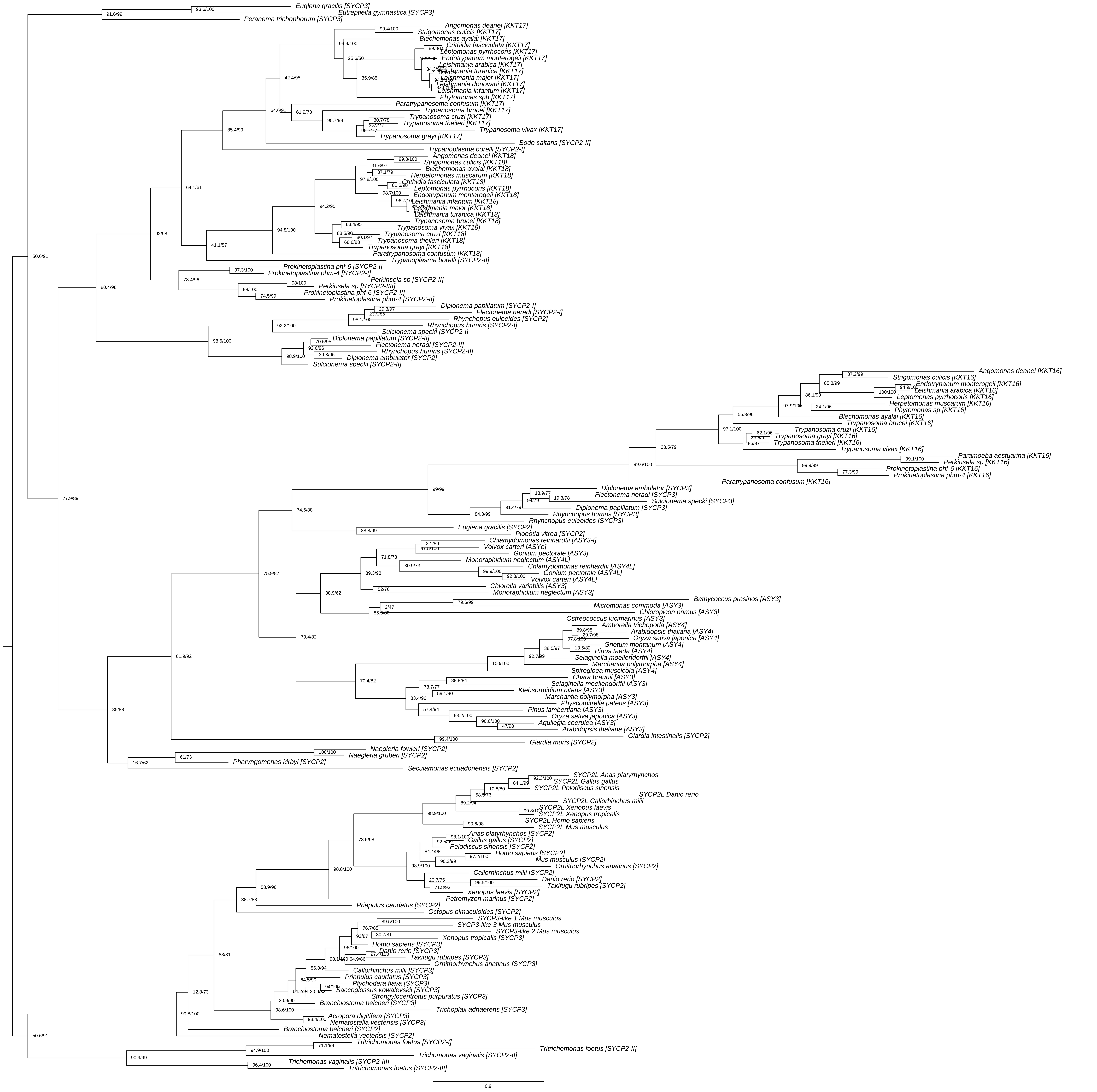

Supplement: File S4 [file 430040_file05.zip › File_S4/File_S4_coiled_coil_70.uncollapsed.treefile [UFBOOT|SH-aLRT].pdf]

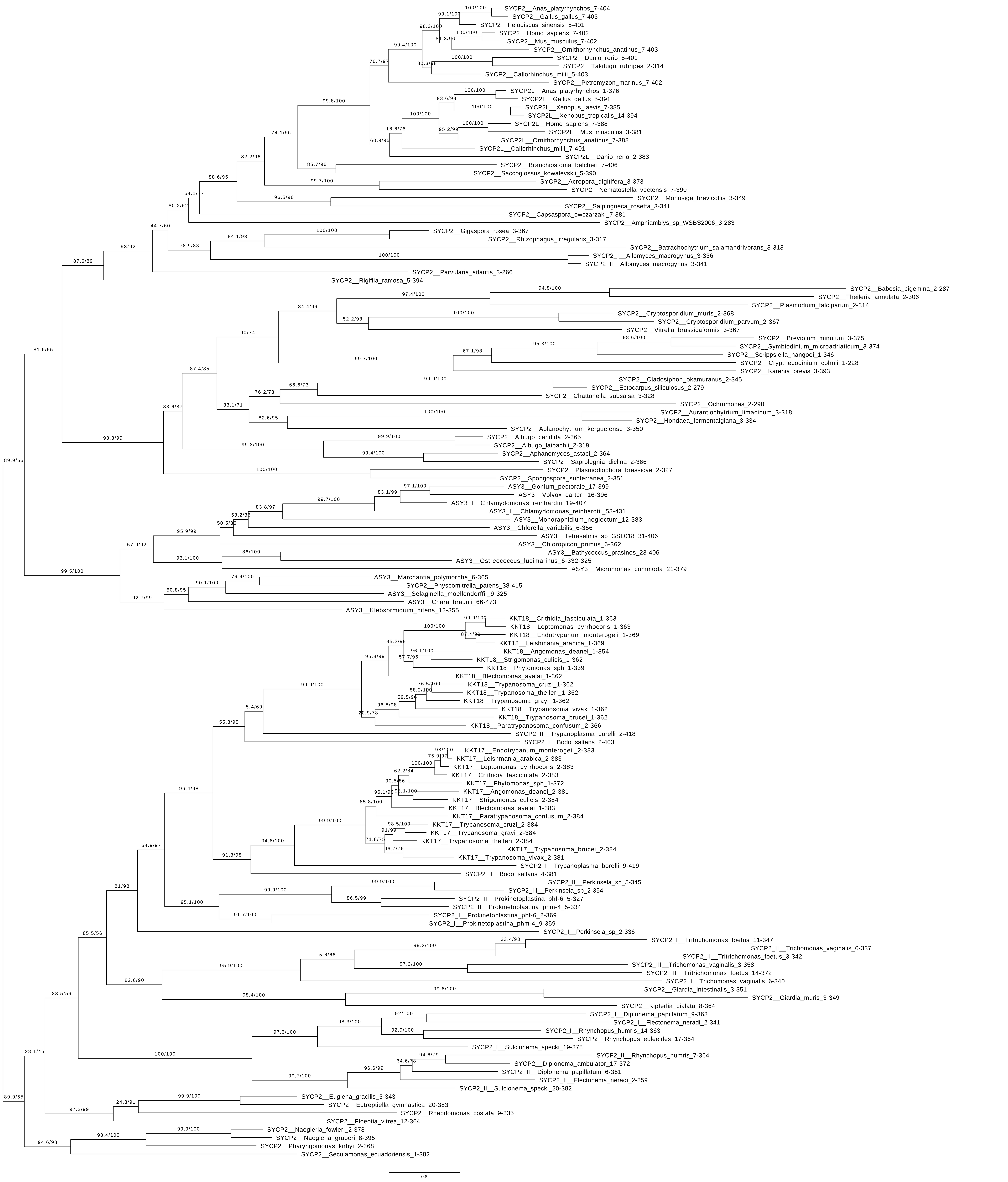

Supplement: File S4 [file 430040_file05.zip › File_S4/File_S4_ARM_PH_30.uncollapsed.treefile [UFBOOT|SH-aLRT].pdf]

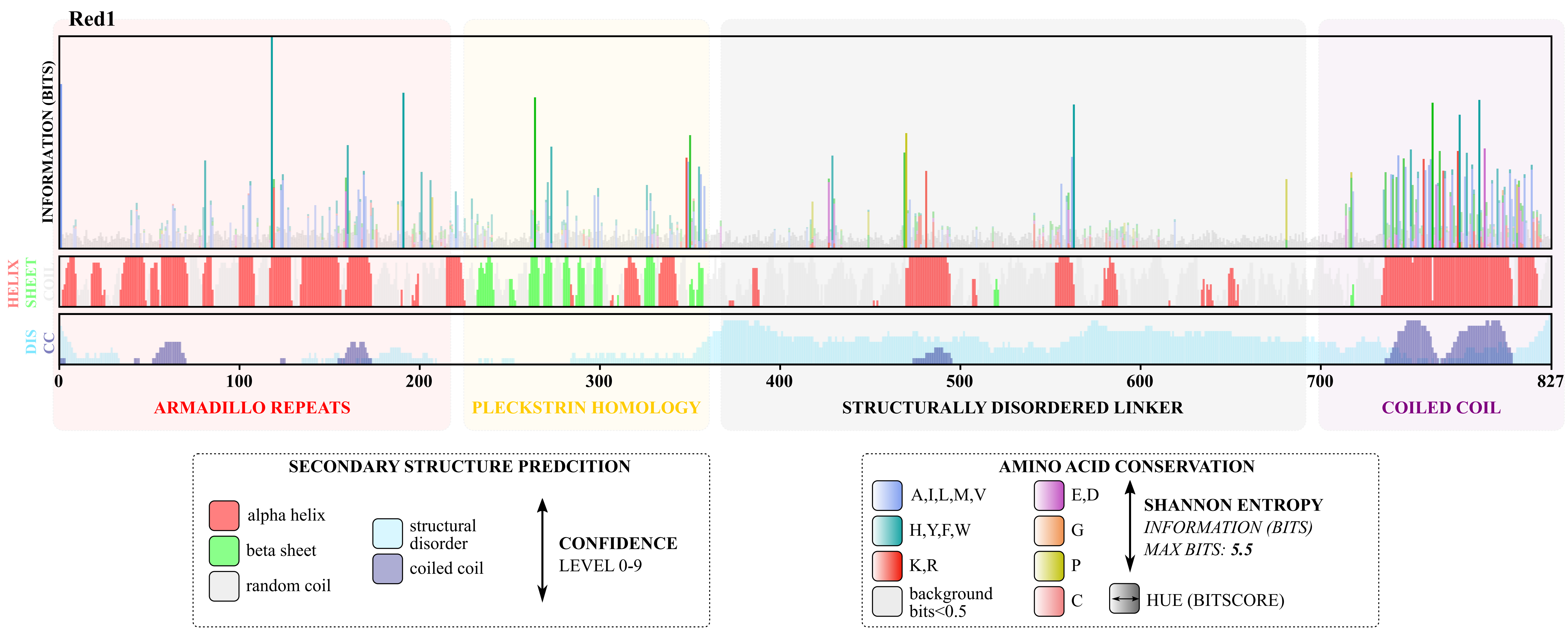

### ASY3_Archaeplastida.pdf

ASY3_Archaeplastida

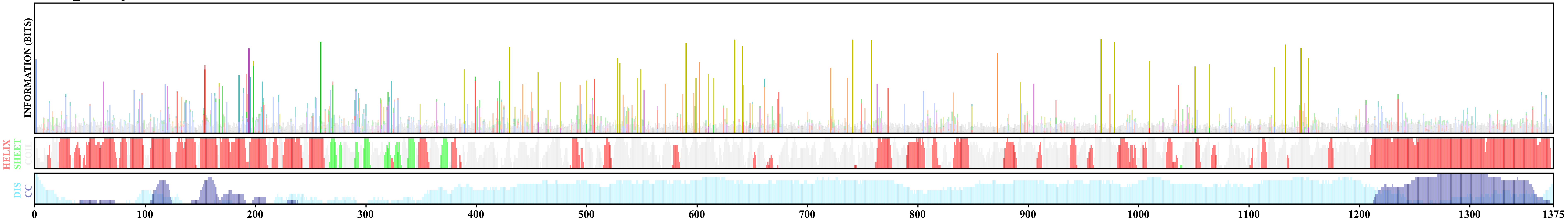

### ASY3_Spermatophyta_loss_ARM_PH.pdf

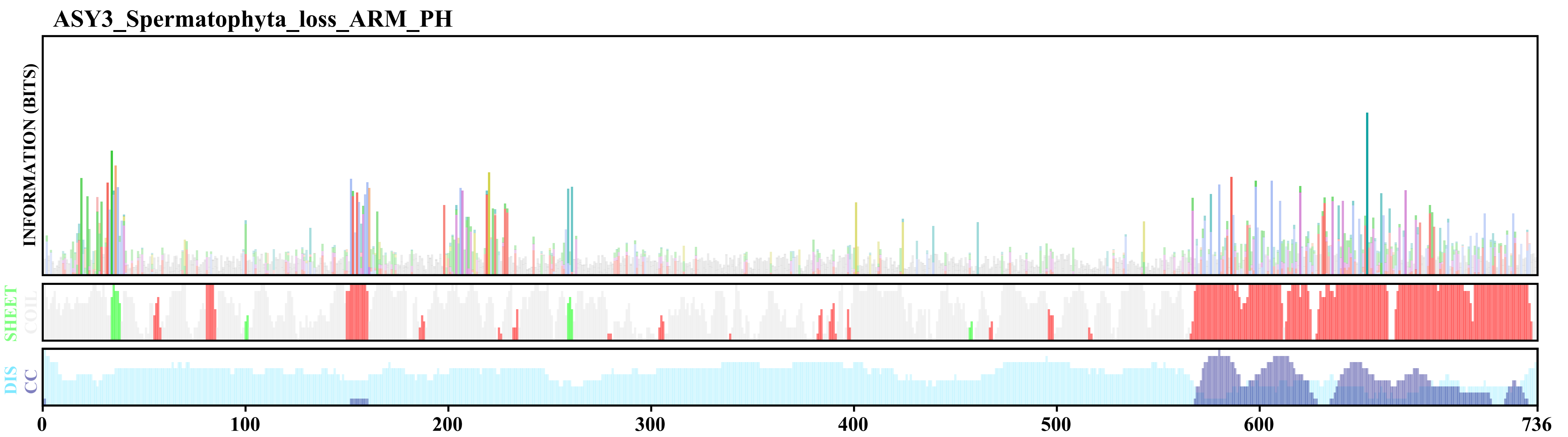

### ASY4_Streptophyta.pdf

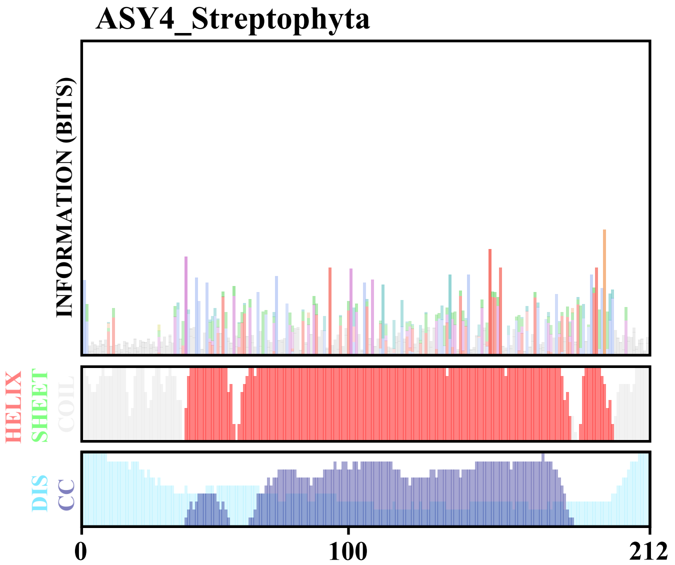

### ASY4L_Chlorophyceae.pdf

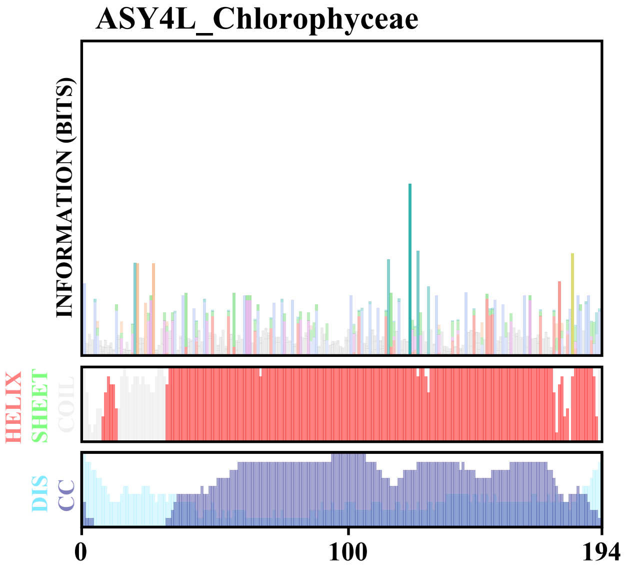

### Figure S1

#### FIGURE S1

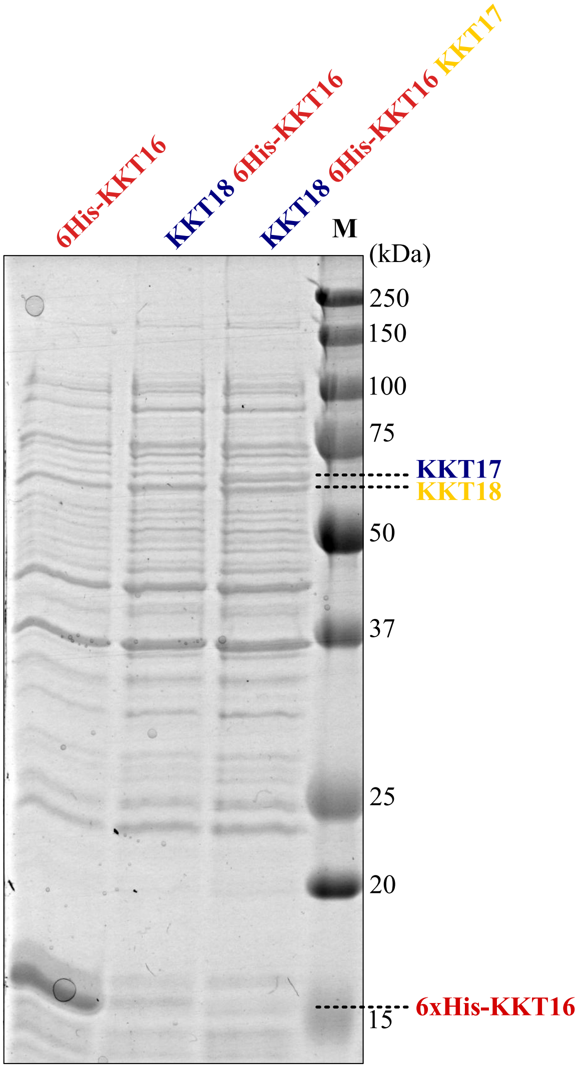


Tromer *et al.*2021

### Figure_2_KKT17_18_consensus.pdf

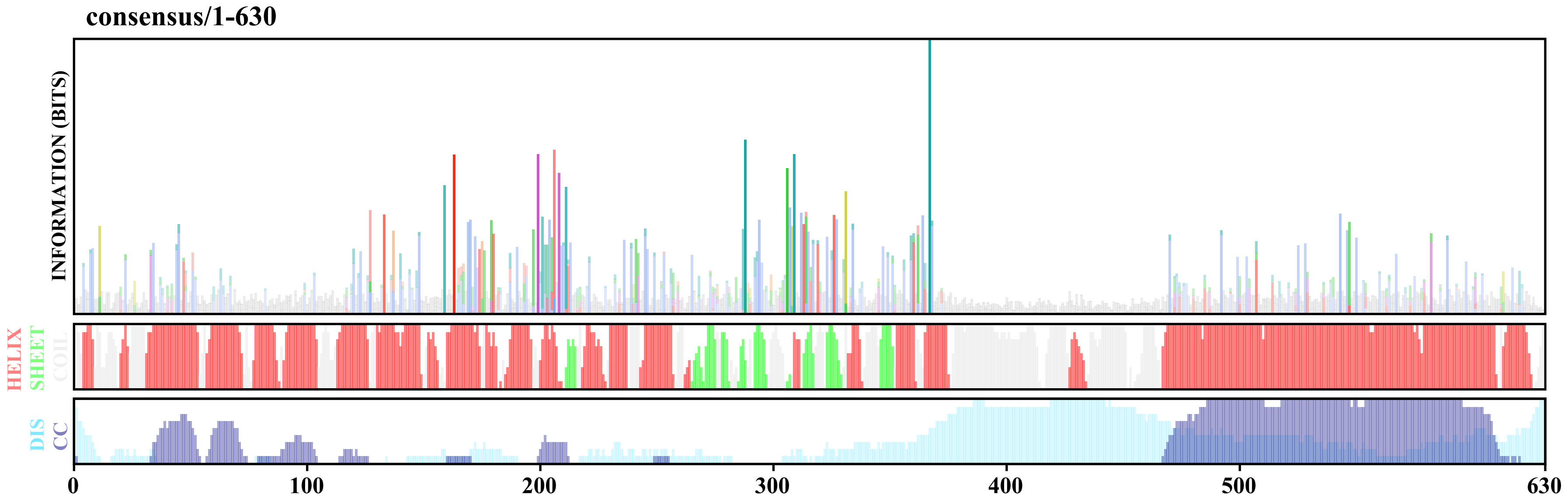

### Figure_3_KKT16_consensus.pdf

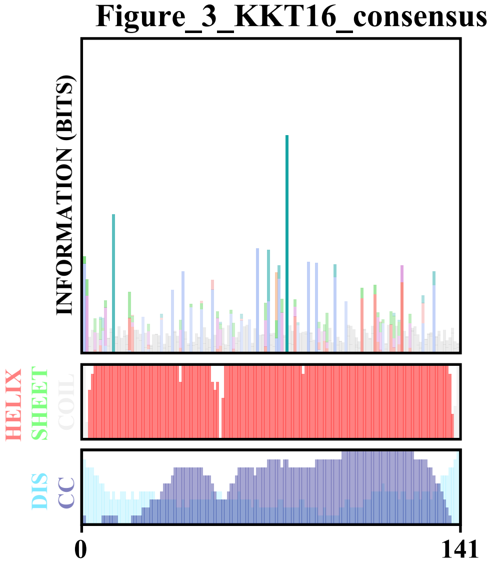

### File_S4_ARM_PH.pdf

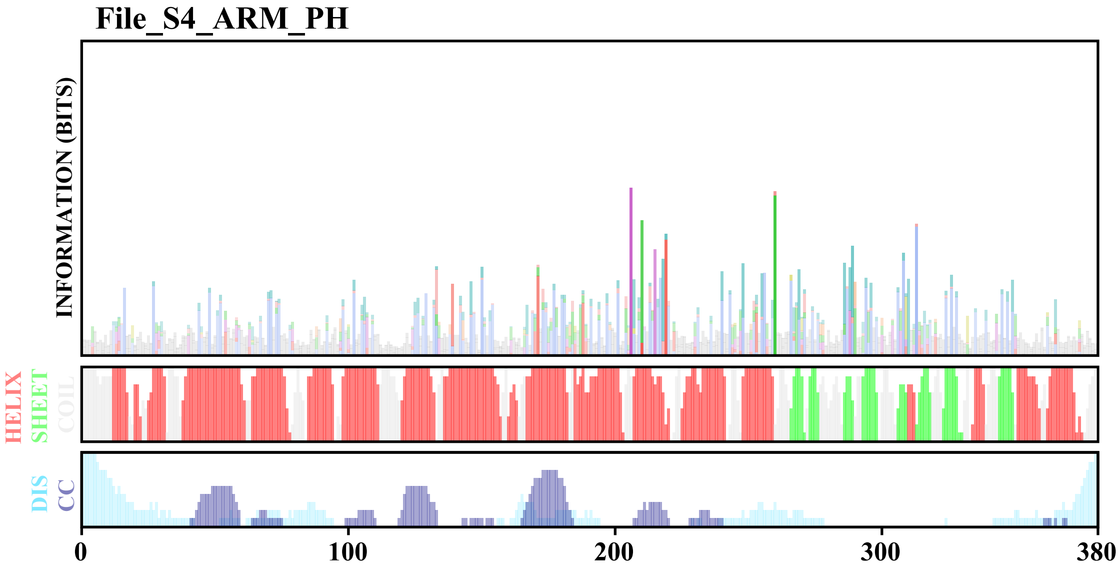

### File_S4_ARM_PH_30.pdf

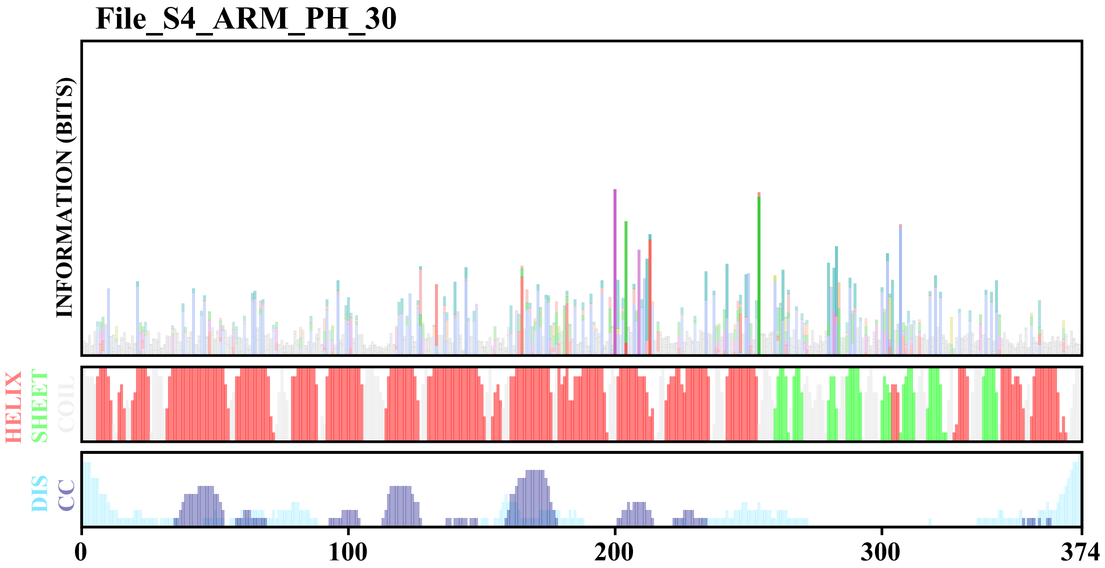

### File_S4_coiled_coil.pdf

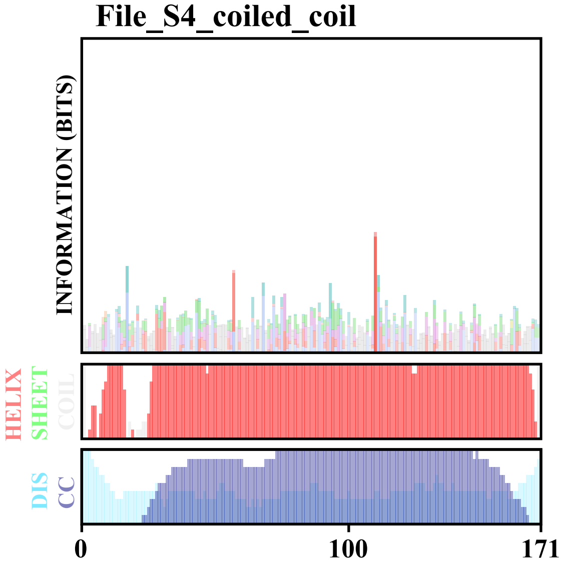

### File_S4_coiled_coil_70.pdf

#### File_S4_coiled_coil_70

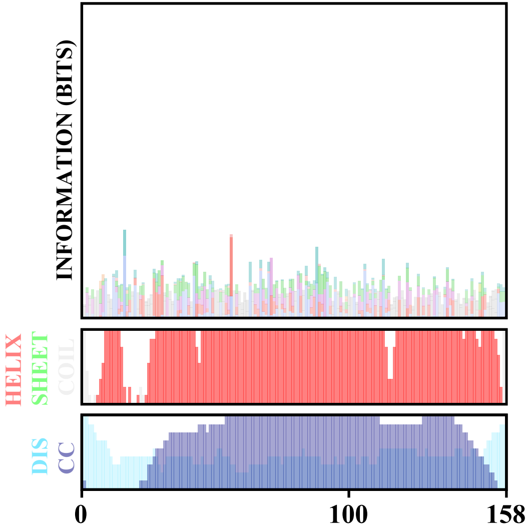

### KKT16_Kinetoplastida.pdf

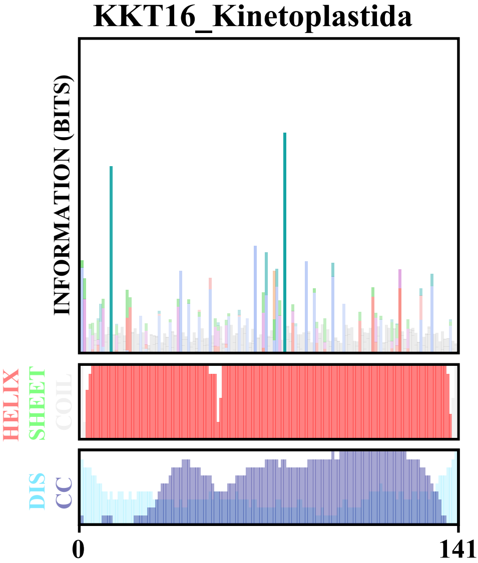

### KKT17_Metakinetoplastina.pdf

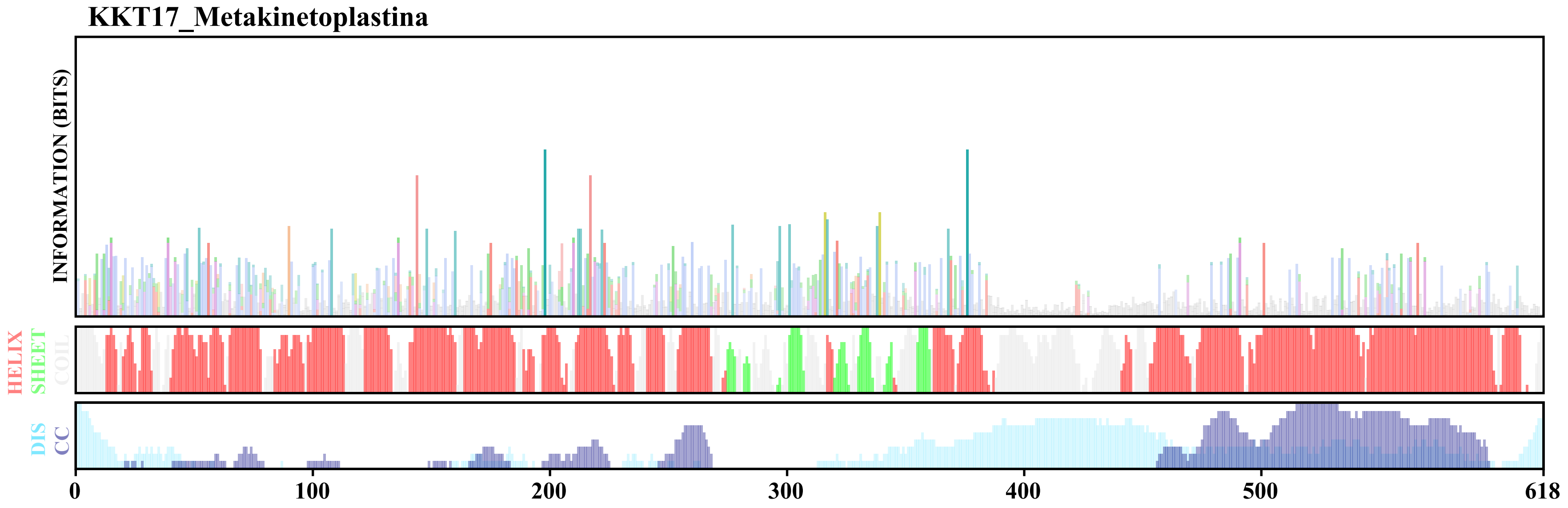

### KKT18_Metakinetoplastina.pdf

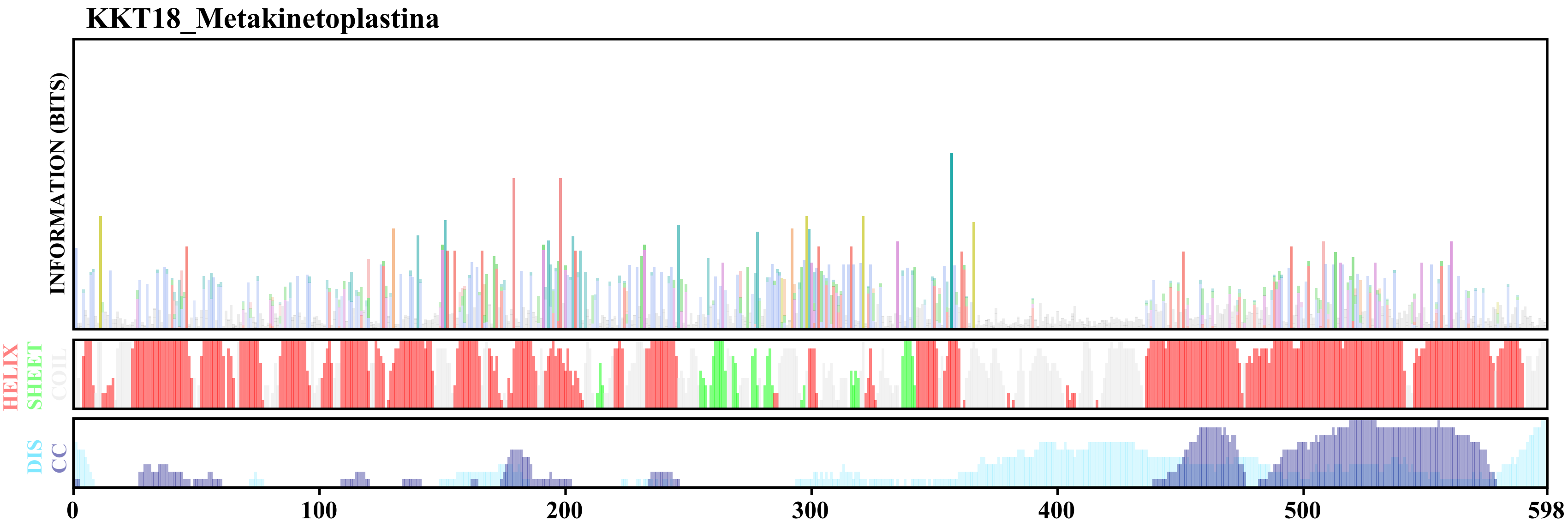

### Rec10.pdf

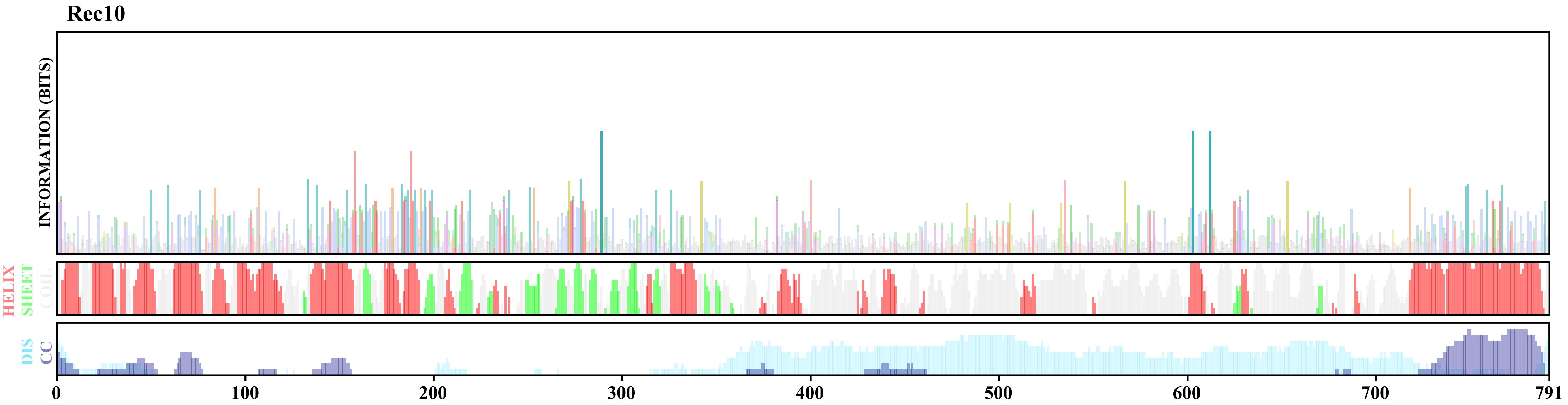

### Rec10_Red1_SYCP2_Fungi.pdf

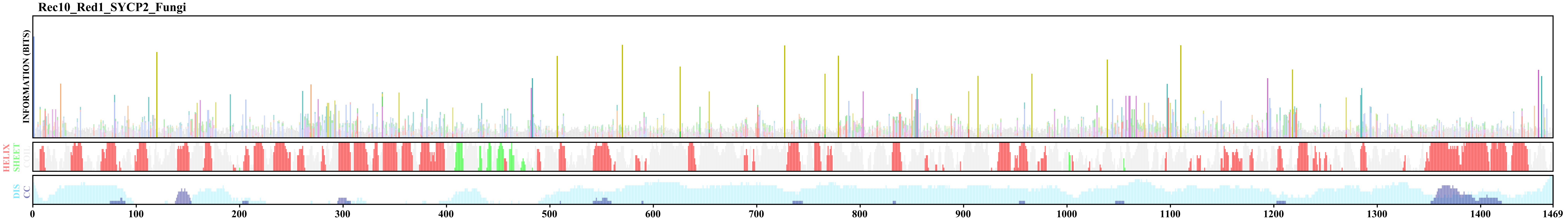

### Rec27.pdf

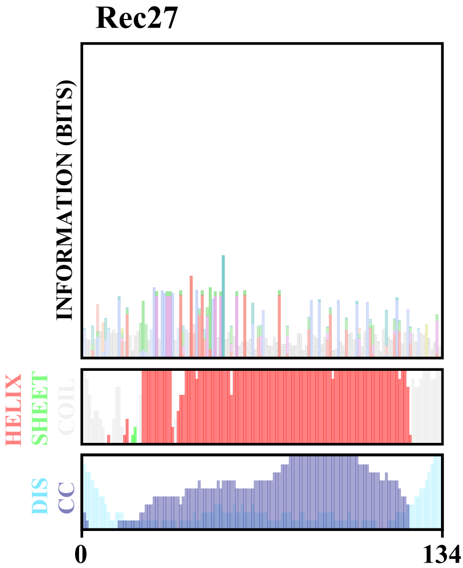

### Rec27_SYCP3_fungi.pdf

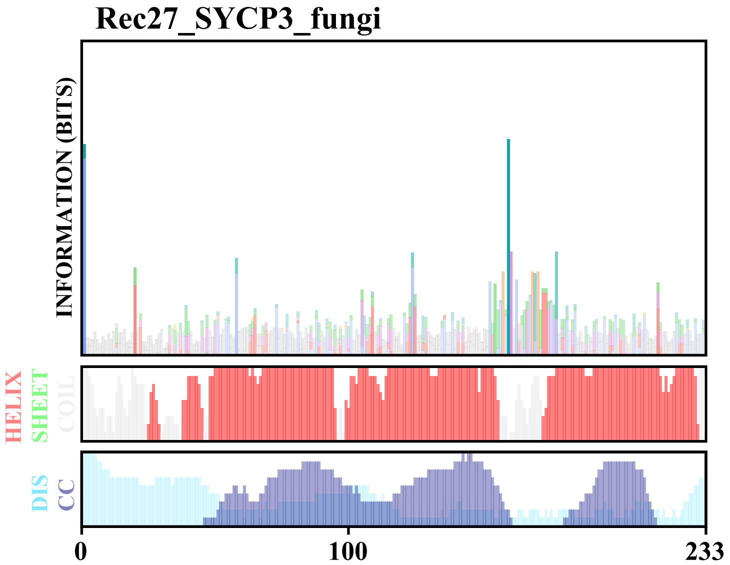

### Red1.pdf

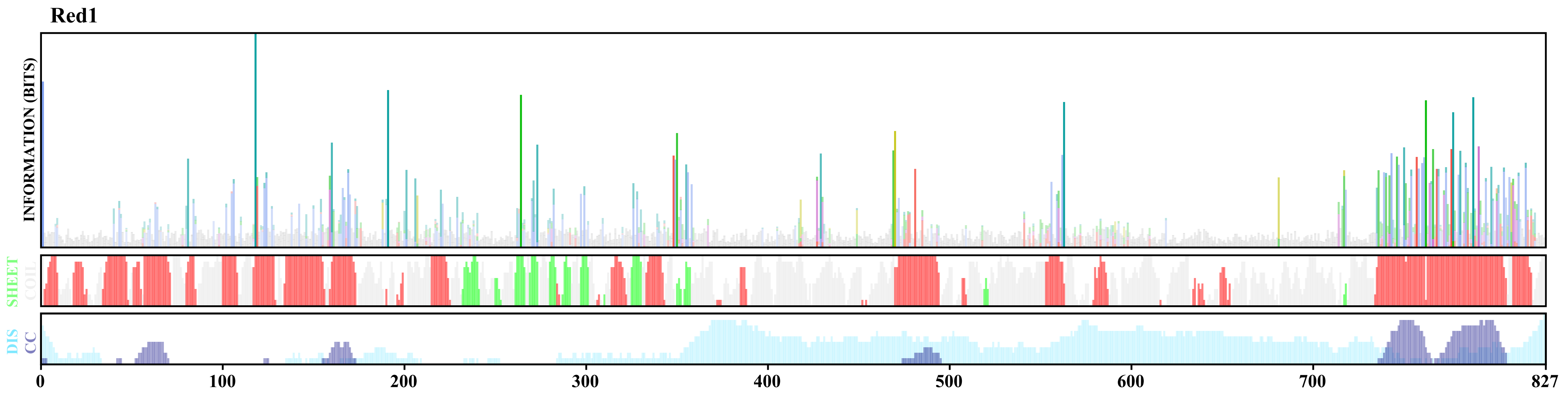

### SYCP2_Apicomplexa.pdf

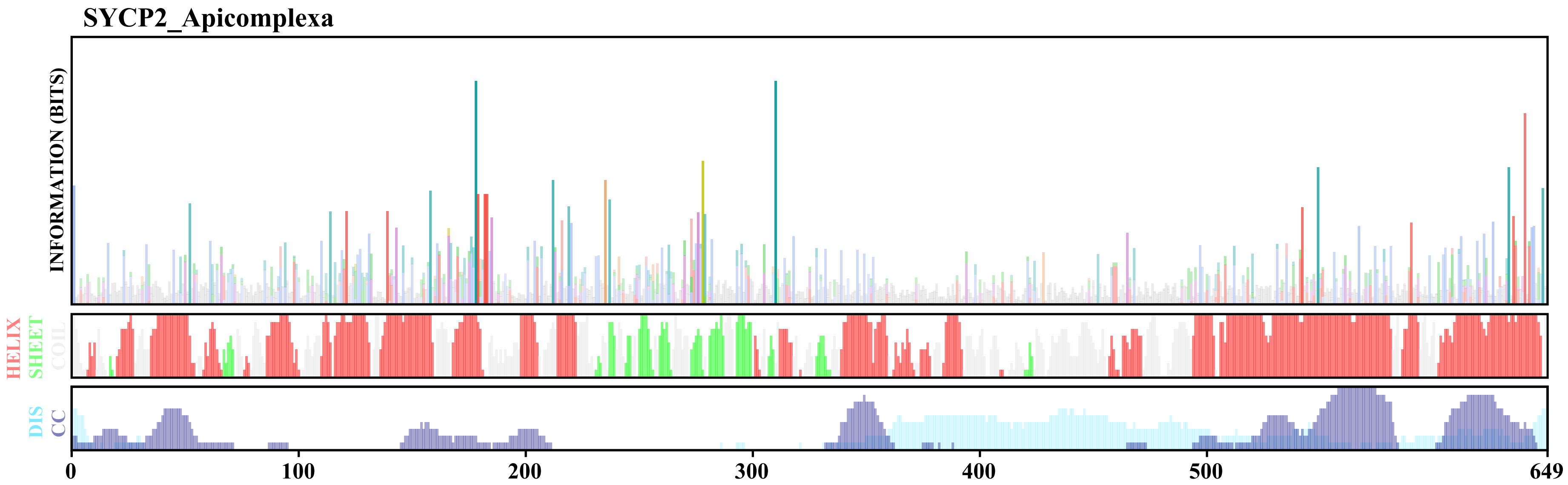

### SYCP2_Arthropoda.pdf

##### SYCP2_Arthropoda

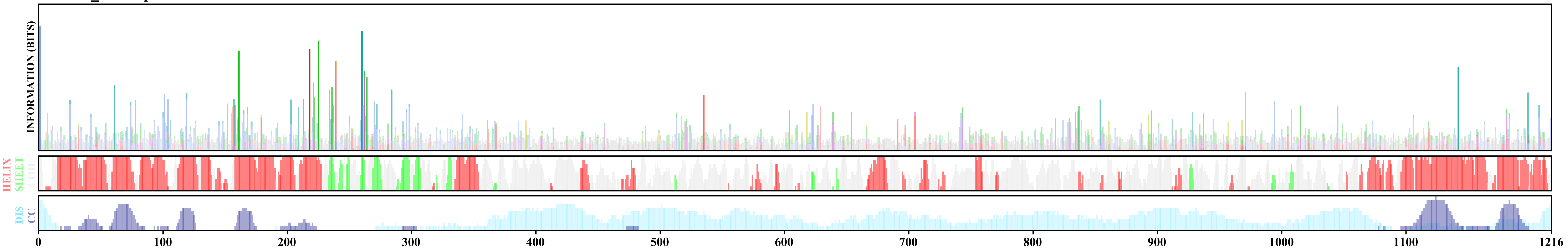

### SYCP2_Dinoflagellates_amino_terminus.pdf

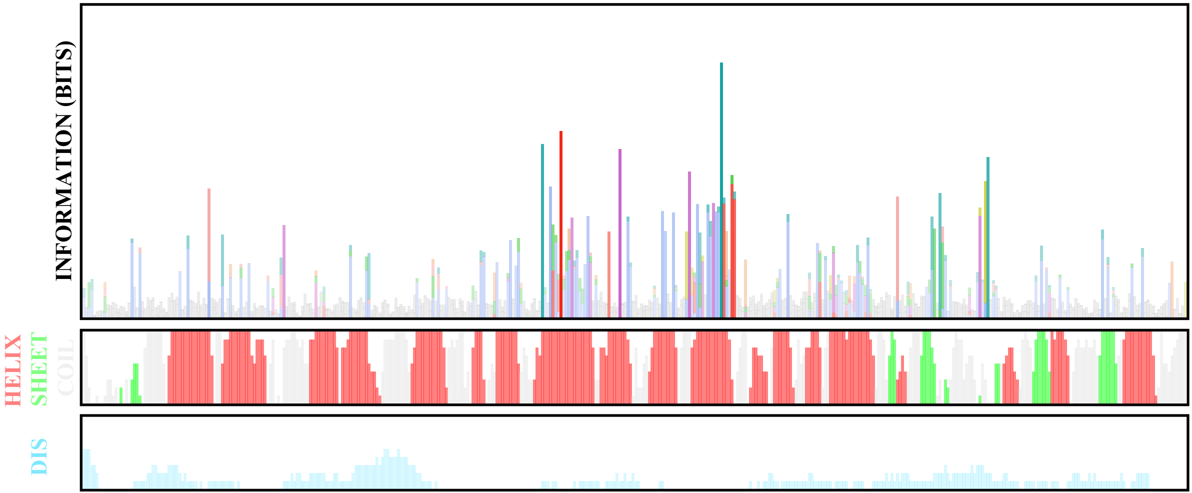

### SYCP2_Diplonemids_1.pdf

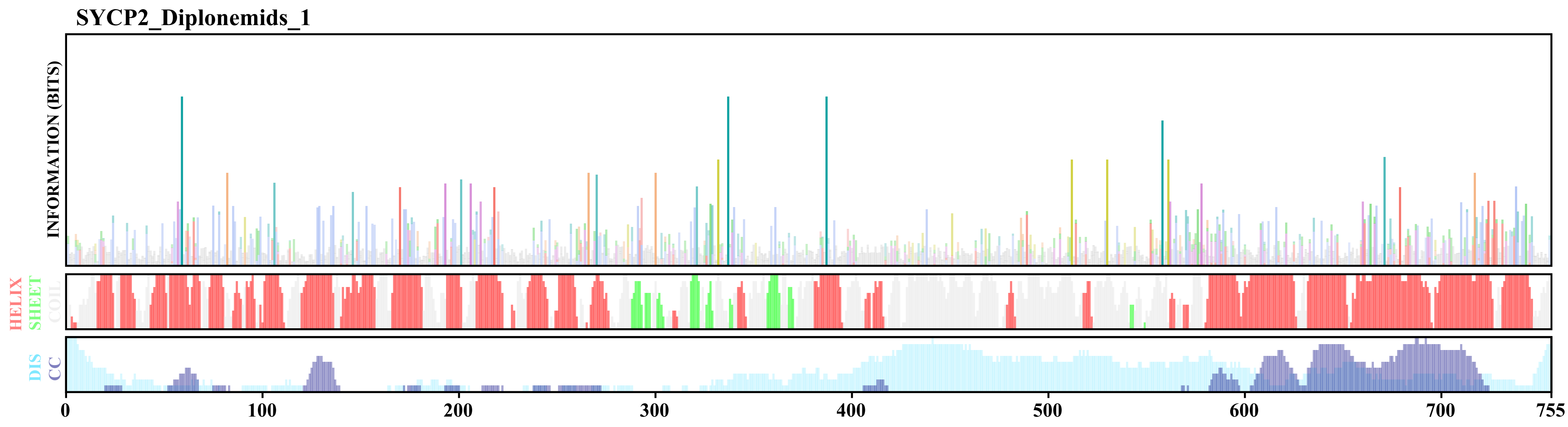

### SYCP2_Diplonemids_2.pdf

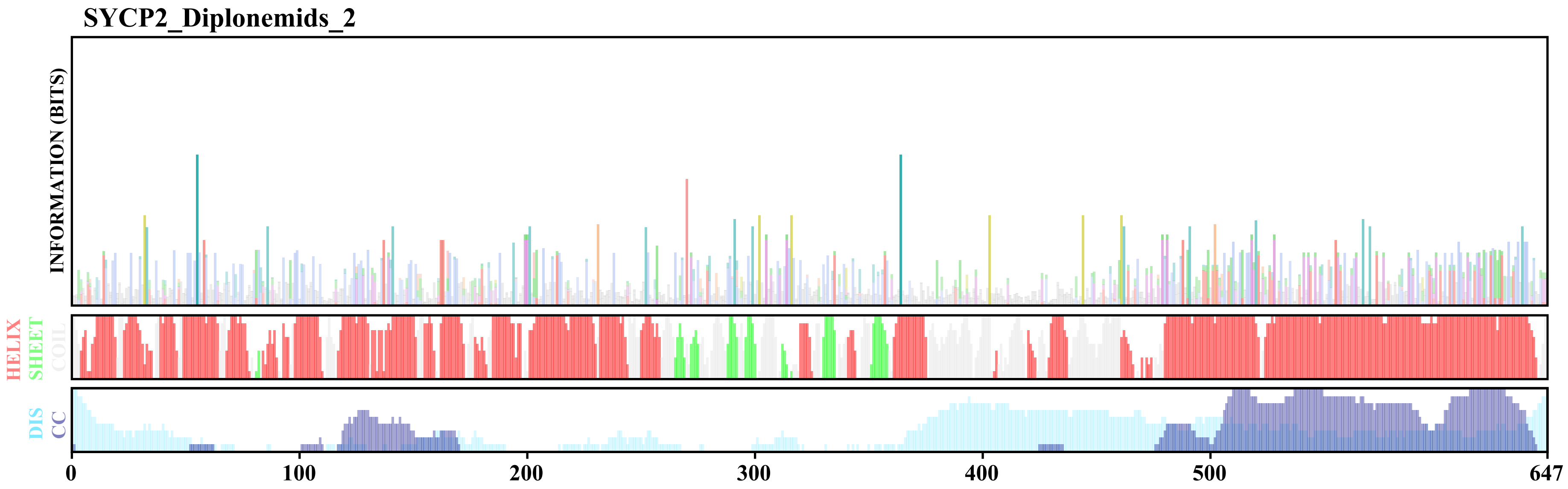

### SYCP2_Discoba.pdf

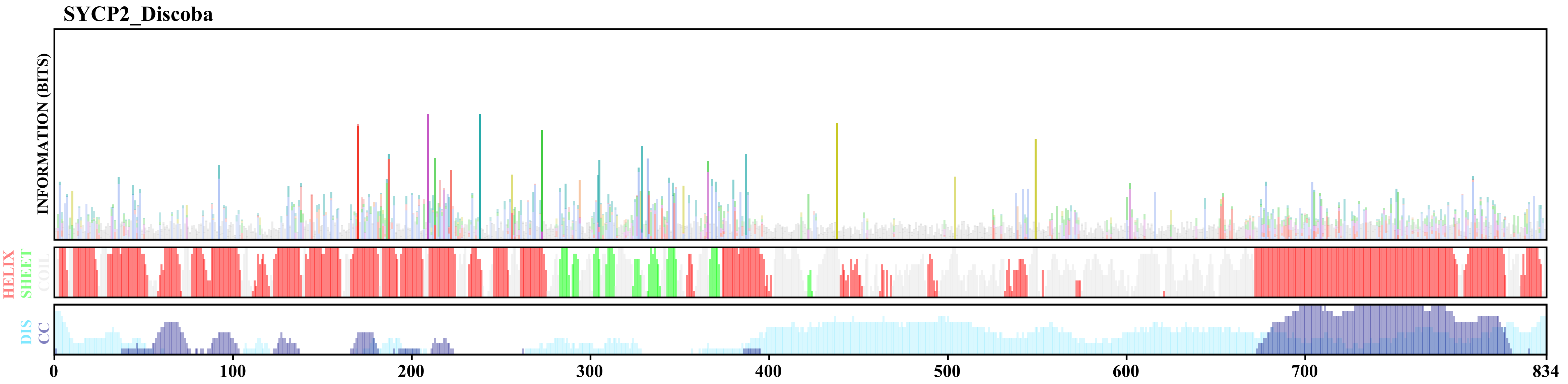

### SYCP2_DIscoba_2.pdf

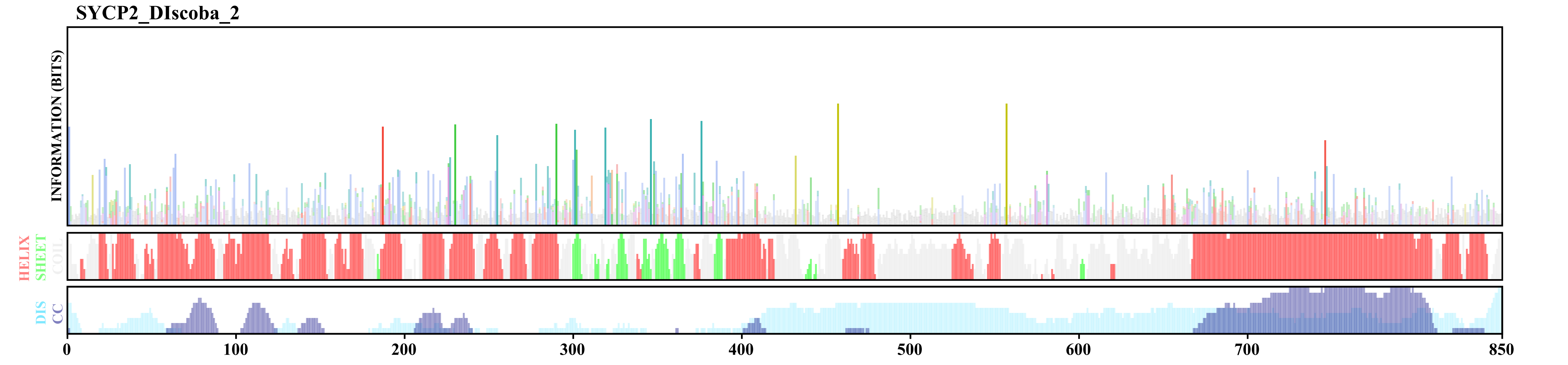

### SYCP2_Euglenids.pdf

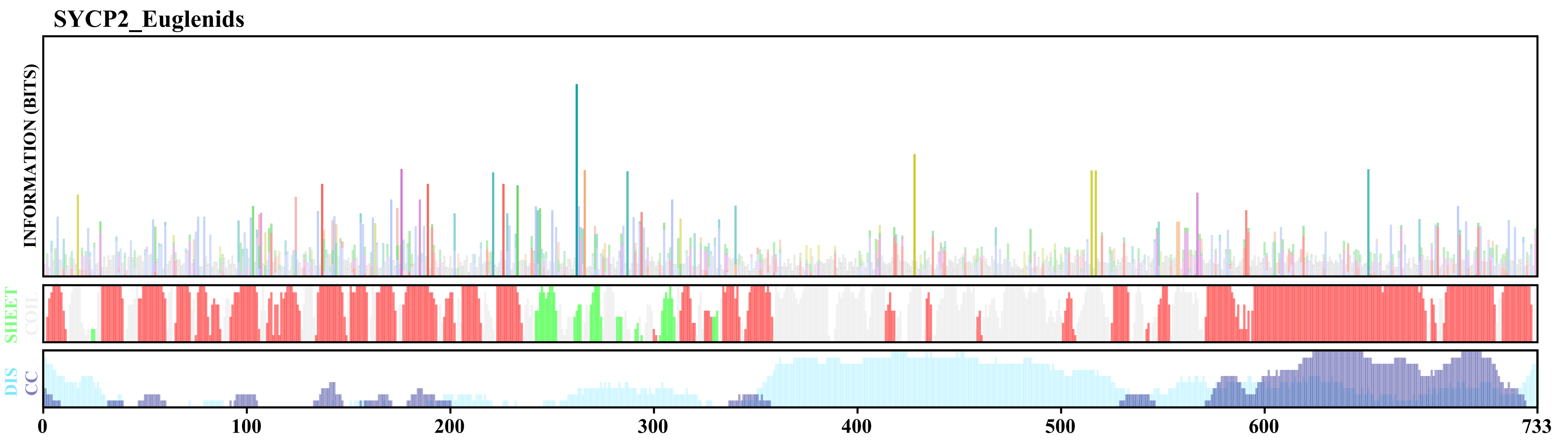

### SYCP2_Fornicata.pdf

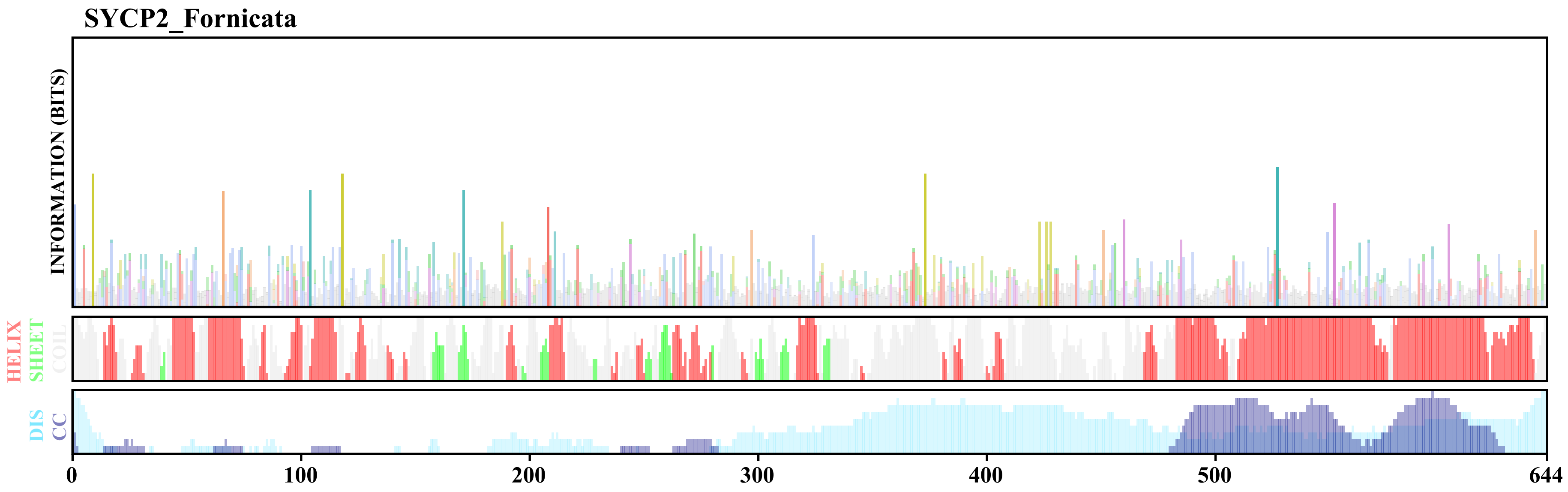

### SYCP2_Fungi.pdf

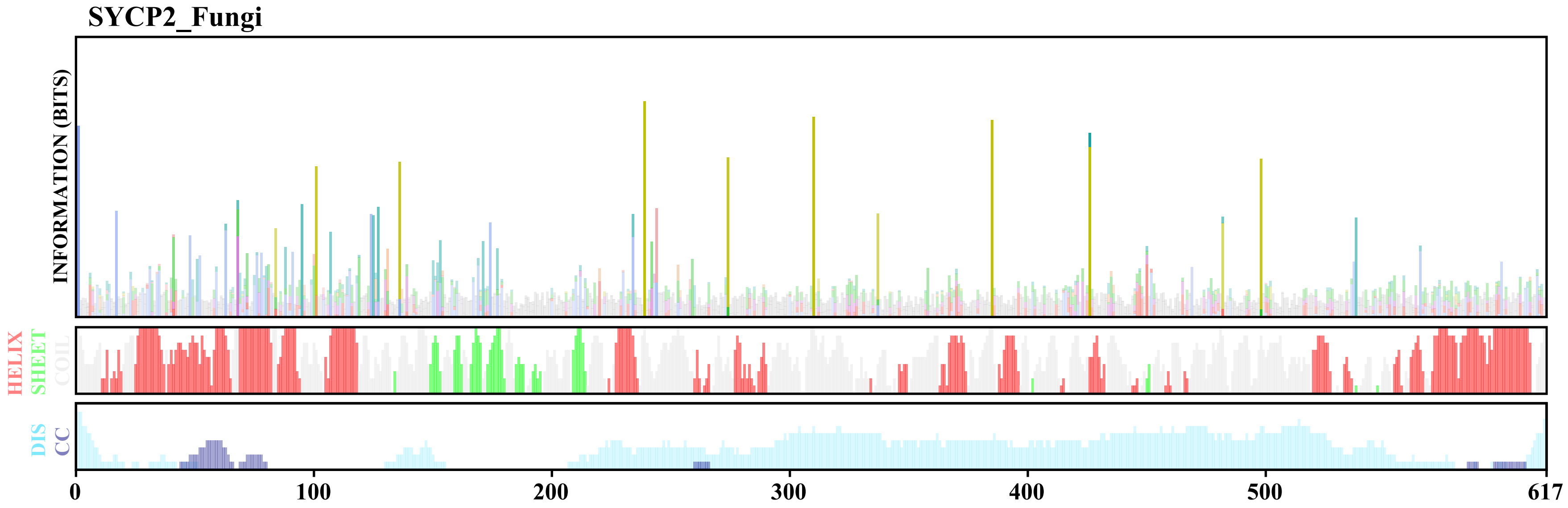

### SYCP2_Fusarium_loss_ARM_PH.pdf

SYCP2_Fusarium_loss_ARM_PH

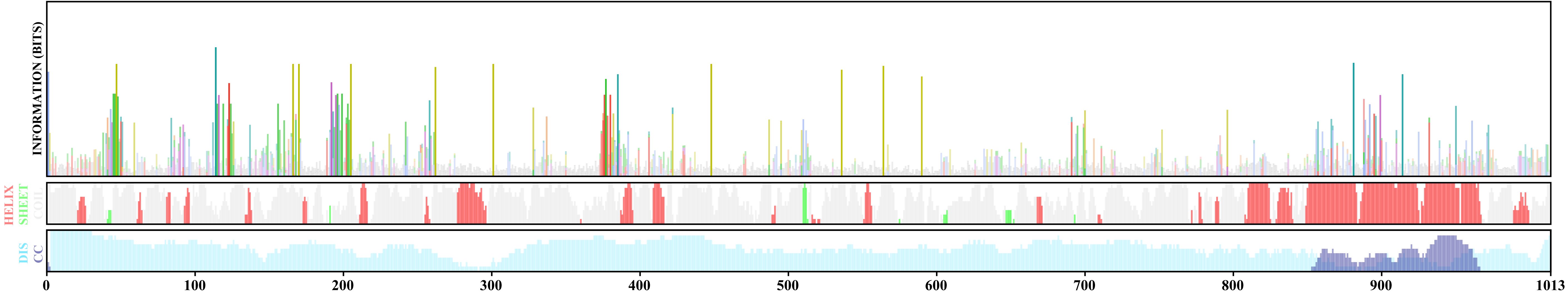

### SYCP2_Labyrinthulomycetes.pdf

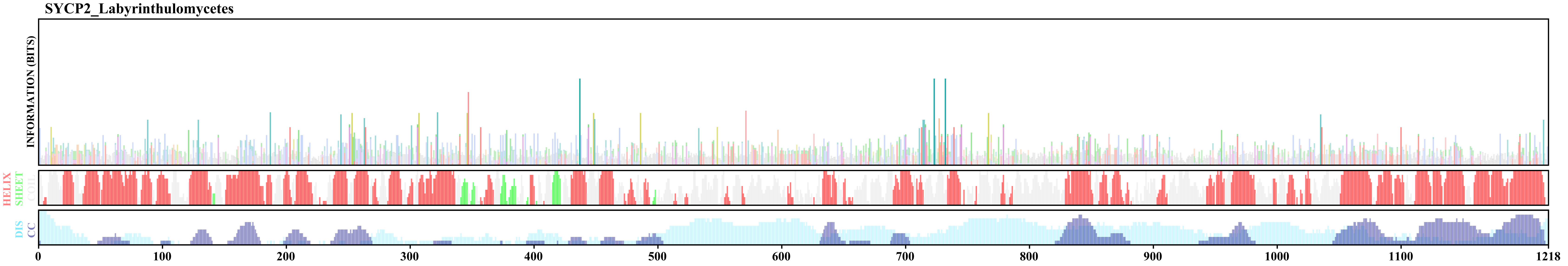

### SYCP2_like_Eurotiomycetes_and_other_ascomycetes.pdf

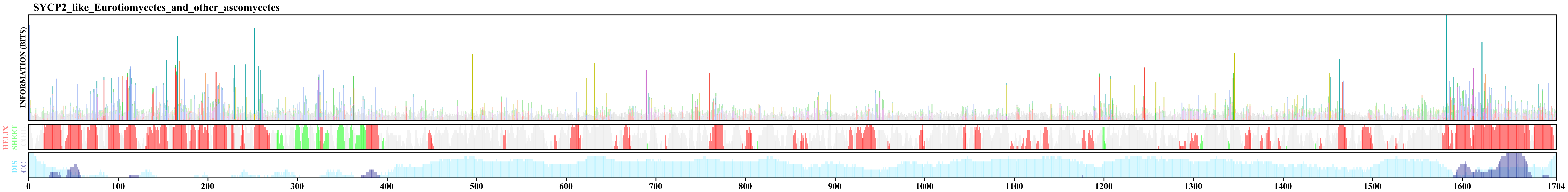

### SYCP2_Metamonada.pdf

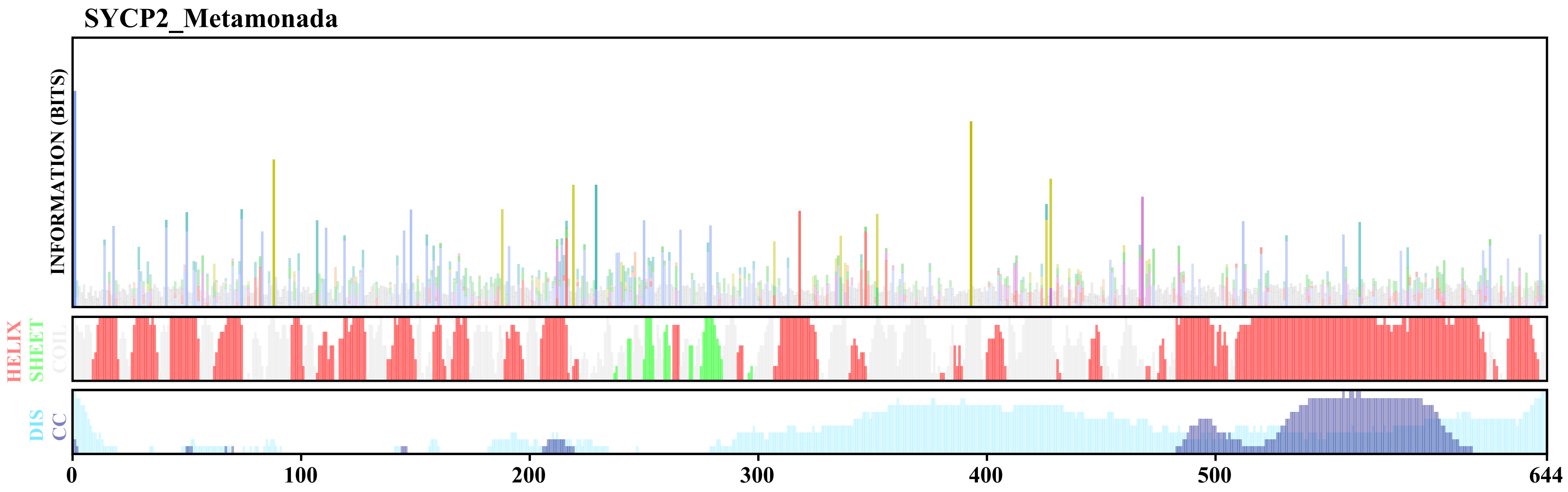

### SYCP2_Metazoa.pdf

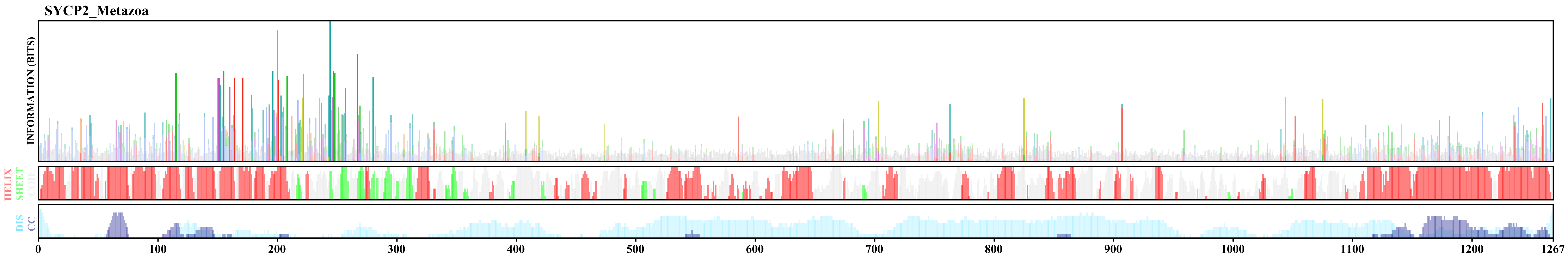

### SYCP2_Oomycetes.pdf

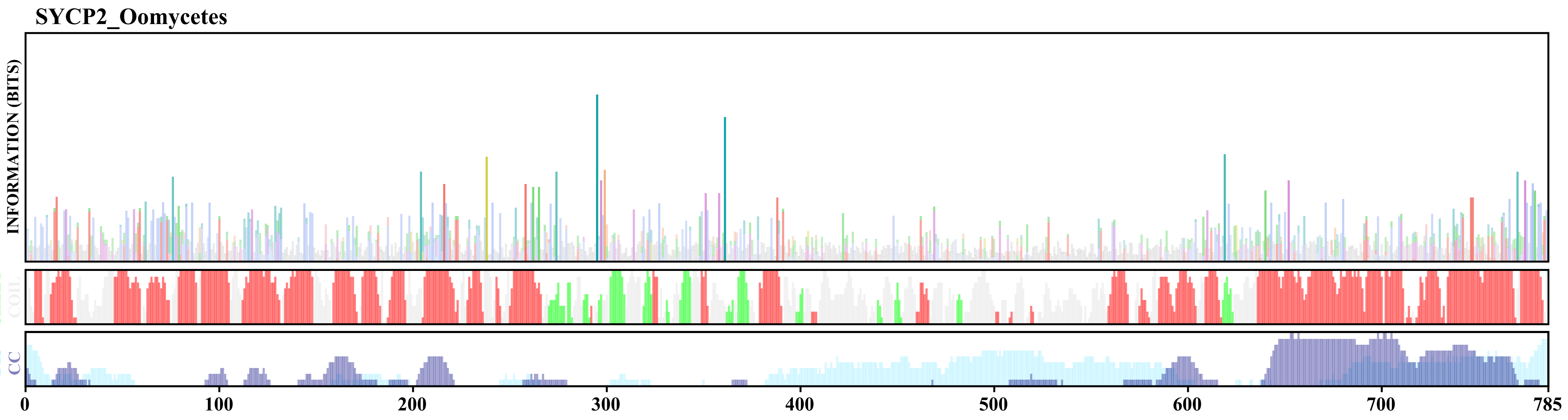

### SYCP2_Opimoda.pdf

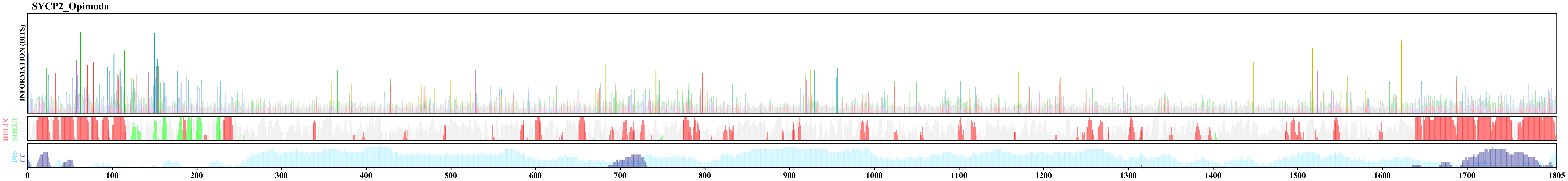

### SYCP2_Parabasalia_II.pdf

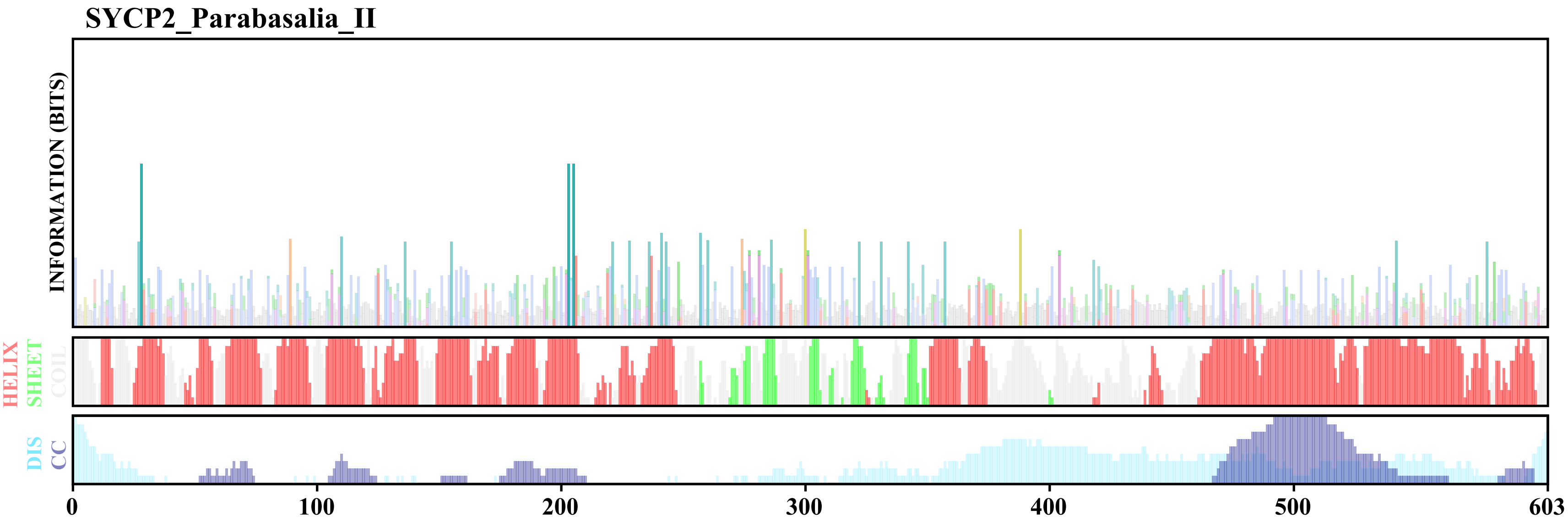

### SYCP2_Parabasalia_III.pdf

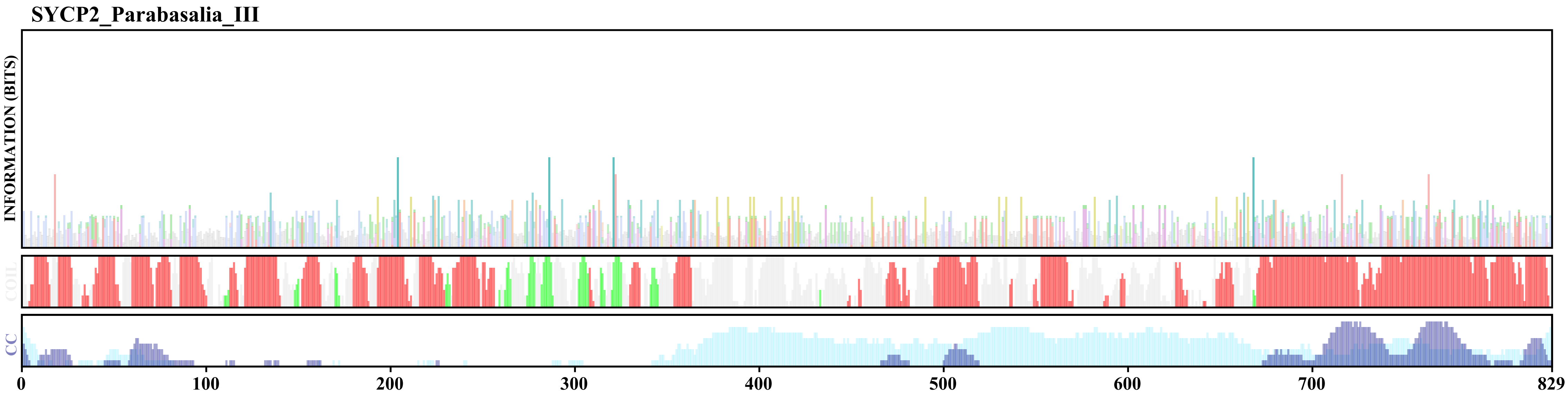

### SYCP2_Perkinsela_I.pdf

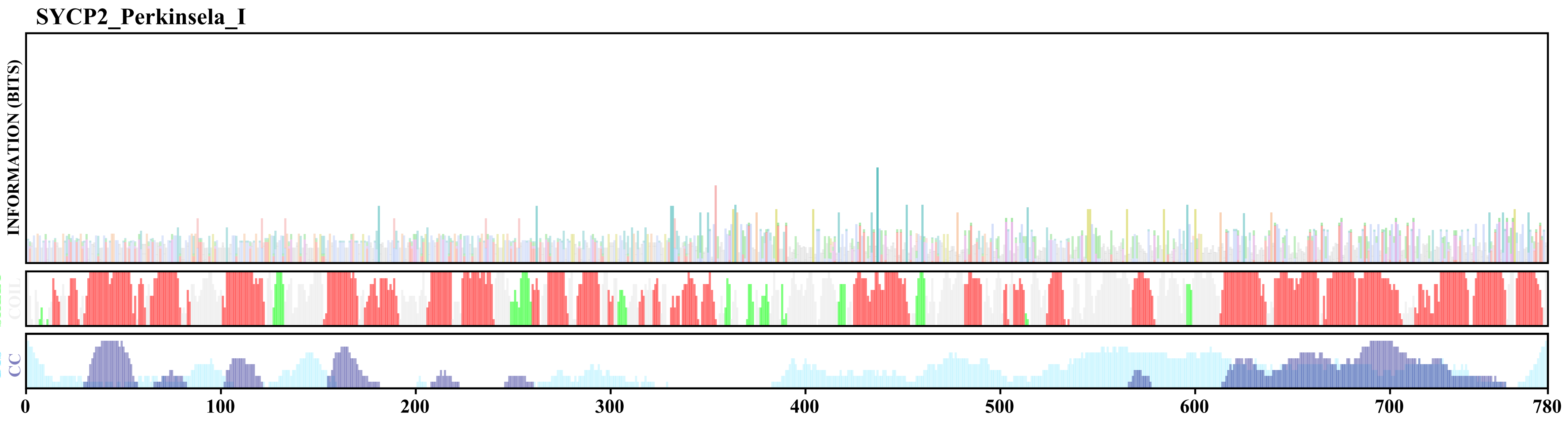

### SYCP2_Perkinsela_II_III.pdf

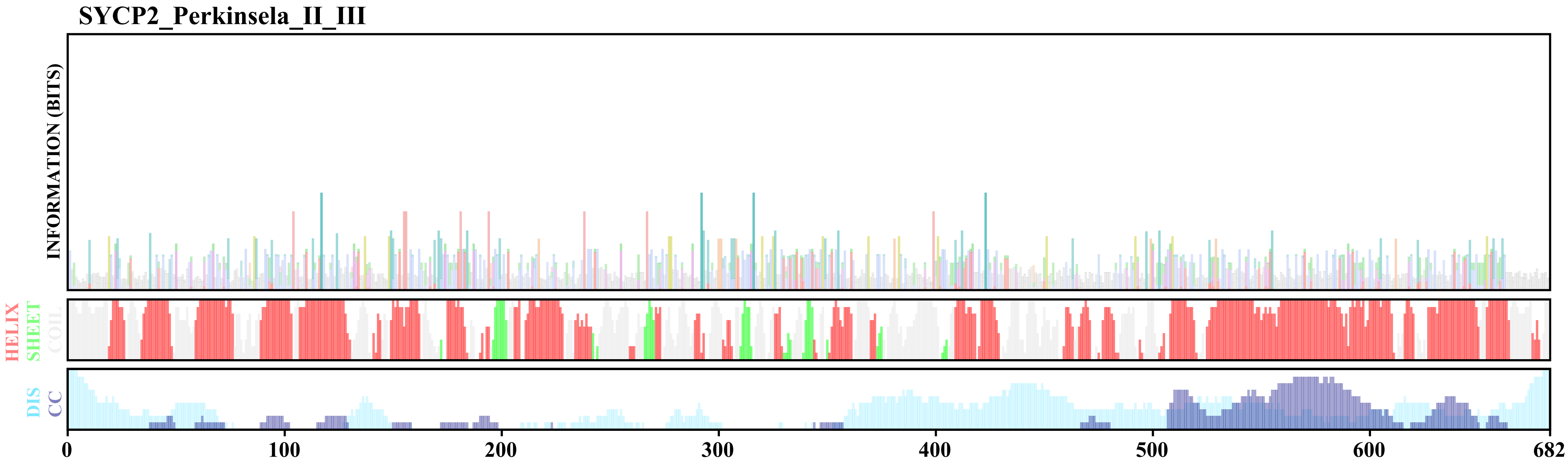

### SYCP2_Pezizomycota.pdf

SYCP2_Pezizomycota

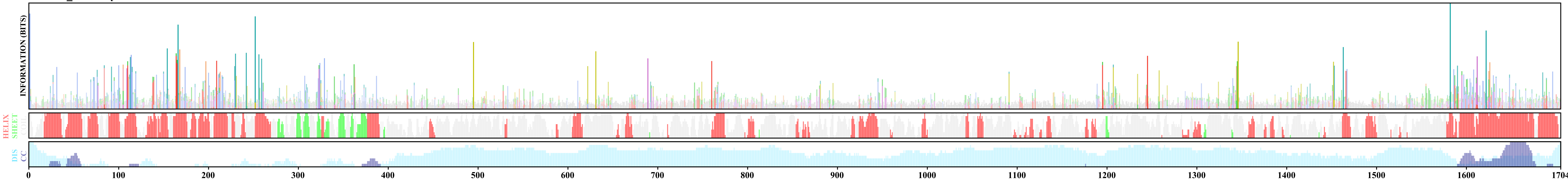

### SYCP2_Prokinetoplastina_I.pdf

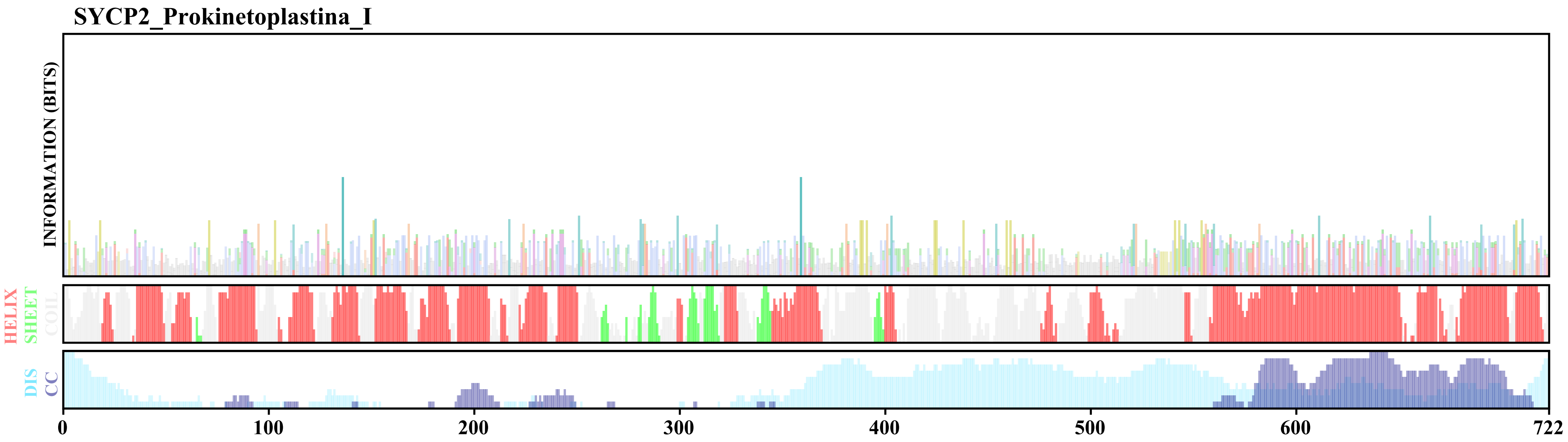

### SYCP2_Prokinetoplastina_II.pdf

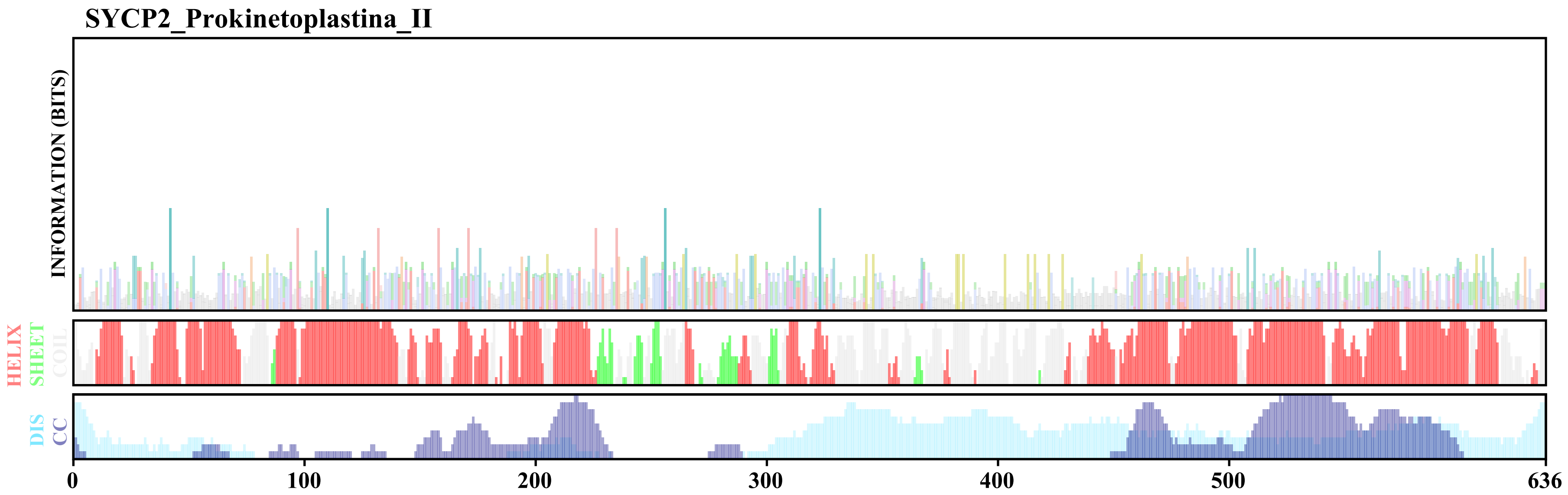

### SYCP2_Rhizaria.pdf

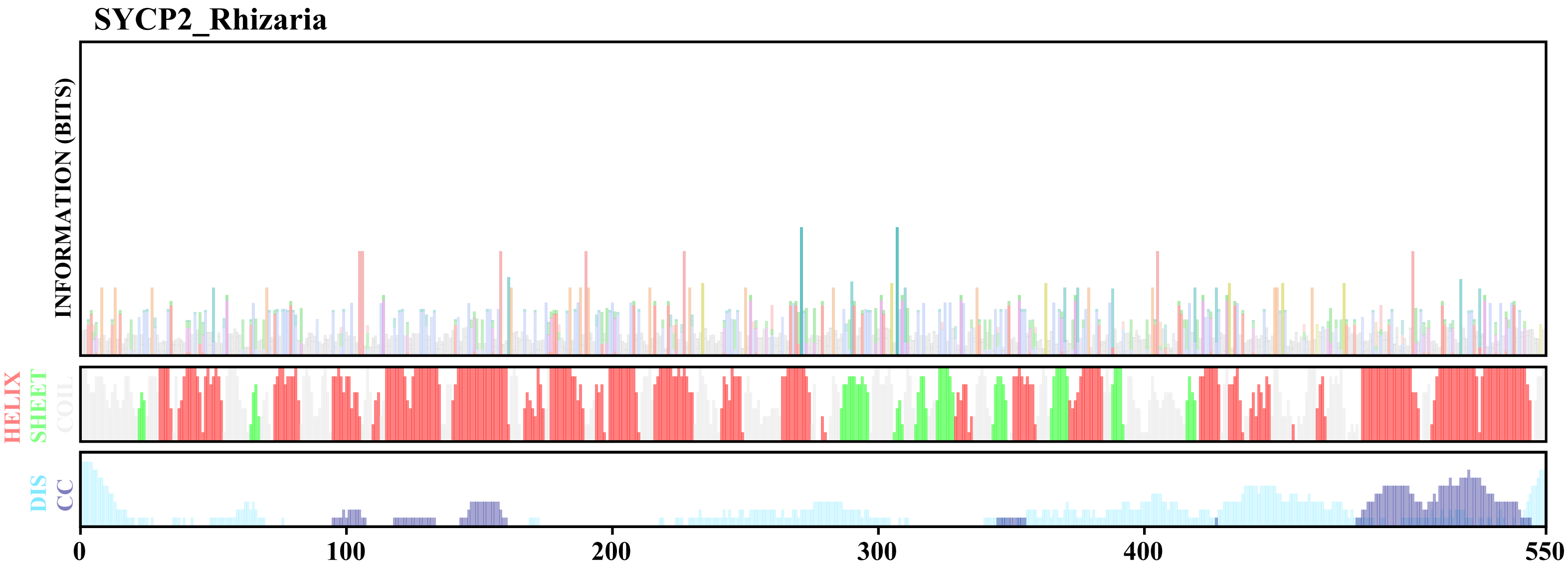

### SYCP2_SAR.pdf

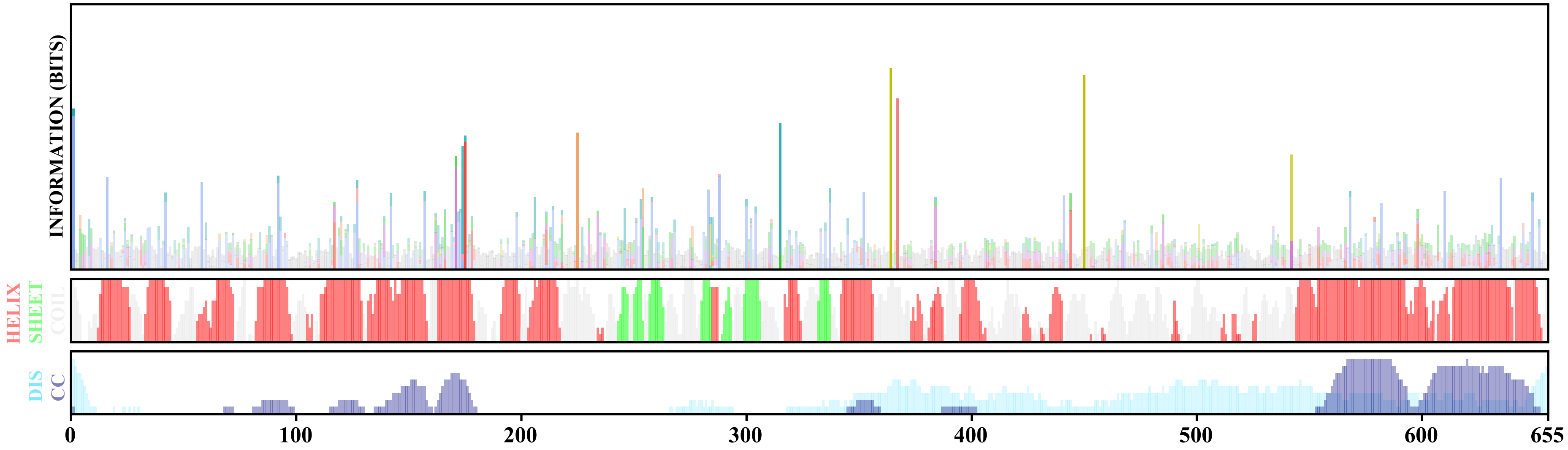

### SYCP2_Trichomonas_vaginalis_I.pdf

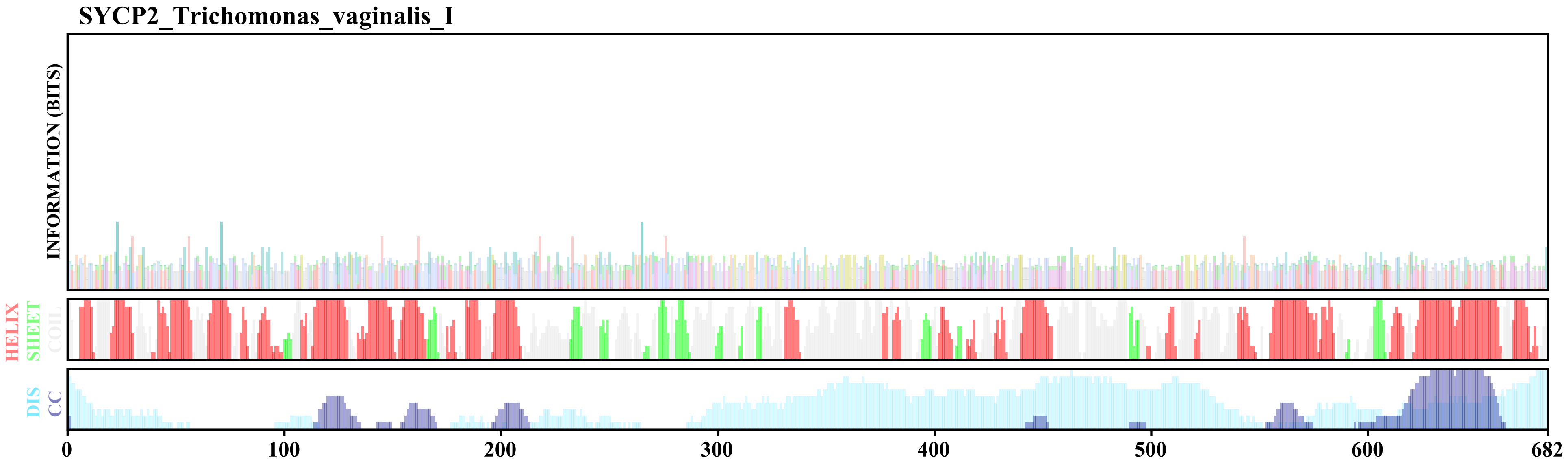

### SYCP2_Vertebrata.pdf

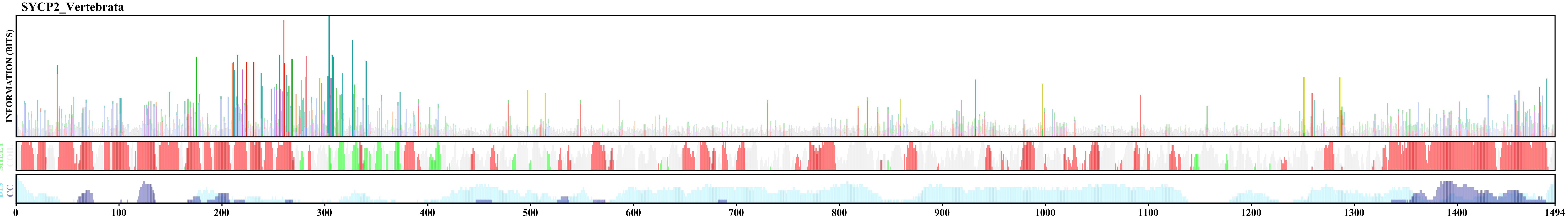

### SYCP2L_Vertebrata.pdf

SYCP2L_Vertebrata

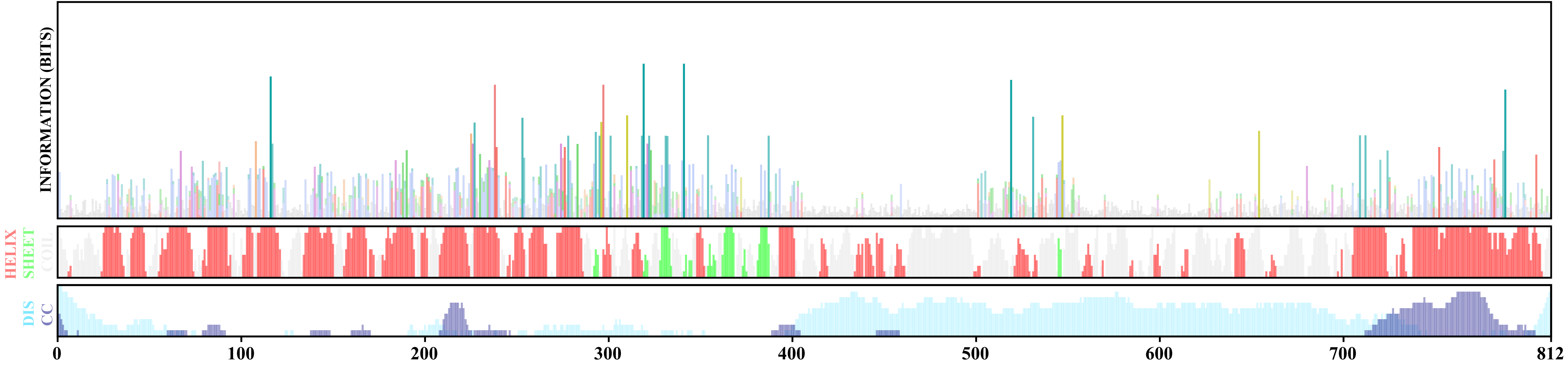

### SYCP3_Diplonemids.pdf

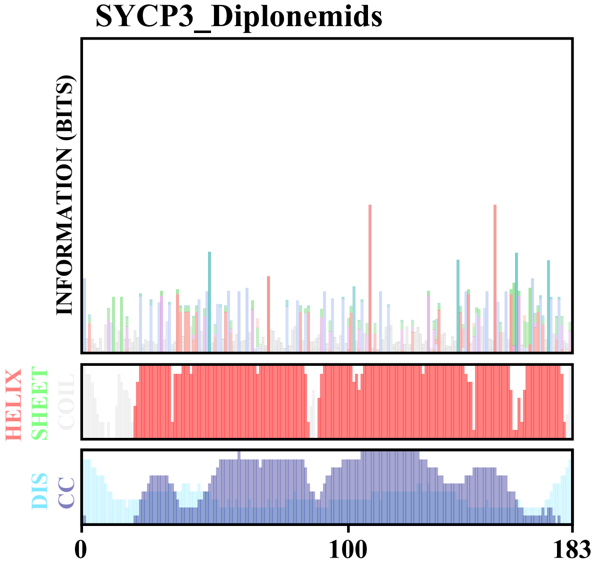

### SYCP3_Euglenids.pdf

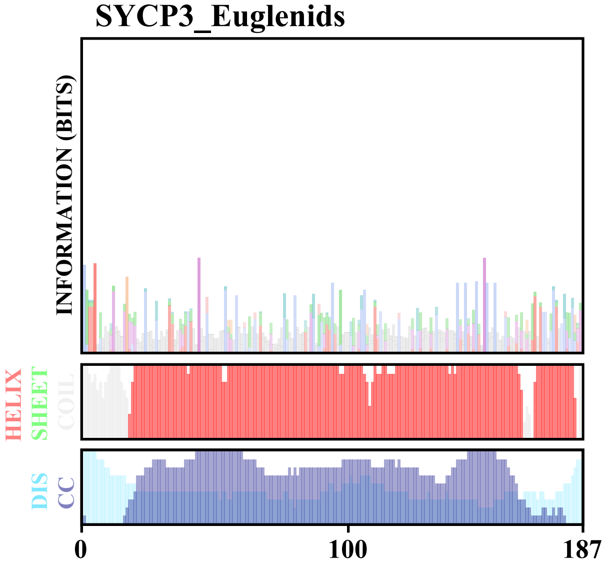

### SYCP3_Metazoa.pdf

SYCP3_Metazoa

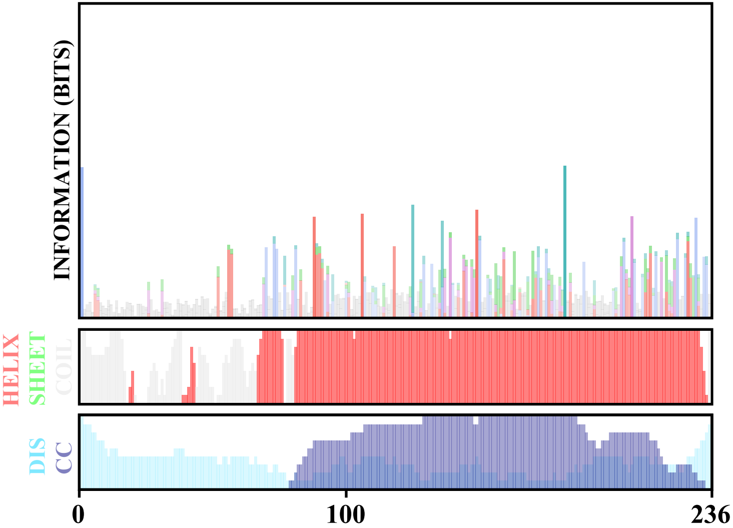
