## Supplementary material for "Repurposing of Synaptonemal Complex Proteins for Kinetochores in Kinetoplastida": File S6: make_fasta_sequence_single_lined.pl

```
#!/usr/bin/perl 

use strict;
use warnings;

#fasta_file input
foreach (@ARGV) {
					#declare
					my $prot_id;
     				my %id2seq;
     				my $id;
					my $fasta_file = $_;
		
					#open and obtain data
					open(FASTA_FILE, "<$fasta_file");
					while (<FASTA_FILE>) {
						chomp;
						if (/^\>/) 				{$prot_id = $_, $id2seq{$prot_id} ="", print "$prot_id\n"}
						elsif (!/^\>/)	 		{$id2seq{$prot_id} .= $_}
						else 					{print "something went wrong\n"}
					}
	
					close FASTA_FILE;
	
					#print all IDs + fasta sequence
					open (OUTPUT_ALL, ">$fasta_file");
					foreach my $key (keys %id2seq) 		{print OUTPUT_ALL "$key\n" , "$id2seq{$key}\n";}
					close OUTPUT_ALL;				
}				
					
exit;
```
