## Supplementary material for "Repurposing of Synaptonemal Complex Proteins for Kinetochores in Kinetoplastida": File S6: OutputFigure.py

```
#!/usr/bin/env python

import math
import cairo
import numpy as np
import sys
import json
import ParsedHMM
import colorsys
import collections
from scipy.stats import rankdata

class OutputFigure:
    """ Set of utilities for producing output figure

    Uses Cairo to generate a PDF figure with the name specified in the configuration.

    """

    cr = None
    padding_left = 0
    padding_top = 0
    config = None
    parsed_hmm_master = None

    def __init__(self, config):
        # Load the HMM
        self.parsed_hmm_master = ParsedHMM.ParsedHMM(config)
        # Store the config and HMM
        self.config = config
        # Set up the page
        self.padding_left = config['page']['padding_left']
        self.padding_top = config['page']['padding_top']
        surface = cairo.PDFSurface(config['output']['file_name'], self.parsed_hmm_master.length + config['page']['horizontal_padding'], config['page']['height'])
        self.cr = cairo.Context(surface)
        # Draw the master sequence
        self.draw_master_sequence()
        # Save the file
        self.save_file()
        
      
    def plot_clustal(self, scale, max_bitscore, position, offset_horizontal, draw_full_rectangle, hmm):
        # Plot text area
        self.cr.set_font_size(8.5)
        self.cr.select_font_face("Times New Roman", cairo.FONT_SLANT_NORMAL, cairo.FONT_WEIGHT_BOLD)
        (x, y, width, height, dx, dy) = self.cr.text_extents("INFORMATION (BITS)")
        self.cr.move_to(offset_horizontal - height / 2, position-(1.5*scale) + width + ((scale * max_bitscore) - width)/2)
        self.cr.rotate(-math.pi / 2)
        self.cr.show_text("INFORMATION (BITS)")
        self.cr.rotate(math.pi / 2)
                
        # Plot the clustal plot on top
        for i in range(len(hmm.clustal_colours)):
            height_offset = 0
            for j in range(len(self.config['colours'])):
                # Calculate colour
                if self.config['output']['conservation_plot']['type'] == 'skylign':
                    if np.sum(hmm.clustal_colours[i]) > 0.5:
                        col = [self.config['colours'][j]['rgb'][0], self.config['colours'][j]['rgb'][1],self.config['colours'][j]['rgb'][2], self.config['colours'][j]['rgb'][3]]
                        col[3] = (np.divide(np.sum(hmm.clustal_colours[i])*2.75, (np.max(hmm.height_array))))
                    else:
                        col = [0.5, 0.5, 0.5, 0.15]
                
                else:
                    if np.sum(hmm.clustal_colours[i]) > 1.5:
                        col = [self.config['colours'][j]['rgb'][0], self.config['colours'][j]['rgb'][1],self.config['colours'][j]['rgb'][2], self.config['colours'][j]['rgb'][3]]
                        col[3] = (np.divide(np.sum(hmm.clustal_colours[i]), (np.max(hmm.height_array))))
                    else:
                        col = [0, 0, 0.8]
                
                self.cr.set_source_rgba(col[0], col[1], col[2], col[3])

                height = scale * hmm.clustal_colours[i][j]
                if height>(max_bitscore *scale):
                    height = max_bitscore*scale
                self.cr.rectangle(offset_horizontal + 0.5 + i, position - height + (scale * 4) + height_offset, 1,height)
                height_offset -= height
                self.cr.fill()

        #plot box        
        self.cr.set_source_rgb(0, 0, 0)
        self.cr.set_line_width(1)
        self.cr.rectangle(self.padding_left, position-(1.5*scale), self.parsed_hmm_master.length + 1, scale * max_bitscore)
        self.cr.stroke()
        self.cr.close_path()

    def draw_master_sequence(self):
        self.plot_clustal((30/7)*5, 5.5, self.padding_top, self.padding_left, self.config['output']['split'],self.parsed_hmm_master)
        if self.config['output']['subplot_type'] == "secondary":
            self.draw_ss(self.parsed_hmm_master, self.padding_top + 225-65, self.padding_left, -1, self.parsed_hmm_master.length)
        elif self.config['output']['subplot_type'] == "psiplot":
            self.bar_ss(self.parsed_hmm_master, self.padding_top + 225-65, self.padding_left, -1, self.parsed_hmm_master.length)
            self.bar_disorder(self.parsed_hmm_master, self.padding_top + 257-65, self.padding_left, -1, self.parsed_hmm_master.length) 
            self.bar_cc(self.parsed_hmm_master, self.padding_top + 257-65, self.padding_left, -1, self.parsed_hmm_master.length)
                      
        
        # Draw a box to represent the main sequence
        draw_masters_at = [self.padding_top]
        if self.config['output']['split']:
            draw_masters_at.append(self.padding_top)

        for vertical_pos_index in range(len(draw_masters_at)):
            vertical_pos = draw_masters_at[vertical_pos_index]
            above_or_below = np.remainder(vertical_pos_index, 2) * 1

            # Draw box for protein - not in use anymore - but other parts are dependent on this segment so left in - needs to be cleaned up
            height_rectangle = 20
            #self.cr.set_source_rgb(0, 0, 0)
            #self.cr.set_line_width(1)
           # self.cr.rectangle(self.padding_left, vertical_pos + 219, self.parsed_hmm_master.length + 1, height_rectangle)
           # self.cr.stroke()
           # self.cr.close_path()
            self.cr.set_font_size(12)
            self.cr.set_source_rgb(0, 0, 0)
            self.cr.select_font_face("Times New Roman", cairo.FONT_SLANT_NORMAL, cairo.FONT_WEIGHT_BOLD)
            (x, y, width, height, dx, dy) = self.cr.text_extents(self.parsed_hmm_master.name)
            self.cr.move_to(self.padding_left + 5, vertical_pos - (1.5*((30/7)*5)) + height - 20 + ((height_rectangle - height)/2))
            self.cr.show_text(str(self.parsed_hmm_master.name))

            # Add on some residue numbering beneath the box
            self.cr.set_font_size(10)
            for i in range(0, self.parsed_hmm_master.length - 50, 100):
                (x, y, width, height, dx, dy) = self.cr.text_extents(str(i))
                self.cr.move_to(self.padding_left - 0.75 + i - width / 2,
                                vertical_pos + 219-65 + 5 + height / 2 + above_or_below * (180 - 151 + 40))
                self.cr.show_text(str(i))
                self.cr.move_to(self.padding_left + i, vertical_pos + 210-65 + 5+ above_or_below * (180 - 151 + 40))
                self.cr.line_to(self.padding_left + i, vertical_pos + 210-65 + 5 + 3 + above_or_below * (180 - 151 + 40))

            (x, y, width, height, dx, dy) = self.cr.text_extents(str(self.parsed_hmm_master.length))
            self.cr.move_to(self.padding_left - 0.75 + 1 + self.parsed_hmm_master.length - width / 2,
                            vertical_pos + 219-65 + 5 + height / 2 + above_or_below * (180 - 151 + 40))
            self.cr.show_text(str(self.parsed_hmm_master.length))
            self.cr.move_to(self.padding_left + 1 + self.parsed_hmm_master.length,
                            vertical_pos + 210-65 + 5 + above_or_below * (180 - 151 + 40))
            self.cr.line_to(self.padding_left + 1 + self.parsed_hmm_master.length,
                            vertical_pos + 210-65 + 5 + 3 + above_or_below * (180 - 151 + 40))
            self.cr.stroke()

            
    def bar_ss (self,hmm,pos_y, pos_x, hit_start, hit_end, right_cutff=100000):
        
        max_bitscore = 9
        offset_horizontal = pos_x
        position = pos_y
        scale = 3
        
        self.cr.set_source_rgba(0, 0, 0, 1)
        self.cr.set_line_width(1)
        self.cr.rectangle(self.padding_left, position - 70 , self.parsed_hmm_master.length + 1, scale * 9 + 1)
        self.cr.stroke()
        self.cr.close_path()
        
        # text
        self.cr.set_font_size(8.5)
        self.cr.set_source_rgba(1, 0, 0, 0.5)
        self.cr.select_font_face("Times New Roman", cairo.FONT_SLANT_NORMAL, cairo.FONT_WEIGHT_BOLD)
        self.cr.close_path()
        self.cr.set_font_size(8.5)
        
        (x, y, width, height, dx, dy) = self.cr.text_extents("HELIX")
        self.cr.move_to(offset_horizontal - height / 2 - 20, position - 41 + (scale * max_bitscore - width) / 2)
        self.cr.rotate(-math.pi / 2)
        self.cr.show_text("HELIX")
        self.cr.rotate(math.pi / 2)
        
        self.cr.set_font_size(8.5)
        self.cr.set_source_rgba(0, 1, 0, 0.5)
        self.cr.select_font_face("Times New Roman", cairo.FONT_SLANT_NORMAL, cairo.FONT_WEIGHT_BOLD)
        self.cr.close_path()
        self.cr.set_font_size(8.5)
        
        (x, y, width, height, dx, dy) = self.cr.text_extents("SHEET")
        self.cr.move_to(offset_horizontal - height / 2 - 10, position - 41 + (scale * max_bitscore - width) / 2)
        self.cr.rotate(-math.pi / 2)
        self.cr.show_text("SHEET")
        self.cr.rotate(math.pi / 2)
        
        self.cr.set_font_size(8.5)
        self.cr.set_source_rgba(0.9, 0.9, 0.9, 0.6)
        self.cr.select_font_face("Times New Roman", cairo.FONT_SLANT_NORMAL, cairo.FONT_WEIGHT_BOLD)
        self.cr.close_path()
        self.cr.set_font_size(8.5)
        
        (x, y, width, height, dx, dy) = self.cr.text_extents("COIL")
        self.cr.move_to(offset_horizontal - height / 2, position - 48 + (scale * max_bitscore - width) / 2)
        self.cr.rotate(-math.pi / 2)
        self.cr.show_text("COIL")
        self.cr.rotate(math.pi / 2)
        
        # Plot the clustal plot on top
        for i in range(len(hmm.probs)):
            height_offset = -69.5

            #helix SS is in red. The sheet SS is in green. The coil SS is in gray
            col = [0.9, 0.9, 0.9, 1]
            if hmm.ss[i] == "H":
                col = [1, 0, 0, 0.5]
            elif hmm.ss[i] == "E":
                col = [0, 1, 0, 0.5]
            elif hmm.ss[i] == "C":
                col = [0.9, 0.9, 0.9, 0.5]
                #col[3] = np.square(int(hmm.ss_probs[i]))/np.square(max_bitscore)
            
            # Height is the ss_prob
            height = int(hmm.ss_probs[i]) * scale
            #height = max_bitscore * scale
            

            self.cr.set_source_rgba(col[0], col[1], col[2], col[3])

            self.cr.rectangle(offset_horizontal + 0.5 + i, position - height + scale * max_bitscore + height_offset, 1,
                              height)
            height_offset -= height
            self.cr.fill()
    
    
    
    def bar_disorder(self, hmm, pos_y, pos_x, hit_start, hit_end, right_cutoff=100000):
        # (self, scale, max_bitscore, position, offset_horizontal, draw_full_rectangle, hmm):
        # H is green, E is yellow, C is grey
        max_bitscore = 9
        offset_horizontal = pos_x
        position = pos_y
        scale = 3
        
        self.cr.set_source_rgba(0, 0, 0, 1)
        self.cr.set_line_width(1)
        self.cr.rectangle(self.padding_left, position - 70 , self.parsed_hmm_master.length + 1, scale * 9 + 1)
        self.cr.stroke()
        self.cr.close_path()
        
        # Draw axes
        self.cr.set_source_rgba(0, 0.82, 1, 0.5)
        self.cr.set_font_size(8.5)
        self.cr.select_font_face("Times New Roman", cairo.FONT_SLANT_NORMAL, cairo.FONT_WEIGHT_BOLD)
        self.cr.close_path()
        self.cr.set_font_size(8.5)
        (x, y, width, height, dx, dy) = self.cr.text_extents("DIS")
        self.cr.move_to(offset_horizontal - height/2 - 10, position - 54 + (scale * max_bitscore - width) / 2)
        self.cr.rotate(-math.pi / 2)
        self.cr.show_text("DIS")
        self.cr.rotate(math.pi / 2)

        #print(hmm.ss_probs)

        # Plot on top
        for i in range(len(hmm.probs)):
            height_offset = -69.5

            #colour disorder positions
            col = [0, 0.82, 1, 0.15]
            if hmm.disorder[i] == "D":
                col = [0, 0.82, 1, 0.15]
                #col[3] = np.square(int(hmm.disorder_probs[i]))/np.square(max_bitscore)
            elif hmm.disorder[i] == "O":
                col = [0, 0.82, 1, 0.15]
                #col[3] = np.square(int(hmm.disorder_probs[i]))/np.square(max_bitscore)
                
            # Height is the ss_prob
            height = int(hmm.disorder_probs[i]) * scale
            #height = max_bitscore * scale
            

            self.cr.set_source_rgba(col[0], col[1], col[2], col[3])

            self.cr.rectangle(offset_horizontal + 0.5 + i, position - height + scale * max_bitscore + height_offset, 1,
                              height)
            height_offset -= height
            self.cr.fill()
    
    def bar_cc (self,hmm,pos_y, pos_x, hit_start, hit_end, right_cutff=100000):
        
        max_bitscore = 9
        offset_horizontal = pos_x
        position = pos_y
        scale = 3
        
        self.cr.set_source_rgba(0, 0, 0, 1)
        self.cr.set_line_width(1)
        self.cr.rectangle(self.padding_left, position - 70 , self.parsed_hmm_master.length + 1, scale * 9 + 1)
        self.cr.stroke()
        self.cr.close_path()
        
        # Draw axes
        self.cr.set_source_rgba(0, 0, 0.5, 0.5)
        self.cr.set_font_size(8.5)
        self.cr.select_font_face("Times New Roman", cairo.FONT_SLANT_NORMAL, cairo.FONT_WEIGHT_BOLD)
        self.cr.close_path()
        self.cr.set_font_size(8.5)
        (x, y, width, height, dx, dy) = self.cr.text_extents("CC")
        self.cr.move_to(offset_horizontal - height / 2, position - 55.5 + (scale * max_bitscore - width) / 2)
        self.cr.rotate(-math.pi / 2)
        self.cr.show_text("CC")
        self.cr.rotate(math.pi / 2)

        # Plot on top
        for i in range(len(hmm.probs)):
            height_offset = -69.5

            #F is coiled-coil, N is not a coiiled-coil
            col = [0, 0, 0.5, 0.35]
            if hmm.cc[i] == "F":
                col = [0, 0, 0.5, 0.35]
                #col[3] = np.square(int(hmm.cc_probs[i]))/np.square(max_bitscore)
            elif hmm.cc[i] == "N":
                col = [0, 0, 0.5, 0.35]
                #col[3] = np.square(int(hmm.cc_probs[i]))/np.square(max_bitscore)

            # Height is the cc_prob
            height = int(hmm.cc_probs[i]) * scale
            #height = max_bitscore * scale
            self.cr.set_source_rgba(col[0], col[1], col[2], col[3])
            self.cr.rectangle(offset_horizontal + 0.5 + i, position - height + scale * max_bitscore + height_offset, 1,
                              height)
            height_offset -= height
            self.cr.fill()
        
    

    def save_file(self):
        self.cr.save()
        self.cr.show_page()
```
