## Supplementary material for "Repurposing of Synaptonemal Complex Proteins for Kinetochores in Kinetoplastida": File S6: parse_IUPred_and_Deepcoil.pl

```
#!/usr/bin/perl 
use strict;
use warnings;
use POSIX;

#declare
my ($iupred_file) 	= $ARGV[0];
my ($deepcoil)		= $ARGV[1];
my ($file_name)		= $ARGV[2];

my %pos2seq;
my %iupred_data;
my @order;
my @deepcoil_order;

#parse IUPRED_file
open(IUPRED, "<$file_name/$file_name.iupred");
while (<IUPRED>) {	
	chomp;
	if (/^#/)			{next}
	elsif (/^[0-9]/)		{	my @a=split(/\t/,$_);
		 						my $id = $a[1].$a[0];
								$iupred_data{$id}=$a[2];
								push (@order, $id);
							}	
	else				{next}				
}
close IUPRED;

#parse Deepcoil_order
open(DEEPCOIL, "<$file_name/deepcoil_order");
while (<DEEPCOIL>) {	
	chomp;
	push @deepcoil_order, $_;
	next;
}
close DEEPCOIL;

#parse_Deepcoil_files
my $iteration = 0;
my $counter = 0;
my $protein_length = scalar @order;
print "$protein_length\n";
my $boundary_1=0;
my $boundary_2=0;
my $total=0;

#due to length limit of 500 in Deepcoil1.1 we used a moving window of 500 aa with 150 aa overlap  to predict coiled coils, the new version of Deepcoil: 2 has no limit so this part will be obsolete
for (@deepcoil_order) {
	$iteration+=1;
	print "$_\n";
	open (DEEP_COILS, "<$file_name/$_") or die "Something is going wrong\n";
	while (<DEEP_COILS>) {
		chomp;
		if ($iteration==1){
			my $start = 0;
			if (/^[A-Z]/){
				$counter++;
				if ($counter<401){
					my $total = $counter + $start;
					my @b = split (/\s/,$_);
					my $id = $b[0].$total;
					print "$id $b[2]\n";
					push (@{$pos2seq{$id}},{iupred=>$iupred_data{$id}, deep_coil=>$b[2]});
				}
				else {next}
			}
			else {next}
		}		
		elsif ($iteration>1){
			if (/^[A-Z]/){
				$counter++;
				$boundary_1 = $iteration*350 + 50 - 350;
				$boundary_2 = 401 + $iteration*350 - 350;
				$total = $counter - (150*$iteration-150);
				if ((401 + $iteration*350-350+100)>$protein_length) {$boundary_2 = $protein_length+1}
				if ($total>$boundary_1 && $total<$boundary_2){	
					my @b = split (/\s/,$_);
					my $id = $b[0].$total;
					print "$id $b[2]\n";
					push (@{$pos2seq{$id}},{iupred=>$iupred_data{$id}, deep_coil=>$b[2]});
				}
				else {next}
			}
			else {next}
		}
		
	}
	close (DEEP_COILS);

}
# print to file
open (OUTPUT, ">$file_name/deepcoil.iupred.output");
my $iupred_score;
my $iupred_state;
my $deep_coil_score;
my $deep_coil_state;
my $disordered_state = 'D';
my $ordered_state = 'O';
my $coiled_state = 'F';
my $non_coiled_state = 'N';
foreach my $positions (@order) {
	foreach my $residues (@{$pos2seq{$positions}}) {
		$iupred_score .= floor(10*$residues->{iupred});
		if (floor(10*$residues->{iupred})>4) 	{$iupred_state .=$disordered_state}
		else 					{$iupred_state .=$ordered_state}
		$deep_coil_score.= floor(10*$residues->{deep_coil});
		if (floor(10*$residues->{deep_coil})>4) {$deep_coil_state .=$coiled_state}
		else 					{$deep_coil_state .=$non_coiled_state}
	}
}
print OUTPUT "\>IUPRED_score\n";
for my $buffer ($iupred_score =~ m/.{1,100}/gs) {
	print OUTPUT $buffer, "\n";
}
print OUTPUT "\>IUPRED_state\n";
for my $buffer ($iupred_state =~ m/.{1,100}/gs) {
	print OUTPUT $buffer, "\n";
}  	
print OUTPUT "\>DEEPCOIL_score\n";
for my $buffer ($deep_coil_score =~ m/.{1,100}/gs) {
	print OUTPUT $buffer, "\n";
}
print OUTPUT "\>DEEPCOIL_state\n";
for my $buffer ($deep_coil_state =~ m/.{1,100}/gs) {
	print OUTPUT $buffer, "\n";
}  
close (OUTPUT);


exit;
```
