## Supplementary material for "Repurposing of Synaptonemal Complex Proteins for Kinetochores in Kinetoplastida": File S6: parse_MSA_cut.pl

```
#!/usr/bin/perl 
use strict;
use warnings;

#declare
my ($input_file) = $ARGV[0];
my ($name) = $ARGV[1];
my ($base_name) = $ARGV[2];
my $first_ID ='none';
my $name_2;
my $name_3;
my $name_4;	
my %id2seq;
my $id;

#parse database
open(INPUT_FILE, "<$input_file");
while (<INPUT_FILE>) {	
	chomp;
	if (/^\>/ && $first_ID eq 'none')	{$id = $_, $id2seq{$id} ="", $first_ID = $_}
	elsif (/^\>/)						{$id = $_, $id2seq{$id} =""}
	elsif (/^[A-Za-z0-9\-\*]+$/)		{$_ =~ s/X/-/g,$id2seq{$id} .= $_}				
}
close INPUT_FILE;

#parse names
open(NAME, "<$name");
while (<NAME>) {	
	chomp;
	if (/^\>/) {$name_3 = $_}
	else {next}
}
close NAME;


#parse numbers and print files
my @a = split (/\//,$name_3);
$name_2 = substr($a[0], 1, length($a[0]));
$name_4 = $name_2;
print "$name_4\n";
$name_4 =~ s/[^a-zA-Z0-9\_]//g;
print "$name_4\n";

my $increment = 350;
my $length = 500;
my $end = 150;
if ($end>length($id2seq{$first_ID})) {$end=0}

open (PRINT_OUT_2, ">","$base_name/deepcoil_order") or die "Couldn't open: $!";

for (my $pos = -1; $end<length($id2seq{$id}); $pos+= $increment) { 
	$end+=$increment;
	print "$end\n";
	my $sequence='';
	my$pos1 = $pos + 2;
	my$pos2;
	if ($pos<0) {$pos2 = 0}
	else {$pos2 = $pos+1}
	if ($end>length($id2seq{$id})) {$end = length($id2seq{$id})}
	print PRINT_OUT_2 "$name_4"."$pos1"."$end"."\.out","\n";
	open (PRINT_OUT,">","$base_name/$name_4$pos1$end.cut") or die "Couldn't open: $!";
	print PRINT_OUT "\>$name_2\/$pos1\-$end\n";
	print PRINT_OUT substr($id2seq{$name_3}, $pos2, $length), "\n";
	foreach my $prot_id (keys %id2seq)	{
		if ($prot_id eq $name_3) {next}
		else {
		my @b = split (/\//,$prot_id);
		my $corrected_protein_id = $b[0];
		$sequence = substr($id2seq{$prot_id}, $pos2, $length);
		if ($sequence =~ /[A-Z]/)	{		
			print PRINT_OUT "$corrected_protein_id\n";
			print PRINT_OUT $sequence, "\n";			
		}
		}
	}
	close (PRINT_OUT);

}
close (PRINT_OUT_2);

exit;
```
