## Supplementary material for "Repurposing of Synaptonemal Complex Proteins for Kinetochores in Kinetoplastida": File S6: ParsedHMM.py

```
#!/usr/bin/env python

import math
import requests
import numpy as np
import json

class ParsedHMM:
    """ Represents a parsed HMM file

    This stores representations of HHSuite HMMs.
    """

    def __init__(self, config):
        """ Parse HMM file given in configuration, and load into memory

        :param config: parsed JSON configuration object
        """
        # Define the important parameters as state variable
        self.hmm_string = ''
        self.alphabet = []
        self.states = []
        self.probs = []
        self.state_probs = []
        self.clustal_colours = []
        self.height_array = []
        self.ss = ''
        self.ss_probs = ''
        self.cc = ''
        self.cc_probs = ''
        self.disorder = ''
        self.disorder_probs = ''
        self.nulls = []

        # Process HMM
        print(config['master']['hmm_file'])
        with open(config['master']['hmm_file']) as hmm:
            in_hmm_section = False
            in_psipred_ss = False
            in_psipred_ss_probs = False
            in_psipred_cc = False
            in_psipred_cc_probs = False
            in_psipred_disorder = False
            in_psipred_disorder_probs = False
            
            for line in hmm:
                if line.startswith("LENG"):
                    self.length = int(line.split()[1])
                if line.startswith("NAME"):
                    if ".fa" in line or "_" in line:
                        self.name = config['master']['name']
                    else:
                        self.name = line.split()[1]
                if line.startswith("FILT"):
                    self.num_seqs = float(line.split()[1])
                if line.startswith("HMM"):
                    in_hmm_section = True
                if in_hmm_section:
                    self.hmm_string += line

                # Process null
                if line.startswith("NULL"):
                    splitLine = line.split()
                    for i in range(1, len(splitLine)):
                        self.nulls.append(2 ** (int(splitLine[i]) / -1000))

                # Clear current section if we're at a new FASTA header
                if line.startswith(">"):
                    in_psipred_ss = False
                    in_psipred_ss_probs = False
                    in_psipred_cc = False
                    in_psipred_cc_probs = False
                    in_psipred_disorder = False
                    in_psipred_disorder_probs = False

                # NB, unlike for HMM, we don't want the ss to include the FASTA header, so we check section before
                if in_psipred_ss:
                    self.ss += line
                if "ss_pred PSIPRED predicted secondary structure" in line:
                    in_psipred_ss = True
                    
                if in_psipred_ss_probs:
                    self.ss_probs += line
                if "ss_conf PSIPRED confidence values" in line:
                    in_psipred_ss_probs = True
                    
                if in_psipred_cc:
                    self.cc += line
                if "DEEPCOIL_state" in line:
                    in_psipred_cc = True
                    
                if in_psipred_cc_probs:
                    self.cc_probs += line
                if "DEEPCOIL_score" in line:
                    in_psipred_cc_probs = True
                    
                if in_psipred_disorder:
                    self.disorder += line
                if "IUPRED_state" in line:
                    in_psipred_disorder = True
                    
                if in_psipred_disorder_probs:
                    self.disorder_probs += line
                if "IUPRED_score" in line:
                    in_psipred_disorder_probs = True
                    

        # Remove linebreaks/whitespace from ss and ss_probs
        self.ss = ''.join(self.ss.split())
        self.ss_probs = ''.join(self.ss_probs.split())
        self.cc = ''.join(self.cc.split())
        self.cc_probs = ''.join(self.cc_probs.split())
        self.disorder = ''.join(self.disorder.split())
        self.disorder_probs = ''.join(self.disorder_probs.split())
        

        # Now process the HMM string we have
        hmm_lines = self.hmm_string.split("\n")
        for idx, line in enumerate(hmm_lines):
            # Extract the fields within the HMM file
            if line.startswith("HMM"):
                for hmm_line in range(len(line.split())):
                    if hmm_line != 0:
                        self.alphabet.append(line.split()[hmm_line])
                for hmm_line in range(len(hmm_lines[idx + 1].split())):
                    self.states.append(hmm_lines[idx + 1].split()[hmm_line])

        # Now move on to the match states themselves
        for l in range(self.length):
            alphabet_line = hmm_lines[l * 3 + 3].split()
            stats_line = hmm_lines[l * 3 + 4].split()
            current_alphabet_probs = []
            for a in range(len(alphabet_line)):
                if a > 1 and a < len(alphabet_line) - 1:
                    if alphabet_line[a] is '*':
                        f = 0
                    else:
                        f = 2 ** (int(alphabet_line[a]) / -1000)
                    current_alphabet_probs.append(f)
            self.probs.append(current_alphabet_probs)
            current_states = []
            for s in range(len(stats_line)):
                if stats_line[s] is '*':
                    f = 0
                else:
                    f = 2 ** (int(stats_line[s]) / -1000)
                current_states.append(f)
            self.state_probs.append(current_states)


        if 'output' not in config or config['output']['conservation_plot']['type'] == 'traditional':
            # Go through each position and calculate the height
            for height_idx in range(len(self.probs)):
                entropy = self.get_shannon_entropy(height_idx)

                height = entropy
                self.height_array.append(height)
            for aa_idx in range(len(self.probs)):
                height = self.height_array[aa_idx]
                heights = [0, 0, 0, 0, 0, 0, 0, 0]
                for j in range(len(self.probs[aa_idx])):
                    aa_prob = self.probs[aa_idx][j]
                    aa_name = self.alphabet[j]
                    for k in range(len(config['colours'])):
                        if aa_name in config['colours'][k]['aa']:
                            if aa_prob > self.nulls[j]:
                                heights[k] += aa_prob * height
                self.clustal_colours.append(heights)
        elif config['output']['conservation_plot']['type'] == 'skylign':
            # Get the HMM file and send it to skylign via post
            print("Making request")
            r = requests.post('http://skylign.org/',
                              files = {
                                  'file': ('hmm.a3m', open(config['master']['alignment_a3m_original'], 'rb')),
                                  'processing': (None, 'hmm_all'),
                                  'path': (None, '/'),
                                  'letter_height': (None, 'info_content_above')
                              },
                              headers={"Accept": "application/json"}, timeout=5)
            # Now retrieve the heights from skylign
            parsed_response = json.loads(r.text)
            print (parsed_response)
            print(parsed_response['message'])
            r = requests.get(parsed_response['url'],
                             headers={"Accept": "application/json"}, timeout=5)
            skylign_model = json.loads(r.text)
            #print (r.text)
            for position in skylign_model['height_arr']:
                # Sum all the heights
                height = 0
                for aa in position:
                    height += float(aa.split(":")[1])
                self.height_array.append(height)
                # Now get the heights for each AA group
                heights = [0, 0, 0, 0, 0, 0, 0, 0]
                for aa in position:
                    aa_prob = float(aa.split(":")[1])
                    aa_name = aa.split(":")[0]
                    for k in range(len(config['colours'])):
                        if aa_name in config['colours'][k]['aa']:
                            heights[k] += aa_prob
                self.clustal_colours.append(heights)

        return

    def getKullbackLeiblerDistance(self, position):
        return

    def get_shannon_entropy(self, height_idx):
        position = self.probs[height_idx]
        shannon_entropy = 0
        for i in range(len(position)):
            if position[i] > 0:
                if position[i] > self.nulls[i]:
                    shannon_entropy += position[i] * math.log2(position[i] / self.nulls[i])
        return shannon_entropy

    def get_small_sample_correction(self):
        return 1 / math.log(2) * (20 - 1) / (2 * self.num_seqs)
```
