## Supplementary material for "Repurposing of Synaptonemal Complex Proteins for Kinetochores in Kinetoplastida": File S6: simple-figgen.py

```
#!/usr/bin/env python

import sys
import json
import ParsedHMM
import OutputFigure

#################################################### CONFIGURATION ####################################################
# Edit this section to set up the script
name = "file_name"

# Path to the HHsuite HHM file (with ss info)
hhm_file = "file_name/file_name.iupred.deepcoil.ss.a3m.hhm"

# A user-friendly display name of HHM
hhm_name = "file_name"

# Path to the a3m file (with ss info)
a3m_file = "file_name/file_name.ss.a3m"
a3m_file_original = "file_name/file_name.a3m"

# Output file name
output_file = "file_name/file_name.pdf"

# Page configuration
page_config = {
    "horizontal_padding": 80,
    "height": 240,
    "padding_left": 40,
    "padding_top": 62.5
  }

################################################## End of config#######################################################
# Set a config
config = {
  "master": {
    "name": hhm_name,
    "hmm_file": hhm_file,
    "alignment_a3m": a3m_file,
    "alignment_a3m_original": a3m_file_original,
  },
  "page": page_config,
  "output": {
    "file_name": output_file,
    "max_hits": 10,
    "split": False,
    "split_at": 500,
    "subplot_type": "psiplot", #psiplot, logo, secondary
    "conservation_plot": {
      "type": "skylign", #traditional, skylign
      "args": {
        "letter_height": "info_content_above"
      }
    }
  },
  "colours": [
    {
      "name": "blue",
      "aa": "A,I,L,M,V",
      "rgb": [0.5020, 0.6275, 0.9412, 1]
    },
    {
      "name": "red",
      "aa": "K,R",
      "rgb": [0.9412, 0.0824, 0.0196, 1]
    },
    {
      "name": "magenta",
      "aa": "E,D",
      "rgb": [0.7529, 0.2824, 0.7529, 1]
    },
    {
      "name": "green",
      "aa": "N,Q,S,T",
      "rgb": [ 0.0824, 0.7529, 0.0824, 1]
    },
    {
      "name": "pink",
      "aa": "C",
      "rgb": [ 0.9412, 0.5020, 0.5020, 1]
    },
    {
      "name": "orange",
      "aa": "G",
      "rgb": [0.9412, 0.5647, 0.2824, 1]
    },
    {
      "name": "yellow",
      "aa": "P",
      "rgb": [0.7529, 0.7529, 0, 1]
    },
    {
      "name": "cyan",
      "aa": "H,Y,F,W",
      "rgb": [0.0824, 0.6431, 0.6431, 1]
    }
  ]
}

# Generate the output figure
output_figure = OutputFigure.OutputFigure(config)
```
