## Supplementary material for "Repurposing of Synaptonemal Complex Proteins for Kinetochores in Kinetoplastida": File S6: 4_manual_on_how_to_generate_annotated_HHM_based_on_fasta_alignment.txt

#this is an overview of the bash commands that are used to generate PDF files of annotated HHM.

##present in directory from which to execute script (make sure only these files are present)
	#(1) alignment_file in fasta format with extension .fasta
	#(2) simple-figgen.py
	#(3) ParseHMM.py
	#(4) OutputFigure.py
	#(5) perl scrips: make_fasta_single_lined; parse_MSA_cut; parse_IUPred_and_Deepcoil

#standard environment
	#python 3.5
	#legacy_blast 2.2.26 (32-bit so no good on MacOS Catalina and newer versions)
	#HHsuite3
	#psipred 4.02
	#IUpred2a

#environment for Deepcoil (setup using for instance conda)
	#python 3.6
	#NCBI_BLAST+

#assuming you have these safety measures in place - don't forget to put these back after running the script
alias rm="rm -i";
alias cp="cp -i";

#unalias safety measure to run script
unalias rm;
unalias cp;

#declare - change here if you want
extension=fasta;
#notes on fasta files:
	#(1) no spaces in file name
	#(2) make sure each sequence has a unique ID
	#(3) remove X's for unknown amino acids
	#(4) need more than one sequence

#the following loops are needed to run to generate files - each loop can in principle be copy pasted to the command line - most should work in a standard linux/MacOS environment. To test I suggest you run loop by loop manually and later collate them into one script that works for you.

#loop 1: make directories and files
for file in *.$extension; do name=$(basename $file .$extension); 
	rm -r $name;
	mkdir $name; 
	cp simple-figgen.py $name/;
	cp OutputFigure.py $name/;  
	cp ParsedHMM.py $name/;
	#preparing the simple-figgen.py file for generating all annotation files
	perl -pi -w -e 's/file\_name/"'$name'"/g' $name/simple-figgen.py; 
done;	

#loop 2: run HHsuite over alignment to generate HHM file
for file in *.$extension; do name=$(basename $file .$extension); 
	# 32 bit old version of psipred script/legacy blast needed  - put HHsuite/psipred scripts on your path
	export PATH=$PATH:/directory/to/HHsuite/scripts/
	# for HHsuite/psipred you need to have legacy blast 2.2.26 installed
	reformat.pl fas a3m $name.$extension $name/$name.a3m -g '-' -r; 
	addss.pl $name/$name.a3m $name/$name.ss.a3m;
	hhmake -i $name/$name.ss.a3m -o $name/$name.ss.a3m.hhm; 
done;
	
#loop 3: run IUPRED 
for file in *.$extension; do name=$(basename $file .$extension); 
	perl -00 -ne '/(>[^>]+)/ && print $1' $name/$name.a3m > $name/$name.reference;
	perl make_fasta_sequence_single_lined.pl $name/$name.reference;
	python3.6 /Users/ecet2/Software/iupred2a/iupred2a.py $name/$name.reference short > $name/$name.iupred; 
done;

#loop 4: run Deepcoil 1.1 (moving window of 500 with 150 overlap)
for file in *.$extension; do name=$(basename $file .$extension); 
	#some neccessary cutting and pasting
	head -n 1 $name/$name.reference > $name/$name.reference.ID;
	tail -n 1 $name/$name.reference > $name/$name.reference.sequence;
	perl parse_MSA_cut.pl $name/$name.a3m $name/$name.reference.ID $name;
	
	for files in $name/*.cut; do names=$(basename $name/$files .cut); 
		perl -00 -ne '/(>[^>]+)/ && print $1' $name/$names.cut > $name/$names.cut.reference;
		psiblast -subject $name/$names.cut.reference -in_msa $files -out_ascii_pssm $name/$names.pssm -ignore_msa_master;
		#generate a conda environment named deepcoil --> needs python3.5
		conda activate deepcoil; 
		python3.5 /Users/ecet2/Software/Deepcoil/deepcoil.py -i $name/$names.cut.reference -pssm -pssm_path $name/;
		conda deactivate;
		mv *.out $name/;
	done;
done;

#loop 5: parse IUPred/Deepcoil files and generate final annotated HHM
for file in *.$extension; do name=$(basename $file .$extension); 
	#parse IUPred and Deepcoil files
	perl parse_IUPred_and_Deepcoil.pl $name/$name.iupred $name/deepcoil_order $name;
	
	#make final annotated HHM file
	perl -e 'while(<>){print;if($_=~/^SEQ/){system("cat '$name'/deepcoil.iupred.output")}}' $name/$name.ss.a3m.hhm > $name/$name.iupred.deepcoil.ss.a3m.hhm;
done;

#loop 6: generate PDF files
for file in *.$extension; do name=$(basename $file .$extension); 
	python3 $name/simple-figgen.py;
done;
